# Tropical-temperate dichotomy falls apart in the Asian Palmate Group of Araliaceae

**DOI:** 10.1101/2021.10.20.465102

**Authors:** Marina Coca-de-la-Iglesia, Nagore G. Medina, Jun Wen, Virginia Valcárcel

## Abstract

**PREMISE:** The use of climatic data on phylogenetic studies has greatly increased in the last decades. High-quality spatial data and accurate climatic information are essential to minimize errors in the climatic reconstructions to the past. However, despite the huge amount of already available biodiversity digital information, the process of compiling, cleaning, and comparing spatial data from different open data sources is a time-consuming task that sometimes ends up with low-quality geographical information. For this reason, researchers often resort qualitative approximations among which World bioclimatic classification systems or the experts’ criteria are the most used. Our aim is to evaluate the climatic characterization of the genera of the Asian Palmate Group (AsPG) of the ginseng family (Araliaceae), one of the classical examples of tropical-temperate plant families.

**METHODS:** We compiled a curated worldwide spatial database of the AsPG genera. We then created five raster layers representing bioclimatic regionalizations of the World. Finally, we crossed the database with the layers to characterize the AsPG genera.

**RESULTS:** We found large disagreement in the climatic characterization of genera among regionalizations and little support for the tropical-temperate dichotomy. Both results are attributed to the complexity of delimiting tropical, subtropical and temperate climates in the World and to the distribution of the study group in regions with transitional climatic conditions.

**CONCLUSIONS:** The complexity in the climatic classification of this classical example tropical-temperate dichotomy, calls for a general revision in other families. In fact, we claim that to properly evaluate tropical-temperate transitions we cannot ignore the complexity of distribution ranges.

## INTRODUCTION

The use of climatic data on phylogenetic studies has become routine since it often provides insightful information to better understand the evolutionary history of lineages (see for example, Pyron and Wiens, 2013; Edwards et al., 2017; Nürk et al., 2018; Albaladejo et al., 2021). To do so, not only robust phylogenies are needed, but also large amounts of good quality climatic data (Budic and Dormann, 2015). However, to obtain accurate climatic data and conduct robust phylogenetic reconstructions good geographical information of the study group distribution is advocated as the starting input data (Hortal et al., 2007). Because of this, the compilation of high-quality spatial databases is a cornerstone in these integrative approaches.

The huge amount of biodiversity data that have been collected by naturalists and scientists during the last centuries has been gathered together in online services in last decades and it is now generally available with just one click (GBIF; GBIF.org, 2021, iNaturalist; iNaturalist, 2021, TRY; Fraser, 2020, Specieslink SpeciesLink, 2021). However, the process from the initial downloading-click to the final compilation of high-quality spatial databases is often long and complex. First, it is often necessary to handle large volumes of data that require intense data cleaning (data parsing and homogenization) before they are ready for analyses. Dealing with typos and other frequent errors such as, wrong geo-references, incorrect taxonomical identifications of records or outdated nomenclature, is a very time consuming task of the data cleaning (Soberón and Peterson, 2004). This is so even in databases like GBIF that contain categories to facilitate the cleaning of invalid georeferenced records (Yesson et al., 2007). Data homogenization is also a time-consuming step of the data cleaning that may also become a serious limitation when different online data sources are combined (Turnhout and Boonman-Berson, 2011). Once the cleaning is done, an evaluation of the representativeness and accuracy of the spatial database compiled is needed before performing any further analysis. This is an essential step since online databases do suffer from the fact that our knowledge is incomplete and/or unevenly distributed across the World and the tree of life (Hortal et al., 2015; Meyer et al., 2016). For example, well-developed regions, legally protected areas or temperate deciduous woodlands have traditionally been over sampled (Martin et al., 2012). This sampling bias can be explained by different factors that have nothing to do with biodiversity itself but have great impact on data gathering, such as the economic wealth, modern language, geographical location and security of the region (Amano and Sutherland, 2013). Therefore, to take full advantage of the enormous potential of online spatial databases we also need to evaluate our gaps of biodiversity knowledge and consider them in our conclusions (Hortal et al., 2008; Boakes et al., 2010).

As a result, even after a good data cleaning the spatial database compiled may end up revealing a poor representativeness of the distribution of the study group, and so it should not be used to describe its climatic niche. Because of this, qualitative approaches are often used to characterize the climatic preferences of taxa in evolutionary reconstructions. For example, very frequently taxa are classified according to the climatic categories of a given climatic classification system (i.e., Köppen & Geiger, 1936) or according to the taxonomist’s expert criteria and this categorization is used as the climatic input data for the phylogenetic reconstructions (see for example Edwards et al., 2017; Silva et al., 2021). However, the particular climatic classification used for the categorization is rarely stated (Feeley and Stroud, 2018), and if so, the procedure to include taxa in one or another category (based on distribution maps, expert criterion, etc.), is often neglected in the section of materials and methods.

Araliaceae is one of the classical examples of tropical-temperate plant families (Judd, 1994). The largest clade of Araliaceae is the Asian Palmate Group (hereafter AsPG (Plunkett et al., 1996, 2004; Plunkett and Lowry, 2001; Wen et al., 2001) that includes almost half of the genera (23 genera and c. 950 spp.) in the family and all its North temperate genera (Wen et al., 2001). The large distribution of the AsPG extends through Southeast Asia, Europe, North Africa, the Americas and North Oceania. Most of the genera are endemic to Southeast Asia (13), four also extend to other continents (*Hedera* L. in Europe, *Dendropanax* Decne. & Planch. and *Oplopanax* (Torr. & A.Gray) Miq. in the Americas, and *Heptapleurum* Gaertn in Oceania), and the remaining six are endemic to Central and/or South America. Across the worldwide distribution of the clade, genera also tend to be widespread, as 17 of them expand over large areas (including intercontinental and transoceanic distributions) and only a few occur in restricted regions (Frodin and Govaerts, 2003; Fang et al., 2011). According to experts’ criteria (Wen et al., 2001), the clade is mainly tropical, since 16 genera (70%) occur in the tropics or subtropics while only seven in temperate zones. Interestingly, a seemingly easiness to shift niches is inferred from molecular phylogenies (Wen et al., 2001; Valcárcel and Wen, 2019), since temperate genera occur scattered across the AsPG tree which points to independent acquisition of temperate affinity. However, no explicit evaluation of the climatic characterization of the AsPG genera has been done so far.

Our main objective is to provide the characterization of the climatic preferences of the 23 AsPG genera by using explicit semi-quantitative approach. To do so, we used online biodiversity repositories and compiled a worldwide georeferenced database of the AsPG to extract qualitative climatic data for all the records. To extract qualitative climatic data we created two raster layers and modified three layers already available to represent the climatic regionalization of the World according to five bioclimatic classifications frequently used in evolutionary studies. The specific objectives are to: (1) compile a worldwide high-quality point-occurrence database that provide an accurate representativeness of the geographical distribution of the AsPG genera, (2) compile a high-quality climatic database of the AsPG genera, (3) characterize the geographical range and climatic preferences of the AsPG genera, and (4) evaluate the impact of using different climatic classifications on the climatic characterization of the AsPG.

## MATERIALS AND METHODS

### Spatial characterization of the AsPG

#### Spatial data collection

Eight online databases were used to download the spatial records of the AsPG from March 2018 to April 2020 (Table 1). Downloads were done either through the website (Neotropical Plant Portal: https://serv.biokic.asu.edu/neotrop/plantae/, Neotropical Plant Portal, 2021; NBN: nbnatlas.org/, NBN Atlas, 2021; Tropicos: www.tropicos.org/, Tropicos.org, 2021; TRY: www.try-db.org/ and WPKorea: florakorea.myspecies.info/, Chang and Kim, 2015) or using the available R packages (“rgibf” from GBIF Chamberlain et al., 2020a; DOI references for the original downloads are available in Zenodo from Coca-de-la-Iglesia et al., 2021a, “BIEN” from BIEN (Maitner et al., 2018) and “spocc” from iNaturalist (Chamberlain et al., 2020b). Since our target is to provide a geographical and climatic characterization at genus level, searches for downloads were done by genus instead of species, except for six genera (*Cephalopanax* G.M. Plunkett, Lowry & D.A.Neill, *Crepinella* Marchal, *Didymopanax* Decne. & Planch., *Frodinia* Lowry & G.M. Plunkett, *Heptapleurum*, and *Sciadophyllum* P. Browne). These six genera have been recently recognized because of a major taxonomical rearrangement of two independent lineages of the former genus *Schefflera* J. R. Forst. & G. Forst. that were included within the AsPG (Lowry II et al., 2019; Fiaschi et al., 2020; Lowry II et al., 2020; Lowry II and Plunkett, 2020; Plunkett et al., 2021). Because of the polyphyly of the former *Schefflera*, the searches for these six genera were done by species’ names instead of genus. Also, because the taxonomical rearrangements were published in parallel to our downloading process and nomenclature was not updated on online databases, these searches were done under the respective *Schefflera* synonym. To identify which of the former *Schefflera* species belong to the Neotropical and Asian *Schefflera* clades of the AsPG, we matched the species distribution of all *Schefflera* species, as recorded in the World Check list of Araliaceae (Frodin and Govaerts, 2003), with the geographical ranges of the main linages of the former *Schefflera* described in Plunkett et al. (2005) and later checked with the synonyms included in the taxonomical rearrangements’.

**Table 1.** Summary of the compilation and cleaning information of spatial database of the Asian Palmate Group of Araliaceae (AsPG).

| Original source | Records before<br>cleaning | Rercords after<br>cleaning | % of records |
| --- | --- | --- | --- |
| **Online databases** | **679,857** | **471,811** | **99.89** |
| GBIF<sup>1</sup> | 633,315 | 448,846 | 94.17 |
| BIEN | 30,558 | 167,93 | 3.52 |
| NBN | 8,871 | 4,870 | 1.02 |
| iNaturalist | 6,212 | 3,739 | 0.78 |
| Neotropical Plant Portal | 1,560 | 1,119 | 0.23 |
| Tropicos | 690 | 643 | 0.13 |
| WPKorea | 143 | 132 | 0.03 |
| TRY | 68 | 52 | 0.01 |
| **Herbaria<sup>2</sup>** | **426** | **379** | **0.08** |
| AVH | 11 | 7 | - |
| CVH | 134 | 111 | - |
| E | 79 | 68 | - |
| US | 121 | 120 | - |
| MO | 58 | 52 | - |
| NYBG | 22 | 20 | - |
| K | 1 | 1 | - |
| **Literature<sup>2</sup>** | **145** | **131** | **0.03** |

1. DOI references for the original downloads are available in Zenodo (Coca-de-la-Iglesia et al., 2021a).
2. Records not uploaded in any of the online databases analyzed.

To increase the number of records for poorly represented genera at a global or regional scale in the online databases used, a targeted search on geo-referenced specimens was done on eight herbaria (Table 1). Gaps in distribution knowledge persisted for certain genera and/or regions and were fulfilled by targeted searches of localities in systematic studies (*Trevesia* Vis., *Hedera, Macropanax* Miq., *Sciodaphyllum, Gamblea* C.B.Clarke and *Heteropanax* Seem.; Jebb, 1998; Shang et al., 2000; Jamir and Pandey, 2003; Heriyanto and Sawitri, 2007; Prabhu et al., 2010; Tagane et al., 2017; Jiménez-Montoya and Idárraga-Piedrahíta, 2018; Ong, 2018; Amini et al., 2020). Because most of this localities lacked coordinates, a georeferencing process was done using GeoLocate Web Application (Rios and Bart, 2010). A position point was marked in zones with the highest probability of appearance of the specie, according to the description of the locality provided in each record. Google Earth and Google Street View were also used to help precision.

Finally, we extracted the altitude for all records to homogenized or fill the gaps in altitudinal information. To do so, we grouped the 60 altitudinal files available in WorldClim (30 secs, Fick and Hijmans, 2017) to form a new layer named “Elevation_WorldClim_30sec” that can be obtained from Zenodo (Coca-de-la-Iglesia et al., 2021b).

#### Data cleaning

To clean the spatial data compiled we developed a script in R program (R Core Team, 2018) that reduces the timing by automating the process (Coca-de-la-Iglesia et al. in prep.) This step of the process was designed to address two main objectives: (1) homogenize data and remove duplicates and (2) reduce the effect of spatial uncertainty and non-natural records on the climatic characterization of genera further conducted with the spatial database.

For the first objective, we homogenized the administrative information across all records according to two types of standardized country codes and using vector layers in QGIS 3.4.3-Madeira (QGIS Development Team, 2021). We used the coordinates of records to extract the countries for the third level of TDWG geographical information (Brummitt, 2001) from the layer available in GitHub repository (Desmet and Page, 2007) and the 2-letter ISO-3166-1 code for each record as obtained from Admin-0 Countries layer of Natural Earth 4.1.0 (Patterson and Kelso, 2020). This procedure also allowed us to correct country typos in the original data sources. Then, we proceeded removing the duplicates originated by the compilation of different online data sources that sometimes share records. We identified duplicates as those that have the same information for eight fields of the database (species name: “Spp”; collection year: “Year”; country code: “CountryCode”; locality: “Locality”; longitude: “Longitude”; latitude: “Latitude”; elevation; “Elevation”; Specimen voucher or herbaria number: “catalogNumber”; and type of record: “basisOfRecord”). Once identified, only one of the duplicated records was kept and the remaining were removed.

For the second objective of the data cleaning, we first removed all records that contained spatial uncertainty. This includes records with none, zero or less than two decimals in their coordinates, and those with erroneous coordinates. We defined a threshold of 10 km distance from the coastal line to identify erroneous records. Because of the variability in the land surface limits of the different layers, we used one of the 19 Worldclim bioclimatic layers (bio1: Annual Mean Temperature; Fick and Hijmans, 2017; https://www.worldclim.org/) to establish the limits of land surface. Using this layer as template, we removed all records at >10 km distance from coastal limit. We decided to keep records within the 10 km coastal distance buffer (hereafter “coastal line records”) to avoid the loss of seaboard environments that might be potentially informative for the climatic characterization of genera. To keep these coastal line records, their original coordinates were recalculated to meet the nearest climatic cell of the template.

Finally, to avoid nuisance in the climatic inferences coming from the inclusion of cultivars and records outside the natural ranges of genera we identified and removed all non-natural wild records. We identified cultivated records as those in which the locality description included any of the following words: “cultivated”, “cultivado”, “park”, “parque”, “garden”, “jardín”, “castel”, “castillo”, “golf”, “cementerio”, “zoo”, “farm”. Each of the records identified as cultivated was manually checked before removal. To remove non-native records, we first created a vector to represent the natural range of each genus using the botanical countries of the TDWG geographical standard (Brummitt, 2001) as the spatial unit. To do so, we used the third level of TDWG code of the botanical countries of the genus native range as in the World Checklist of Selected Plant Families (WCSP, Govaerts et al., 2008). These vectors were crossed with the country code field that included the third level of TDWG code in the database (“CountryCode_TDWG”) to remove records from non-matching countries (i.e, records outside native range).

#### Spatial data analyses

To represent the native distribution of the AsPG in the World, a point-occurrence map was elaborated including all records of the cleaned database. Toevaluate the spatial biodiversity trends at a global scale we built a bubble map by using the botanical countries of the clade as the spatial units and the number of genera per spatial unit as the value to estimate the size of bubbles. Then, we built heat maps as a proxy to evaluate sampling effort in the AsPG across the World. Because global patterns in sampling effort may hinder other patterns at a finer scale, we decided to build three heat maps: Asia (including Oceania), Europe (including North Africa) and the Americas. To build each heat map, we used a cell area of one geographical degree as the spatial unit and the number of records per spatial unit to estimate the sampling effort in each cell. To categorize cells according to the number of records within a gradient, Jenks natural breaks were used. To identify sampling hotspots in each region (areas including cells with high levels of sampling effort), we first estimated the minimum and maximum sampling effort in each cell category for each heat map. To do so, we compared the minimum and maximum number of records of each cell category in each region with the maximum number of records per cell detected in that given region. As a result, we calculated two sampling effort values for each cell category in each region that represent the range of sampling effort. Cell categories including 25% of sampling effort within their ranges were considered as sampling hotspots. Finally, to evaluate temporal sampling patterns across the World in the AsPG we plotted the accumulated number of occurrences in time per region with “ggplot2” package (Wickham et al., 2021) in R version 3.5 (R Core Team, 2018). We used the three same regions as for the heat maps. Due to the limited volume of data for years prior to 1900 (1,479 records), occurrences recorded before that date were not used for the temporal series.

To represent the native ranges of the AsPG genera, 23 point maps were elaborated (one per genus). Also, to evaluate the sampling effort per genus, 23 World heat maps were built following the same procedure as above described for the clade. Finally, to identify restricted vs. widespread genera we estimated the area of occupancy (AOO) and the extent of occurrence (EOO) in km^2^ for each genus using a distance of 2 km as the buffer to estimate the AOO, as recommended by International Union for Conservation of Nature (IUCN, 2012). These estimates were done in GeoCAT (Bachman et al., 2011) for all genera except for two (*Dendropanax* and *Hedera*). Because of the large disjunction of *Dendropanax* and the great number of *Hedera* occurrences their AOO and EOO were estimated in QGIS. In these cases, we calculated the total area of convex hull created around all occurrences as the estimate for the EOO. Finally, to identify genera as restricted, we applied the minimum threshold of the IUCN for the Vulnerable threatened category at global scale (IUCN, 2012). We considered this proxy as highly conservative to identify restricted genera, since the IUCN categories are intended for species classification and we are applying this threshold at the genus level.

All maps were elaborated with QGIS version 3.4.3-Madeira (QGIS Development Team, 2021) using the shapefile that includes the third level of TDWG code of botanical countries.

### Climatic characterization of the AsPG

#### Climatic data collection and layer compilation

Climatic data were obtained for all the records in the AsPG database using two different approaches: qualitative and semiquantitative. For the qualitative approach (hereafter “expert criterion”), all records from each genus were classified as tropical or temperate according to the climatic preference as stated by the taxonomists of the family (Plunkett et al., 1996; Wen et al., 2001).

For the semiquantitative approach, the records of the AsPG database were crossed with five spatial layers representing five bioclimatic regionalizations of the World from analytical and synthetical classification systems that are frequently used in evolutionary studies. The selected five World bioclimatic classification systems are: (1) The Latitudinal zonation, that divides the World in four zones solely based on latitude; (2) Köppen’s classification (Köppen and Geiger, 1936) that recognizes six zones mainly based on temperature and precipitation; (3) Holdridge’s classification (Holdridge, 1967) that identifies seven world life zones based on biotemperature (BioT), which is the temperature at which plants grow efficiently (between 0° C and 30° C); (4) Metzger’s classification (Metzger et al., 2012) that established a Global Environmental Stratification System with seven broad biomes based on 42 bioclimatic variables; and (5) Ecoregions’ system (Olson et al., 2001) that divides the World in 14 biomes based on environmental conditions and the biogeographical information of the World’s floras and faunas. To make the genera’s climatic characterizations comparable between the most analytical classification systems (Metzger’s classification and Ecoregions’ system) and the most synthetic ones (Latitudinal zonation, Köppen’s and Holdridge’s classifications), we selected the broadest hierarchical category of each classification systems as our analysis scale. To cross our database with each classification system, we needed to adapt three geospatial layers already available (Ecoregion’s: Dinerstein et al., 2017; Köppen’s: Beck et al., 2018; Meztger’s: Metzger et al., 2012) and create two new layers for the remaining classification systems (Latitudinal zonation and Holdridge’s classification).

To adapt Köppen’s layer, we used the improved version to 1 km resolution of traditional classification of Köppen-Geiger developed by Beck et al., 2018. We grouped the second and third level of climatic categories to only consider the main classes: Tropical (A), Dry (B), Temperate (C), Continental (D), Tundra (E) and Polar (F) (available in GitHub: https://github.com/vvalnun/Bioclimatic-classifications-AsPG.git; Coca-de-la-Iglesia et al., 2021c). For Metzger’s classification we used the layer including the global environmental zones provided in Metzger et al. (2012) to obtain the seven broad biomes and transformed the resulting layer to tif format (available in GitHub: https://github.com/vvalnun/Bioclimatic-classifications-AsPG.git; Coca-de-la-Iglesia et al., 2021c). In the case of the Ecoregions system, we used the biomes delimitation of the shapefile available in Dinerstein et al. (2017). Then, we simplified this classification in R (hereafter “Simplified Ecoregions”) considering two new inclusive categories that gathered together some of the original categories recognized (Dinerstein et al., 2017). As a result, we created the “Tropical and subtropical” category that unified four original biomes (“Tropical and subtropical dry broadleaf forests”, “Tropical and subtropical moist broadleaf forests”, “Tropical and subtropical coniferous forests” and “Tropical and subtropical grasslands, savannas, and shrublands”) and the “Temperate” category that included three original biomes (“Temperate broadleaf and mixed forests”, “Temperate conifer forests”, and “Temperate grasslands, savannas, and shrublands”) and transformed the resulting layer to tif format (available in GitHub: https://github.com/vvalnun/Bioclimatic-classifications-AsPG.git; Coca-de-la-Iglesia et al., 2021c).

The new layer created to represent the Latitudinal zonation was built by setting 23.5° as the geographical limit between tropical and subtropical zones, 40° for subtropical and temperate and 66.5° for temperate and polar, both in northern and southern hemispheres (available in GitHub: https://github.com/vvalnun/Bioclimatic-classifications-AsPG.git; Coca-de-la-Iglesia et al., 2021c). To build the new layer for Holdridge’s classification we estimated BioT from the 12 layers including the average monthly temperature from 1970 to 2000 at a 30 secs resolution available in WorldClim (Fick and Hijmans, 2017; https://www.worldclim.org/). First, we reclassified each layer to represent BioT by setting to 0 all temperatures below 0° C and above 30° C and keeping the original temperature value between 0° C and 30° C. With the 12 reclassified monthly layers we created a new layer that contained the mean of BioT. Finally, we classified BioT values from the mean BioT layer according to the latitudinal World Life zones recognized by Holdridge (Holdridge, 1967): Tropical (30-24° BioT), Subtropical (24-17° BioT), Warm Temperate (17-12° BioT), Cool Temperate (12-6° BioT), Boreal (6-3° BioT), Sub-polar (3,5-1,5° BioT) and Polar (1,5-0° BioT) available in GitHub: https://github.com/vvalnun/Bioclimatic-classifications-AsPG.git; Coca-de-la-Iglesia et al., 2021c).

Once the five layers were prepared, they were crossed with the spatial database of the AsPG to assign each record of the database to the corresponding climatic category according to each classification system.

#### Climatic characterization of genera

We provide the characterization of the climatic preferences of the AsPG at two levels (genus and clade) and according to the six classification systems. For the five classification systems analyzed with the semi-quantitative approach, we computed the percentage of records classified within any given category. To avoid overestimations due to taxonomical sampling bias, percentages for the AsPG climatic characterization at the clade level were done using a reduced database. In this reduced database all oversampled genera (>1,500 records) were each represented by 1,000 records regularly selected with the function “spsample” of the package “sp” (Pebesma et al., 2021) in R version 3.5 (R Core Team, 2018). Pie charts were then performed from this reduced database for each bioclimatic classification with “ggplot2” package (Wickham et al., 2021) in R version 3.5 (R Core Team, 2018). To characterize the climatic preference of each genus, we estimated the percentage of occurrences classified in each category per genus per classification system. To avoid overestimations due to geographical sampling bias, percentages were estimated by only retaining one record per coordinate per genus.

## RESULTS

### Spatial representativeness of the AsPG database

The database represents 100% of the AsPG genera, including the most recent taxonomical rearrangements of the former genus *Schefflera*. It contains 476,704 records resulting from a cleaning process of an initial database of 683,207 observations and covers the worldwide distribution range of the clade (Coca-de-la-Iglesia et al., 2021a; Fig. 1). The data cleaning process resulted in losses from 5% (*Metapanax*) to 71% (*Cephalopanax*) of records per genus (Table 2). *Hedera* is the genus with the highest number of occurrences (401,252; Table 2) followed by *Oreopanax* with 20,108 (4%; Table 2), and *Dendropanax* with 13,600 records (3%; Table 2). The genera with the least number of records are *Cephalopanax* (28 occurrences; Table 2) and *Frodinia* (48 occurrences; Table 2). The genus with the highest sampling effort is *Hedera* with a maximum of 8,093 records per cell while the genera with the lowest sampling effort are *Cephalopanax* and *Heteropanax* with a maximum of six records per cell in both cases (Appendix S1; see Supporting Information with this article). Genera point maps encompass the distribution range of each genus (Appendix S2) and revealed that in 12 genera all the botanical countries of their native range are represented while the sampling incompleteness of the remaining 11 varies between 3% in *Didymopanax* and 50% in *Metapanax* where half of the botanical countries of the natural range are missing.

**Table 2.** Spatial information of the genera of the Asian Palmate Group (AsPG) of Araliaceae. NSpp: Number of accepted species for each genus. Range: Distribution range. Ndow: Number of records downloaded. Nred: Number of records kept after the data cleaning. Loss: Percentage of data loss during the data cleaning. RepGen: Percentage of records per genus in the AsPG database. RepSpp: Percentage of species recorded per genus in the AsPG database. NBcountry: Number of botanical countries of the natural range of each genus. SamInc: Sampling incompleteness, estimated as the percentage of botanical countries of the natural distribution of the each genus not represented in the AsPG database. AOO: Area of occupancy in km^2^. EOO: Extent of occurrence in km^2^.

| Genus | N<sup>Spp</sup> | Range | N<sup>dow</sup> | N<sup>red</sup> | Loss<br>(%) | Rep<sup>Gen</sup><br>(%) | Rep<sup>Spp</sup><br>(%) | N<sup>Bcountry</sup> | SamInc<br>(%) | AOO | EOO |
| --- | --- | --- | --- | --- | --- | --- | --- | --- | --- | --- | --- |
| *Brassaiopsis* | 48 | Asia | 1062 | 953 | 10,26 | 0,20 | 64,58 | 21 | 9,52 | 1108 | 9,85E+06 |
| *Cephalopanax* | 3 | South America | 98 | 28 | 71,43 | 0,01 | 100,00 | 2 | 0,00 | 88 | 1,74E+05 |
| *Chengiopanax* | 2 | Asia | 1178 | 1072 | 9,00 | 0,22 | 100,00 | 3 | 0,00 | 2864 | 3,77E+06 |
| *Crepinella* | 33 | South America | 1115 | 788 | 29,33 | 0,17 | 96,97 | 7 | 0,00 | 1292 | 5,71E+06 |
| *Dendropanax* | 103 | Asia, Central &<br>South America | 21914 | 13600 | 37,94 | 2,85 | 91,26 | 51 | 7,84 | 22132 | 2,32E+08 |
| *Didymopanax* | 37 | Central & South<br>America | 10483 | 8788 | 16,17 | 1,84 | 100,00 | 32 | 3,13 | 17552 | 2,11E+07 |
| *Eleutherococcus* | 38 | Asia | 5704 | 4704 | 17,53 | 0,99 | 76,32 | 21 | 4,76 | 6796 | 1,35E+07 |
| *Fatsia* | 3 | Asia | 2826 | 877 | 68,97 | 0,18 | 66,67 | 6 | 0,00 | 2068 | 1,28E+06 |
| *Frodinia* | 2 | Central America | 54 | 48 | 11,11 | 0,01 | 100,00 | 3 | 0,00 | 92 | 4,50E+04 |
| *Gamblea* | 4 | Asia | 1112 | 955 | 14,12 | 0,20 | 100,00 | 13 | 23,08 | 2188 | 1,06E+07 |
| *Hedera* | 12 | Europe, North<br>Africa & Asia | 576291 | 401252 | 30,37 | 84,17 | 100,00 | 68 | 7,35 | 190564 | 4,95+07 |
| *Heptapleurum* | 317 | Asia & North<br>Oceania | 8201 | 6371 | 22,31 | 1,34 | 58,68 | 34 | 8,82 | 9632 | 4,03E+07 |
| *Heteropanax* | 10 | Asia | 228 | 168 | 26,32 | 0,04 | 70,00 | 15 | 33,33 | 328 | 3,22E+06 |
| *Kalopanax* | 1 | Asia | 2117 | 1611 | 23,90 | 0,34 | 100,00 | 10 | 0,00 | 3808 | 6,30E+06 |
| *Macropanax* | 17 | Asia | 471 | 392 | 16,77 | 0,08 | 70,59 | 17 | 23,53 | 908 | 1,02E+07 |
| *Merrilliopanax* | 3 | Asia | 170 | 158 | 7,06 | 0,03 | 100,00 | 6 | 33,33 | 184 | 3,02E+05 |
| *Metapanax* | 2 | Asia | 938 | 894 | 4,69 | 0,19 | 100,00 | 6 | 50,00 | 568 | 1,20E+06 |
| *Oplopanax* | 3 | Asia & North<br>America | 4289 | 3941 | 8,11 | 0,83 | 100,00 | 14 | 0,00 | 6200 | 1,29E+07 |
| *Oreopanax* | 148 | Central & South<br>America | 30168 | 20108 | 33,35 | 4,22 | 97,30 | 35 | 2,86 | 18980 | 2,58E+07 |
| *Sciodaphyllum* | 138 | Central & South<br>America | 12614 | 8874 | 29,65 | 1,86 | 92,75 | 14 | 0,00 | 8512 | 8,32E+06 |
| *Sinopanax* | 1 | Asia | 119 | 111 | 6,72 | 0,02 | 100,00 | 1 | 0,00 | 288 | 1,30E+04 |
| *Tetrapanax* | 1 | Asia | 1676 | 716 | 57,28 | 0,15 | 100,00 | 4 | 0,00 | 880 | 2,26E+06 |
| *Trevesia* | 11 | Asia & North<br>Oceania | 367 | 293 | 20,16 | 0,06 | 90,91 | 17 | 23,53 | 688 | 9,30E+06 |

**Figure 1.**
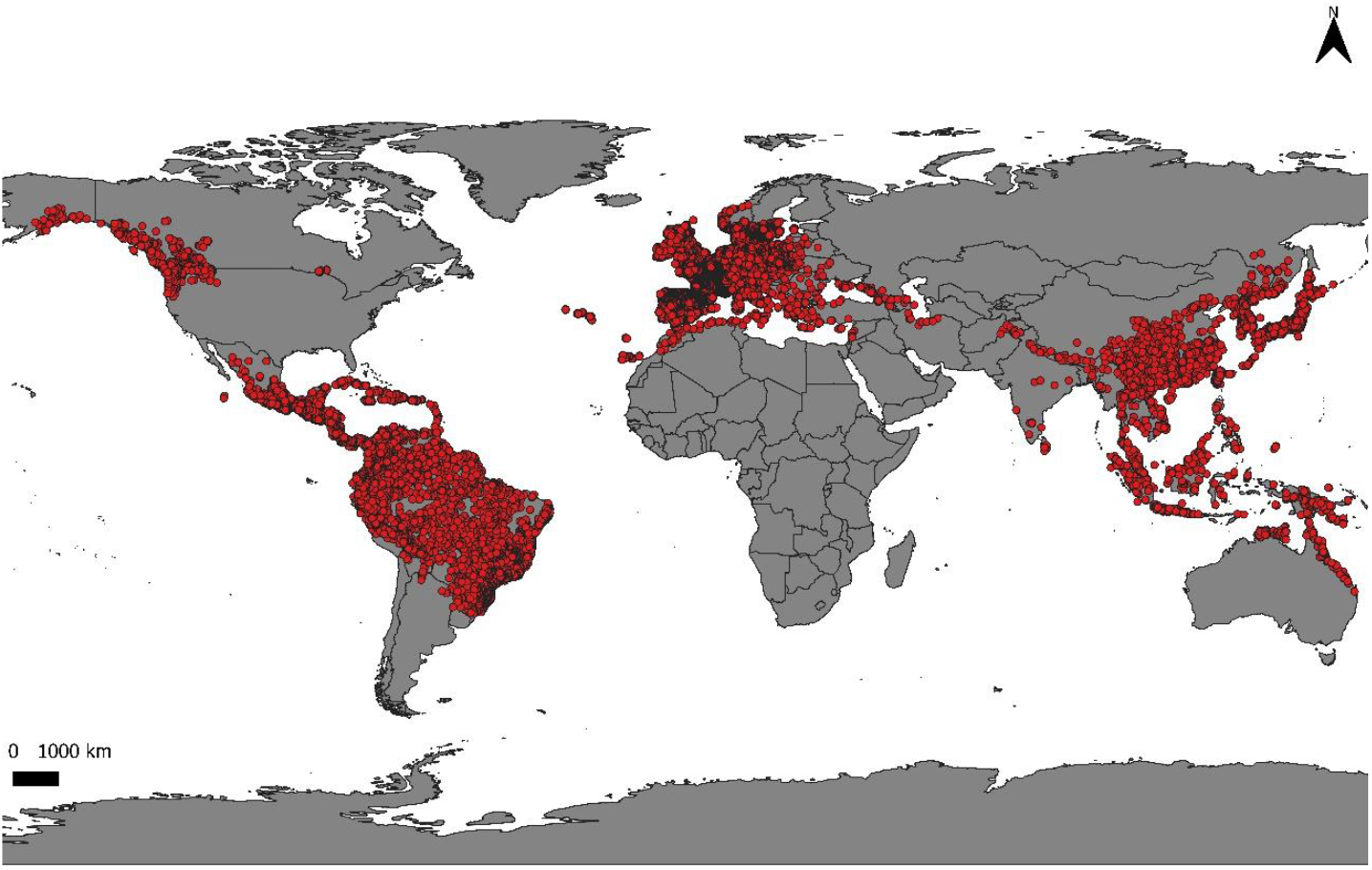
Global point-occurrence map representing the geographical distribution of Asian Palmate Group of Araliaceae.

### Spatial patterns in the AsPG

The analysis of spatial diversity in the AsPG reveals that the number of genera is unevenly distributed across the World (Fig. 2) with Southeast Asia as the area with the highest number of genera per botanical country, followed by Central and South America. Europe and North America have the lowest diversity with only one genus per botanical country (Fig. 2).

**Figure 2.**
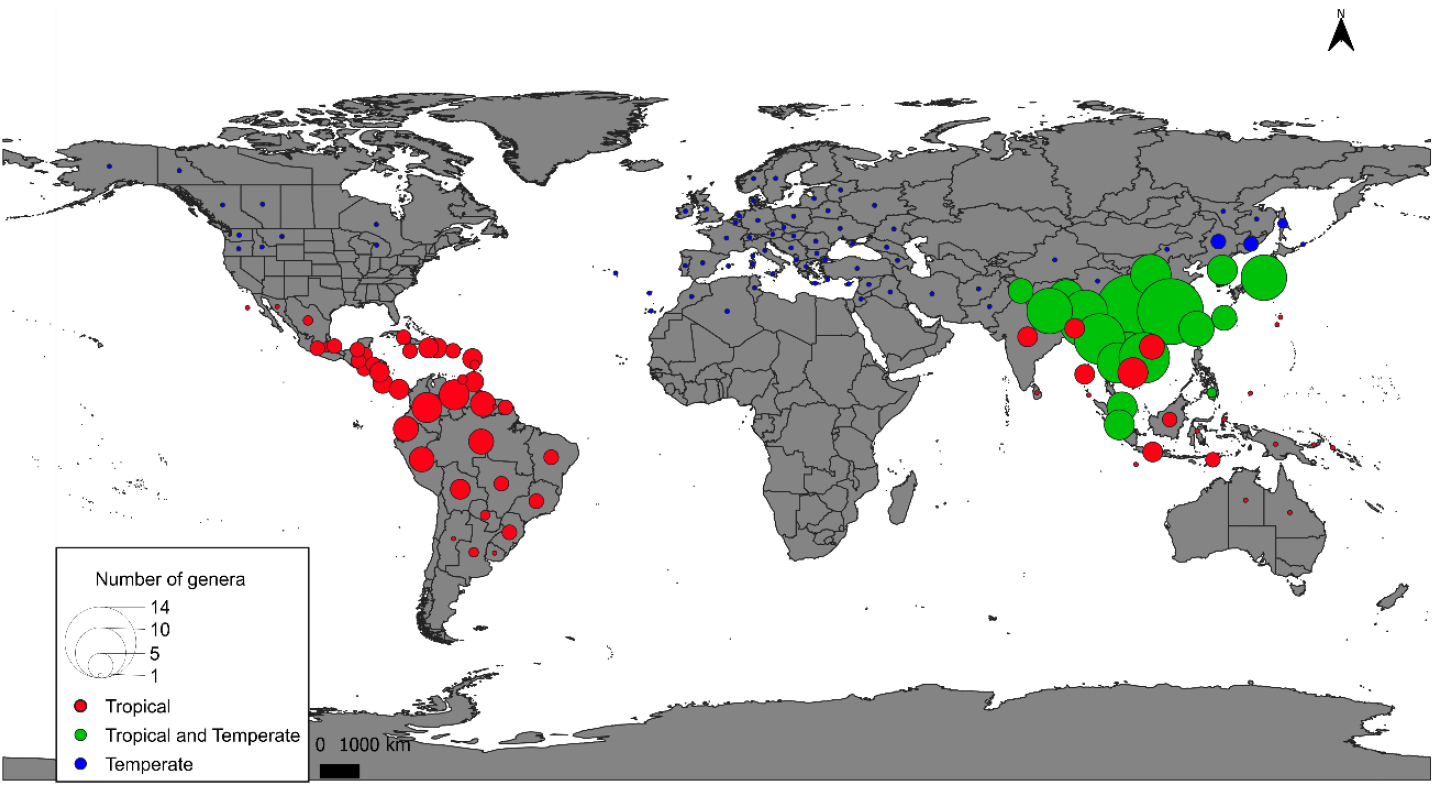
Biodiversity map of the Asian Palmate Group of Araliaceae as estimated from the number of genera per botanical country. The bubble colour depends on the climatic preferences of the genera recorded in each botanical country and according to the experts’ criteria (Plunkett et al., 1996; Wen et al., 2001): Blue represents botanical countries where only temperate genera are recorded, red those for which only tropical genera are recorded, and green those including records from temperate and tropical genera.

Also, the number of AsPG observations is unevenly distributed across the World (Fig. 3). The region of the World with the highest number of observations per area is Europe (only represented by *Hedera*) with a maximum number of observations per cell of 8,093. Most of the European sampling hotspots (i.e., cells with at least 2024 records that is 25% of the maximum sampling effort per cell in that area) are from central Europe (western England, France, Belgium and the Netherlands; Fig. 3A). The second region in terms of observations per area is Asia (maximum number of observations per cell: 1,469) where two sampling hotspots are identified (Taiwan and central Japan, Fig. 3B). The region of the World with the least number of records is America with a maximum number of observations per cell of 979 (Fig. 3C). The American sampling hotspots are concentrated in four main areas within the continent. Two of these hotspot areas are in Central America, one in southern Mexico and the other one in Panama and Costa Rica. The third main hotspot area is in South America and expands from western Ecuador to western Colombia. The fourth American hotspot area is in North America (western Washington State), where there is only one genus *Oplopanax* belonging to the AsPG clade.

**Figure 3.**
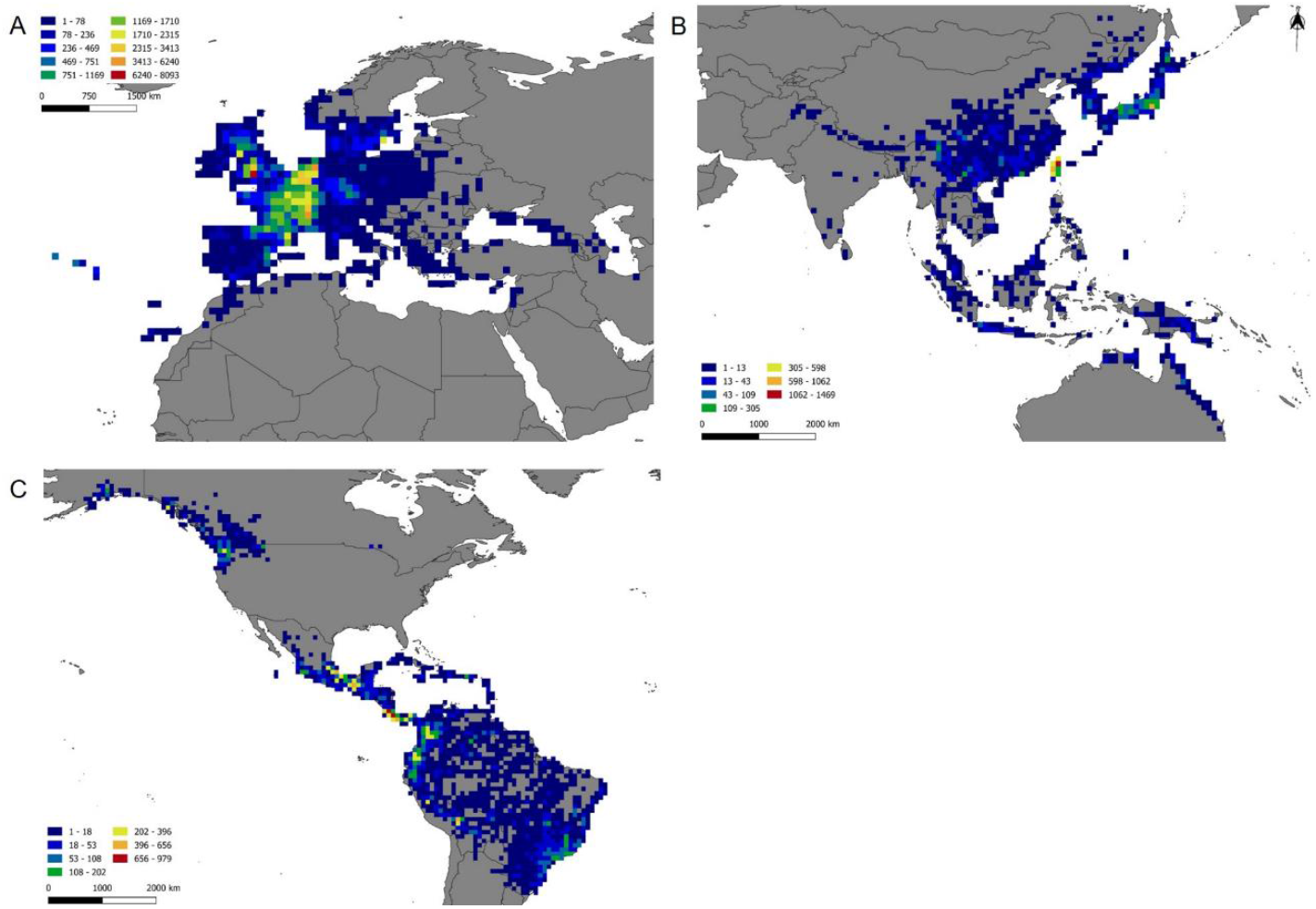
Heat maps of the Asian Palmate Group of Araliaceae based on the total number of point-occurrences per cell (one geographical degree of longitude and latitude). Cells are coloured based on a gradient scale established by Jenks natural breaks. Colour ranges from blue (least number of occurrences per area) to red (highest number of occurrences per area). (A) Europe (including North Africa), with a hotspot area located in the centre of the continent. The four last cell categories (from pale yellow -1,710 to 2,315 records- to red -6,240 to 8,093 records-) are identified as sampling hotspots (cells with 25% of the maximum sampling effort per cell per region, see Materials and methods). (B) Asia (including Oceania), with two main hotspot areas located in islands (Taiwan and Japan). The three last cell categories (from yellow - 305 to 598 records- to red -1,620 to 1,469 records-) are identified as sampling hotspots. (C) America, with four main hotspot areas from North America to Colombia. The three last cell categories (from yellow -202 to 396 records- to red -656 to 979 records-) are identified as sampling hotspots.

Finally, the analyses of sampling patterns across time and space reveal differences in the number of collections per year between temperate and tropical environments and among regions of the World (Fig. 4). In Asia, the number of baseline collections per year remained constant through time with three increases both for tropical and temperate genera according to the expert criterion (Fig. 4A). In the Americas, the number of collections across time describe very different temporal pattern whether the genera are tropical or temperate (Fig. 4B). Most of the temperate collections (only represented by *Oplopanax*) are concentrated in the second decade of the XXI^st^ century (2015 – 2020), whereas a relatively constant high baseline is observed for collections of the tropical genera during the last decades of the XX^th^ century with punctual increases concentrated in the first decades of the XXI^st^ century. Finally, in Europe where only one temperate genus, *Hedera*, occurs there is an increase of collections since 1990 with a peak in the first decades of the XXI^st^ century (Fig. 4C).

**Figure 4.**
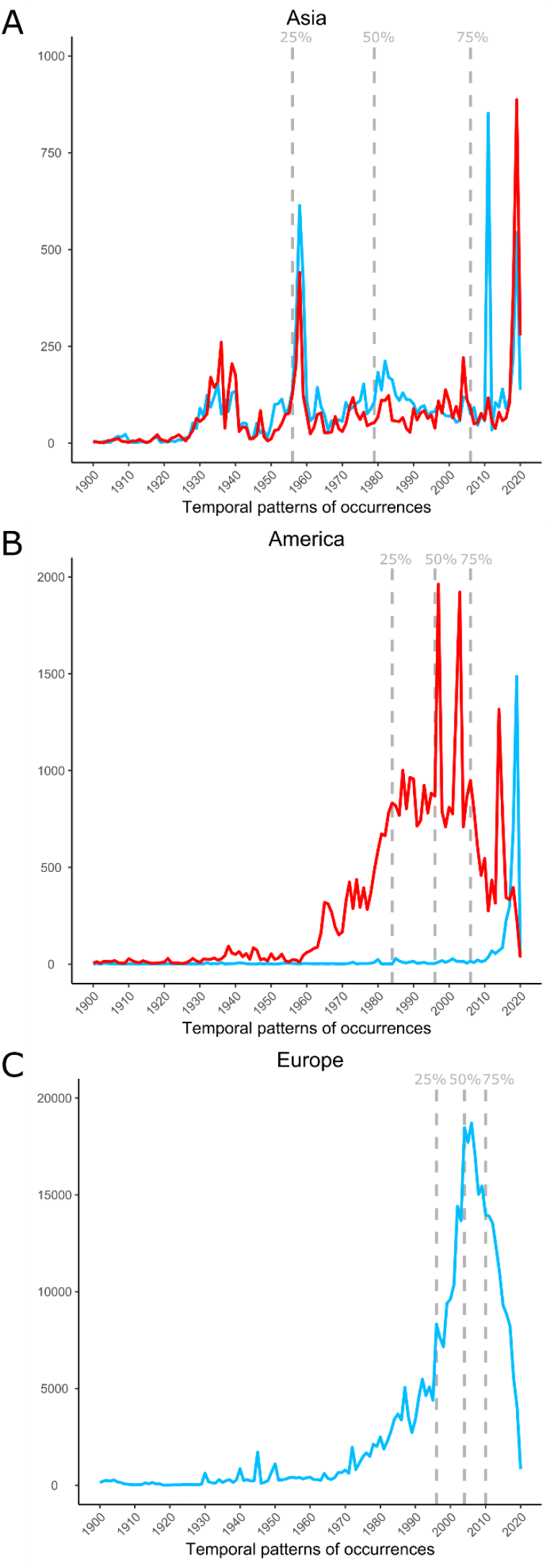
Temporal evolution of the number of occurrences included in the Asian Palmate Group of Araliaceae database between 1900 and 2020 by regions. Colours represent the temporal tendencies detected in the sampling records of temperate (blue) or tropical (red) genera according to the experts’ criteria (Plunkett et al. 1996, Wen et al. 2001). Dashed vertical lines indicate the cumulative frequencies of 25, 50 and 75 percentages of the total data. (A) Cumulative temporal series in Asia (including Oceania). (B) Cumulative temporal series in America. (C) Cumulative temporal series in Europe (including Africa), where only the temperate genus *Hedera* occurs.

### Comparison of World bioclimatic regionalizations

The regionalization of the World is very different depending on the bioclimatic classification system used (Appendix S3). These differences are not evenly distributed across the World or the climatic categories. Indeed, the regionalization of certain areas like low latitudes in the Northern Hemisphere are very similar among classifications, whereas the areas of the World that are classified as Subtropical (when this category is recognized; Holdridge’s, Meztger’s and Latitudinal classifications) are either considered tropical, temperate or dry in the remaining classification systems, which do not recognize the “Subtropical” category (Appendix S3).

Besides, the delimitation of the areas considered as tropical or temperate shows major differences even when classifications that do not recognize the “Subtropical” category are compared. Despite these major differences, there are two main spatial patterns in the regionalizations of the World across regions that emerge in all the classification systems used. First, the bioclimatic regions of North America and Australia extend over large continuous geographical areas describing a clear spatial pattern, either latitudinal or a combination of latitudinal and longitudinal in the classifications that recognize a “Dry” category (Appendix S3). Second, for the rest of the World the bioclimatic regions are patchy and extend over discontinuous geographical areas with no clear latitudinal or longitudinal pattern (Appendix S3).

### Spatial and climatic characterization of the AsPG

The spatial characterization of the genera distribution results in the identification of 11 restricted genera (*Brassaiopsis* Decne. & Planch., *Cephalopanax, Crepinella, Frodinia, Heteropanax, Macropanax, Merrilliopanax, Metapanax* J.Wen & Frodin, *Sinopanax* H.L.Li, *Tetrapanax* (K.Koch) K.Koch, and *Trevesia*; Table 2, Appendix S2) and four widespread genera with areas of occupancy larger than 10,000 km^2^ (*Dendropanax, Didymopanax, Hedera*, and *Oreopanax* Decne. & Planch.; Table 2).

The application of a semi-quantitative approach to characterize the climatic preferences of the AsPG genera reveals a general tendency for each genus to be classified in more than one climatic category in most of the classifications systems analyzed (Fig. 5). According to the most synthetic regionalization (Latitudinal classification; Fig. 5A), 15 genera have more than 75% of their occurrences classified in one category (hereafter “unequivocally assigned genera”): three as temperate (*Chengiopanax* C.B.Shang & J.Y.Huang, *Hedera* and *Oplopanax*), eight as tropical (*Cephalopanax, Crepinella, Dendropanax, Didymopanax, Frodinia, Oreopanax, Sciadophyllum* and *Trevesia*) and four as subtropical (*Merrilliopanax, Metapanax, Sinopanax* and *Tetrapanax*). The remaining eight genera have large proportion of records assigned to two categories (four as tropical and subtropical, four as temperate and subtropical; Fig. 5A). As the classification becomes more analytical, the number of genera unequivocally assigned to one category decreases up to 10 in Köppen’s classification or to none in Holdridge’s or Metzger’s classifications. The only exception is the Simplified Ecoregions classification, where 20 genera are unequivocally assigned to one category (Fig. 5). For the classifications that do not recognise the “Subtropical” category, most of the non-unequivocally classified genera are assigned to the “Tropical” and “Temperate” categories (Köppen’s: 10, Simplified Ecoregions: five; Figs. 5B, 5E). For Holdridge’s and Metzger’s classifications, which recognize the “Subtropical” category, most of the genera classified in more than one category do indeed include the “Subtropical” category as one of them (Fig. 5C-D).

**Figure 5.**
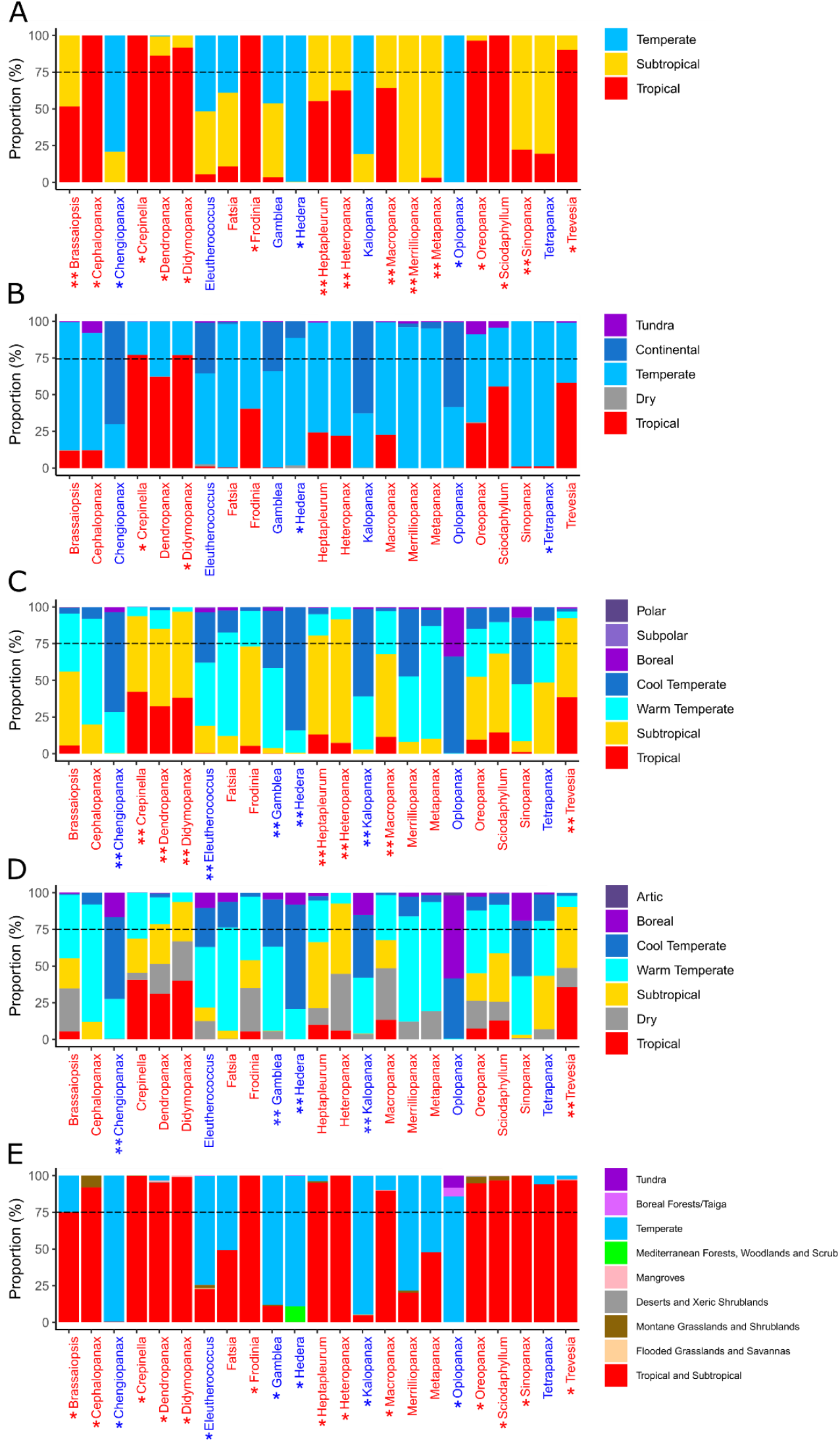
Climatic characterization of the 23 genera of the Asian Palmate Group (AsPG) of Araliaceae according to five bioclimatic regionalization systems of the World. The names of AsPG genera in the first axis are coloured according to experts’ criteria (tropical in red and temperate in blue; Plunkett et al. 1996, Wen et al. 2001). One asterisk indicates the genera for which the classification system assigned a climatic categorization congruent with the one assigned in the experts’ criteria. Two asterisks indicate the congruence with the “Tropical” category of the experts’ criteria when unifying the subtropical and the tropical proportions in the classification system. (A) Latitudinal zonation. (B) Köppen’s classification based on temperature and precipitation (Köppen and Geiger, 1936; Beck et al., 2018). (C) Holdridge’s classification based on Biotemperature (Holdridge, 1967). The congruence with the “Temperate” category of the experts’ criteria is done by considering the Cold and Warm temperate categories together and it is also denoted with two asterisks. (D) Metzger’s classification based on 42 bioclimatic variables (Metzger et al., 2012). Congruence with the “Temperate” category of the experts’ criteria is done by considering the Cold and Warm temperate categories together and it is also denoted with two asterisks. (E) Simplified Ecoregions classification based on environmental conditions and the biogeographical information of the World’s floras and faunas (modified from Dinerstein et al., 2017). “Tropical and Subtropical” category unifies four original biomes of the Dinerstein regionalization (“Tropical and subtropical dry broadleaf forests”, “Tropical and subtropical moist broadleaf forests”, “Tropical and subtropical coniferous forests” and “Tropical and subtropical grasslands, savannas, and shrublands”). “Temperate” category includes three original biomes of Dinerstein regionalization (“Temperate broadleaf and mixed forests”, “Temperate conifer forests”, and “Temperate grasslands, savannas, and shrublands”).

Major differences are detected in the climatic preferences assigned to genera among the five classifications compared and large discrepancy with the climatic characterization of the expert criterion (Fig. 5). The classification system that reveals more similarity with the expert criterion is the Simplified Ecoregions (Fig. 5E), where 19 genera are included in the same category (six temperate, 13 tropical; see asterisks in Fig. 5E). The classifications that depart the most from the expert criterion are the other two most analytical ones (Holdridge’s and Meztger’s classifications), where only one genus is respectively included in a similar category (see genus with one asterisk in Fig. 5C-D), followed by the more synthetical classification of Köppen-Geiger with five genera classified in the same category as in the expert criterion (two tropical, three temperate; see genera with one asterisk in Fig. 5B). In between, the latitudinal classification recovers 11 similarities (three temperate, eight tropical; see genera with one asterisk in Fig. 5A) or 19 if the “Subtropical” category is unified with the “Tropical” one (see double asterisks in Fig. 5A).

When projected in a spatial context, the areas of the World where the AsPG occurs are recognized as different bioclimatic categories depending on the classification used (compare Fig. 1 and Appendix S3). Indeed, the percentage of AsPG records assigned to each category varies largely depending on the classification analysed (Fig. 6). For example, when classifications that recognize the “Subtropical” category are compared, the percentage of occurrences assigned to the “Tropical” category may vary from only 10% according to Metzger’s classification (Fig. 6E) to 43% in the latitudinal zonation (Fig. 6B), or between 33% (Latitudinal zonation, Fig. 6B) and 55% (Holdridge’s classification, Fig. 6D) for the “Temperate” category. Also, the percentage of occurrences assigned to the “Subtropical” category varies from 16% in Metzger’s classification (Fig. 6E) to 31% in Holdridge’s (Fig. 6D).

**Figure 6.**
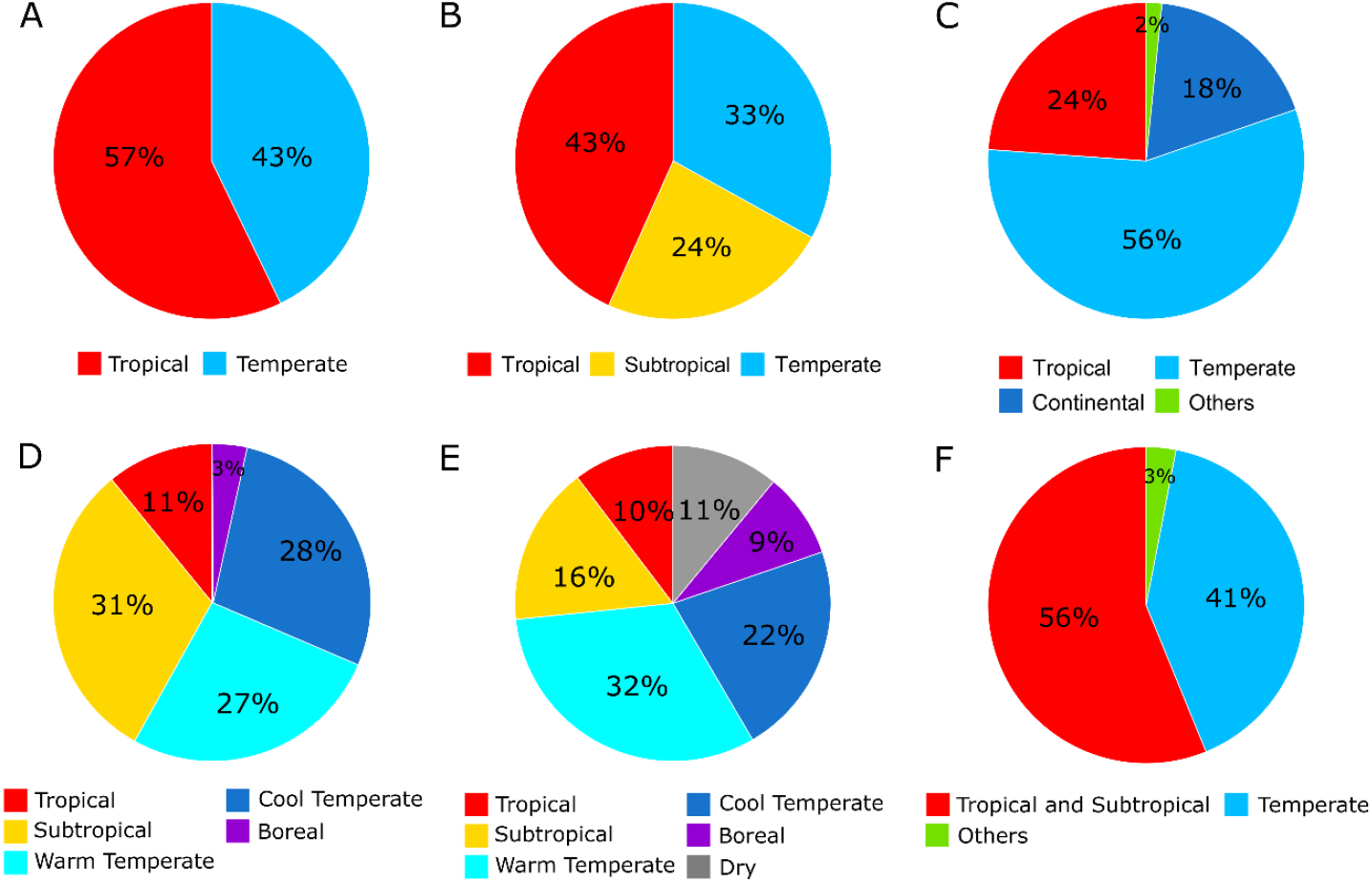
Climatic characterization of the Asian Palmate Group of Araliaceae according to five bioclimatic regionalization systems of the World and the experts’ criteria (Plunkett et al., 1996; Wen et al., 2001). Categories represented by less than 0,1% of the total records are not represented. Categories represented by less than 1% of the total records are grouped under the undefined category “Others”. (A) Experts’ criteria (Plunkett et al., 1996; Wen et al., 2001). (B) Latitudinal zonation. (C) Köppen’s classification based on temperature and precipitation (Köppen and Geiger, 1936; Beck et al., 2018). (D) Holdridge’s classification based on Biotemperature (Holdridge, 1967). (E) Metzger’s classification based on 42 bioclimatic variables (Metzger et al., 2012). (F) Simplified Ecoregions classification based on environmental conditions and the biogeographical information of the World’s floras and faunas (modified from Dinerstein et al., 2017). “Tropical and Subtropical” category unifies four original biomes of the Dinerstein regionalization (“Tropical and subtropical dry broadleaf forests”, “Tropical and subtropical moist broadleaf forests”, “Tropical and subtropical coniferous forests” and “Tropical and subtropical grasslands, savannas, and shrublands”). “Temperate” category includes three original biomes of Dinerstein regionalization (“Temperate broadleaf and mixed forests”, “Temperate conifer forests”, and “Temperate grasslands, savannas, and shrublands”).

## DISCUSSION

### Spatial gaps of knowledge in the AsPG distribution

After the quality control was done on the spatial data gathered from online data sources to represent the distribution of the AsPG genera, we needed to fill spatial and taxonomical gaps of knowledge to avoid reaching misleading biodiversity patterns (Soberón and Peterson, 2004; Boakes et al., 2010). As a result, we herein provide a comprehensive spatial database that provides good taxonomical, geographical and temporal coverage of the AsPG at the genus level (Coca-de-la-Iglesia et al., 2021a; Appendix S2). A bias in sampling effort is detected in our database with the highest number of sampling hotspots detected in Europe and the lowest in Asia and America (Fig. 3). Similar geographically-biased patterns towards a better biodiversity documentation in Europe have already been reported in other studies using digital accessible information (Boakes et al., 2010; Meyer et al., 2016). However, in the AsPG case the sampling bias display a spatial pattern that is just the opposite as the biodiversity pattern since Asia, which is the richest region in terms of genera, arises as the with the region with the least sampling effort while Europe, which is the region with the lowest diversity in terms of genera, arises as the area with the highest sampling effort (Fig. 2). This mismatch between biodiversity documentation and biodiversity pattern have been generally attributed to non-biodiversity factors (Hortal et al., 2007; Yesson et al., 2007; Amano and Sutherland, 2013; Yang et al., 2013). Interestingly, a similar mismatch pattern is observed for the AsPG when downsizing the scale both in Europe and in Asia. Indeed, all European sampling hotspots are in Central Europe, where only one species of *Hedera* occurs, while Southwestern Europe that harbours the greatest diversity including all the European continental species of *Hedera* (Valcárcel et al., 2003, 2017) emerged as an under sampled area. Also, the lowest sampling effort detected in Asia is in Southwestern continental China (Fig. 3B) which is the area with the greatest AsPG generic diversity (Fig. 2). This calls for improved economic funding policies to fill our gaps of biodiversity knowledge (Wen et al., 2015, 2017). We propose a two-steps approach targeted on the areas where sampling effort does not reflect biodiversity patterns, such as Southeastern and Southwestern Europe or Southwestern Asia in the AsPG case. First, the development of geo-referencing and digitization projects focused on regional and National herbarium collections to fill the gaps attributed to insufficient digitization effort or lack of coordinates in specimens’ information, which allows minimizing costs while maximizing efficiency. Then, targeted field sampling campaigns for the regions where the taxonomical coverage remain poor after the digitization process. At a global scale, the targeted areas to fill the AsPG geographical gaps are the boundaries of certain botanical countries in Asia (Bangladesh, Nepal, Bhutan and India; see *Brassaiopsis, Dendropanax, Eleutherococcus* Maxim., *Gamblea, Heteropanax, Macropanax, Merrilliopanax, Metapanax* and *Trevesia*) and America (French Guiana: *Dendropanax*; Uruguay: *Didymopanax* and Surinam: *Oreopanax*; Appendix S1). At a regional scale, our targeted areas are Southwestern Europe for which the geographical and taxonomical coverage of our database remains poor.

### Selection of regionalization systems to characterize the climatic preferences of taxa: a matter of where you are and who you are

The five bioclimatic classification systems analysed are standard regionalizations of the World’s climate that have been applied to many different purposes. They have been used to respond to general questions and evaluate general circulation models (Lohmann et al., 1993) or detect climate change (Beck et al., 2005). Alternatively, they have also been applied for specific biological approaches, like identification of evolutionary patterns (Anest et al., 2021) description of macroecological patterns (Ruhí et al., 2013) or setting conservation planning strategies (Ricketts and Imhoff, 2003). The application in which we focused is the characterization of climatic preferences of taxa to identify and understand diversity patterns and the underlying evolutionary history (i.e., Wen et al., 2001; Silva et al., 2021). As important as climatic information might be to understand the evolutionary history of lineages and the resulting diversity trends (Kozak and Wiens, 2012), the specific climatic criterion used to determine the preferences of taxa is rarely stated in evolutionary studies (Feeley and Stroud, 2018). While very often *ad hoc* qualitative characterization is done based on expert criterion or on rough proxies such as the application of the latitudinal criterion (Kozak et al., 2008). This is so because, the alternative quantitative climatic characterization needs from a high-quality point-occurrence database with good geographical coverage of the taxa (Hortal et al., 2007, 2015; Meyer et al., 2016), which is generally missing due to our pervasive taxonomical and geographical gaps of knowledge in most lineages across the tree of life (Meyer et al., 2016). For these cases, a careful selection of the criterion used for the climatic characterization of taxa should be done since the application of one or another criterion results in very different conclusions (Feeley and Stroud, 2018). Our comparative approach confirms previous findings for the tropics (Feeley and Stroud, 2018) and reveal large differences in the delimitation of the climatic regions in the World depending on the classification system used. We are aware that part of these differences rests on the variables used in each system and on how analytical or synthetical the classification is. However, we herein stress on the part of the discrepancies among classification systems that is due to the oxymoron of using categorical classification systems to capture the dynamic and transitional nature of climate. In any case, from the similarities and differences detected among classifications some patterns emerge that may help minimizing the impact of our assumptions when using a qualitative proxy to define the climatic preferences of taxa. First, the climatic affinity occurring at high latitudes in the Northern Hemisphere are very well captured in all classifications irrespective to the variables used or the categories recognized (compare Polar in Latitudinal classification, Appendix S3.1; to Tundra and Frost in Köppen, Appendix S3.2; Subpolar and polar in Holdridge’s, Appendix S3.3; Artic in Meztger’s, Appendix S3.4; and Tundra in Simplified ecoregions, Appendix S3.5). As a result, the selection of one classification system or another may be irrelevant as to reach robust conclusions for taxa occurring in such latitudes. Second, for the New World (particularly for North America and Australia), the regionalizations result in regular shapes that describe a clear latitudinal or the combination of latitudinal and longitudinal pattern when a “Dry” category is recognized (Appendix S3). This allows delimiting different but compatible continuous bioclimatic regions whether the regionalization used is one or another (Appendix S3). Thus, classification systems can eventually capture the variation in climatic preferences of taxa from these areas. In such cases, selection of the best-fitting regionalization is however key to obtain robust conclusions and ultimately depends on the range of the taxa and the question addressed. Third, bioclimatic zones in the Old World are irregularly shaped and do not follow any clear spatial pattern. The impact of irregularly shaped regions on climatic regionalizations has already been analysed (Aydin et al., 2021) and, as in our study case, result in the delimitation of highly patchy and incompatible regions based on the different classification systems (Appendix S3). Therefore, the use of this qualitative proxy to characterize the climatic preference of taxa in the Old World may be more difficult, in the first place, and may end in conflicting signals leading to contrasting conclusions whether the classification system chosen is one or another.

Interestingly, the fact that major conflicts among classification systems are detected in the areas of the World that are recognized within a “Subtropical” category may point to the already stated limitation of the available classification systems to capture the transitional nature of climate. In fact, this question may be partly responsible for the major differences in perspectives between biogeographers and phylogenetists on what and where the tropics are (Feeley and Stroud, 2018). Consequently, it may also affect our delimitation and understanding of the Temperate zones as well. Since, tropics are often interpreted as opposed to temperate to explain large scale biogeographic and diversity patterns (Wiens and Donoghue, 2004), we should be more cautious on the climatic characterization step in evolutionary studies.

### Climatic preferences within the AsPG are not as contrasted as previously stated

According to the experts’ criteria, 16 genera of the AsPG are tropical while only seven temperate (Plunkett et al., 1996; Wen et al., 2001). However, this temperate vs. tropical dichotomy is no longer supported when an explicit evaluation of other climatic classification systems is done due to two main findings.

First, the idea that each genus can be characterized by the unique climatic conditions of one single category that is implicitly assumed in the expert criterion, is rejected when using our semiquantitative approach since most of the genera are classified in two or more categories (Fig. 5). Classifying one genus in more than one category can be interpreted as evidence of a broad climatic niche. Broad climatic preferences are expected for taxa with wide ranges since geographical extent is correlated to realized climatic niches (Slatyer et al., 2013). However, in our case study, most of the genera are classified in more than one category in any of the classification systems (Fig. 5) irrespective of the extension of their geographical ranges (Table 2). Indeed, not only the most widespread genera are assigned to more than one category (*Didymopanax, Dendropanax* or *Oreopanax*, Table 2; Fig. 5B-D), but also most of the genera with the narrowest distributions (*Cephalopanax, Frodinia* or *Merrilliopanax*, Table 2; see Fig. 5B-5D). Also, some of the most widespread genera are unequivocally assigned to one category (*Hedera* or *Oplopanax*, Table 2, Fig 5). In a recent study, Liu et al. (2020) tested multiple factors to evaluate general hypotheses on climatic niche trends in plants and animals, such as differential correlation between temperature and precipitation niche breadths across latitude or the influence of within-locality niche breadth to understand global scale niche breadths. The consideration of these patterns does not help explaining our contrasting results either, since latitude does not seem to be related in the AsPG to the number of climatic categories recognised per genus nor the local niche breadth (Alonso et al., 2021). Interestingly, classifying one genus in more than one category can be also interpreted as evidence that the categories stablished in the classification systems do not capture well the variability of the climatic preferences of the genus. This does not mean the classification is not robust or accurate enough, it only means that the regionalization of the World according to the given classification does not reflect the climatic variability when it comes to represent the geographical range of the genus (note that the genus range does not necessarily span over the whole area that covers a given category).

Second, in most of the AsPG genera that are unequivocally classified in a given category, this category is different than the “Tropical” or “Temperate” categories recognized in the expert criterion (“Subtropical”, “Warm temperate”, “Cool temperate”, “Continental” or “Boreal”). Indeed, only three genera display similar climatic preferences according to the five classifications and the expert criterion (*Cephalopanax, Chengiopanax* and *Hedera*), whereas the remaining genera show large incongruences. Most of the incongruences detected are due to the assignment of the genera to a Subtropical or Warm temperate category in the classification systems that recognise these categories (Holdridge, 1967; Metzger et al., 2012; Figs. 5A, 5C-5D). These two categories reflect intermediate and progressive climatic conditions between the ones that characterize the “Tropical” and “Temperate” or “Cool temperate” categories (Holdridge, 1967; Metzger et al., 2012). Indeed, the regions of the World that are classified within these two intermediate categories (Subtropical and Warm temperate) coincide with areas that are primarily recognised as Tropical or Temperate in the other classifications (Appendix S3). Thus, we interpret that part of the inconsistencies detected among classifications for the climatic characterization of the AsPG genera does not reflect actual incongruences, but an artifact derived from the limitations of the available World regionalizations to capture the transitional and dynamic nature of climate.

In the light of our results, we reach two conclusions. First, the climatic characterization of the AsPG genera has an inherent complexity due to the geographical regions where they occur. Second, the application of the expert criterion or any of the other classifications for the climatic characterization of the AsPG genera may lead us to reach inconsistent observations. Therefore, the seemingly easiness of the AsPG lineages to shift climatic niches inferred from the scattered phylogenetic placement of “Temperate” genera in molecular studies (Valcárcel and Wen, 2019; Valcárcel et al. 2014; Wen et al., 2001) needs to be reevaluated by using a quantitative approach rather than a qualitative one. Also, the fact that most of the genera are classified within categories that represent intermediate climatic conditions within this classical example of a tropical-temperate family (Plunkett and Lowry, 2001), calls for a general revision of other cases in Apiales (Plunkett et al., 2004) and in other tropical-temperate plant lineages, such as Vitaceae (Wen et al., 2018; Ma et al., 2021), Altingiaceae (Ickert-Bond and Wen, 2013), *Prunus* (Hodel et al., 2021), and *Rhaphiolepis* (Liu et al., 2020). Furthermore, our hypotheses from this study will need to be further tested as more collections become digitized and generally accessible to the biodiversity community (Wen et al., 2015, 2017; Funk, 2018; Wen and Wagner, 2020).

## Supporting information

Heat maps of Asian Palmate Group genera

Distribution maps of Asian Palmate Group genera

World regionalizations

## ACKNOWLEDGMENTS

The authors thank R. Li for providing a curated checklist of the former *Schefflera* of the AsPG and P.P. Lowry II for anticipating the taxonomical rearrangements of the former *Schefflera* before the papers were published. We also thank A. Gallego-Narbón for taxonomical updates and I. Datta de Castro for technical support on cluster analysis. We also indebted to the people who are part of the Writing Workshop developed by the Biology and Ecology Departments of the Universidad Autónoma de Madrid, for all the comments and discussions that have helped to realize this work. Finally, we specially thank all scientists, naturalists and botanical enthusiasts who have contributed to open access databases through botanical collections and observations during the last centuries, which made this study possible.

This study was supported by the Spanish Ministry of Economy, Industry and Competitiveness [CGL2017-87198-P] and the Spanish Ministry of Science an Innovation [PID2019-106840GA-C22]. M. Coca de la Iglesia was supported by the Youth Employment Initiative of European Social Fund and Community of Madrid [PEJ-2017-AI-AMB-6636 and CAM_2020_PEJD-2019-11 PRE/AMB-15871].

## AUTHOR CONTRIBUTIONS

Database compilation: MC & JW. Data cleaning: MC, NGM, JW & VV. Spatial and climatic analyses: MC. Methodological design: MC, NGM & VV. Results interpretations and Manuscript preparation: MC, NGM, JW & VV

## DATA AVAILABILITY STATEMENT

The AsPG database used in this paper is available at Zenodo Repository: https://doi.org/10.5281/zenodo.5578149.

The elevation layer of the World is available in Zenodo Repository: https://doi.org/10.5281/zenodo.5578234.

The five spatial layers of the bioclimatic classifications used in this paper are available in GitHub Repository: https://github.com/vvalnun/Bioclimatic-classifications-AsPG.git.

## SUPPORTING INFORMATION

Appendix S1 – Heat maps of Asian Palmate Group genera

Appendix S2 – Distribution maps of Asian Palmate Group genera

Appendix S3 – World regionalizations

## Notes

https://zenodo.org/record/5578234

https://zenodo.org/record/5578149

https://github.com/vvalnun/Bioclimatic-classifications-AsPG.git

### Competing Interest Statement

The authors have declared no competing interest.

