## Supplementary material for "Tropical-temperate dichotomy falls apart in the Asian Palmate Group of Araliaceae": Distribution maps of Asian Palmate Group genera

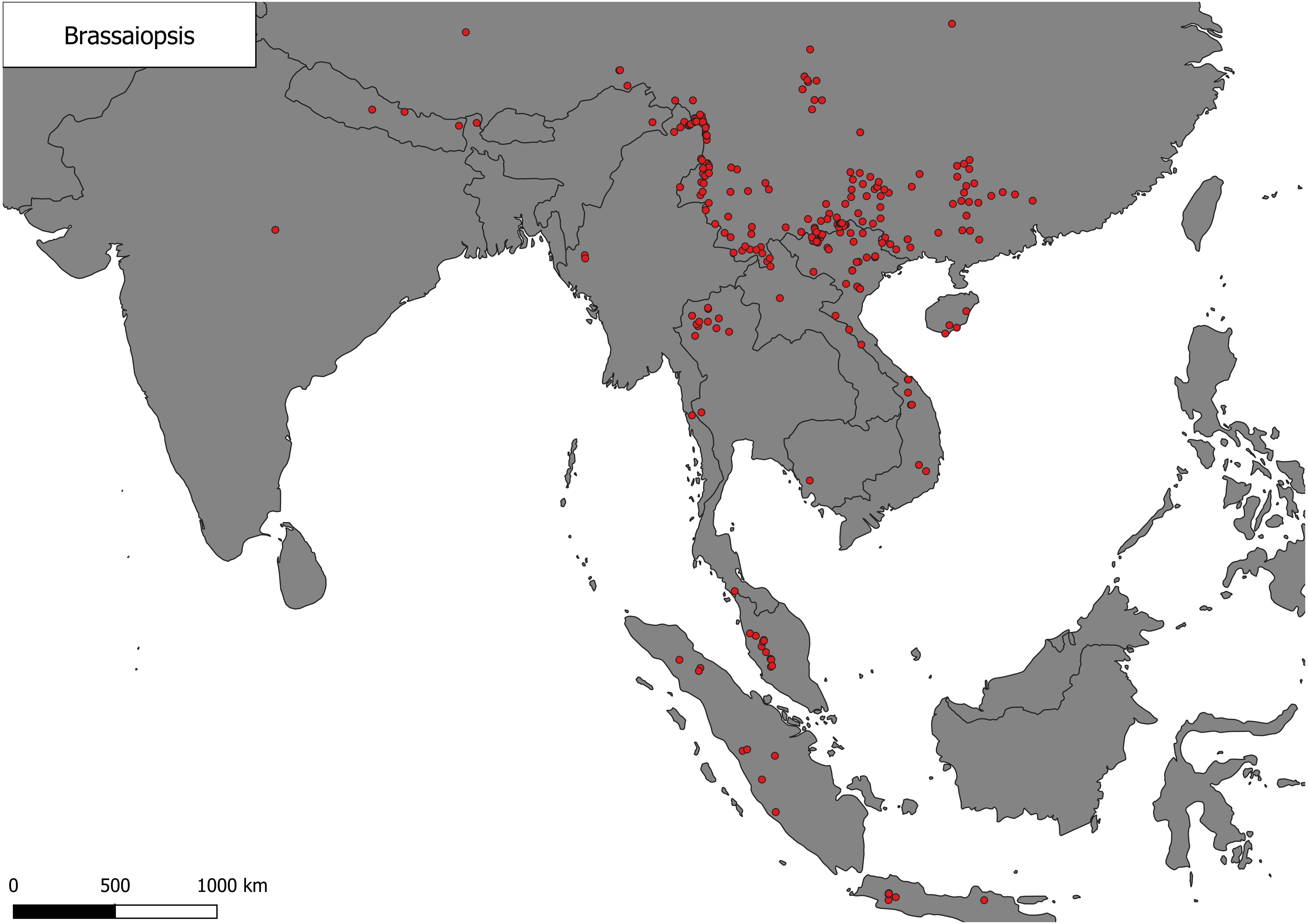
Brassaiopsis
0
500
1000 km

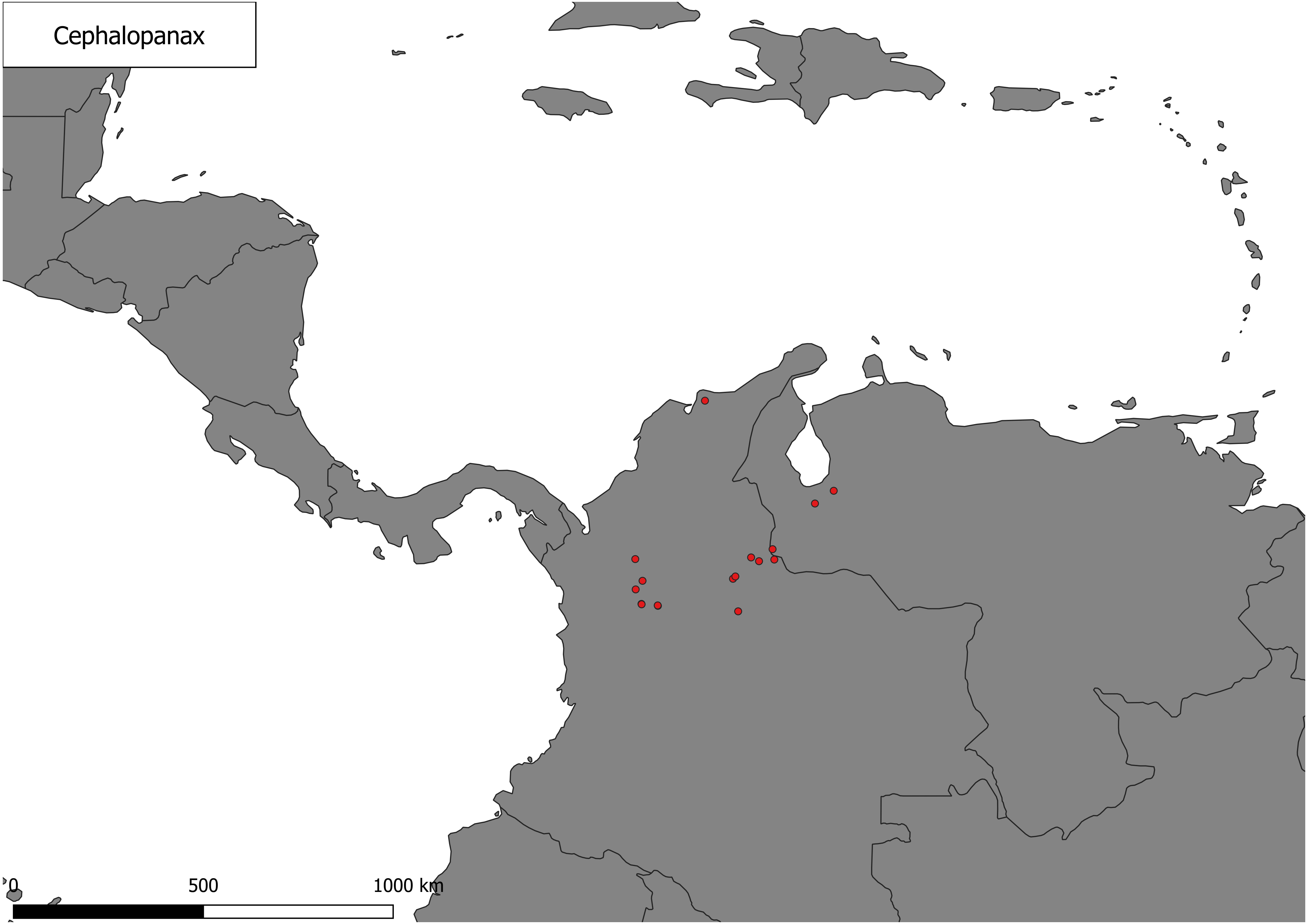
Cephalopanax
0
500
1000 km

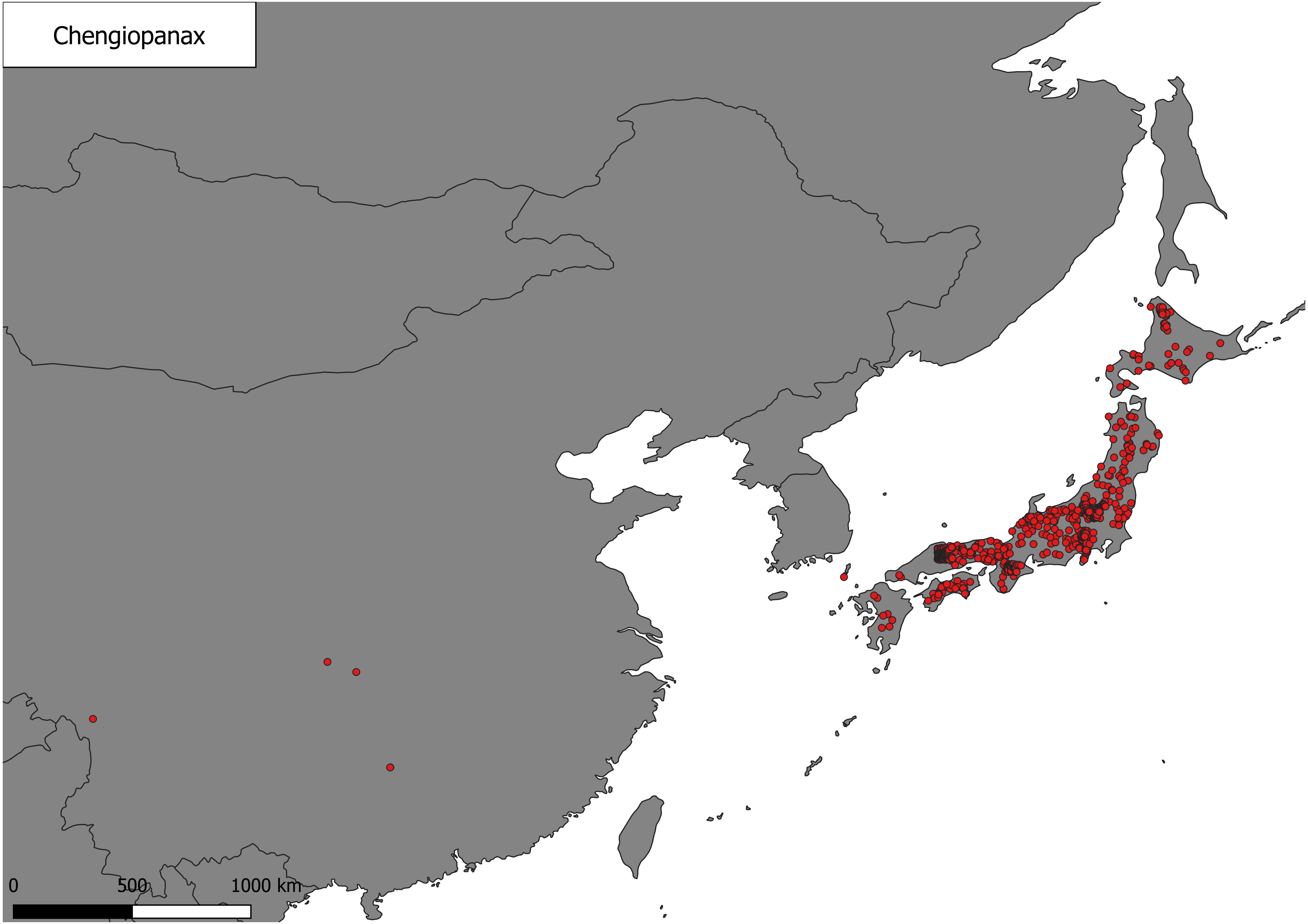
Chengiopanax
0
500
1000 km

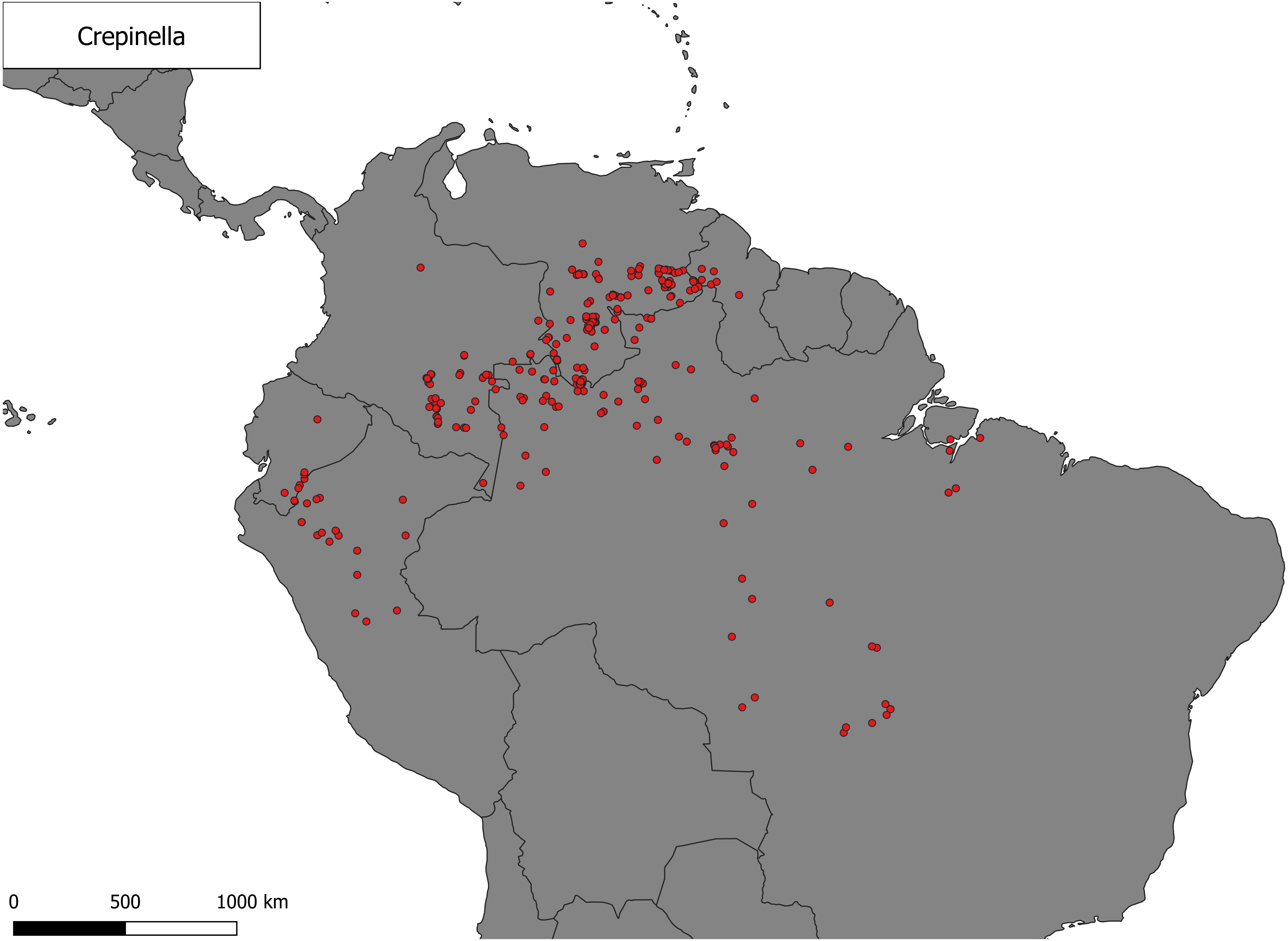
Crepinella
0
500
1000 km

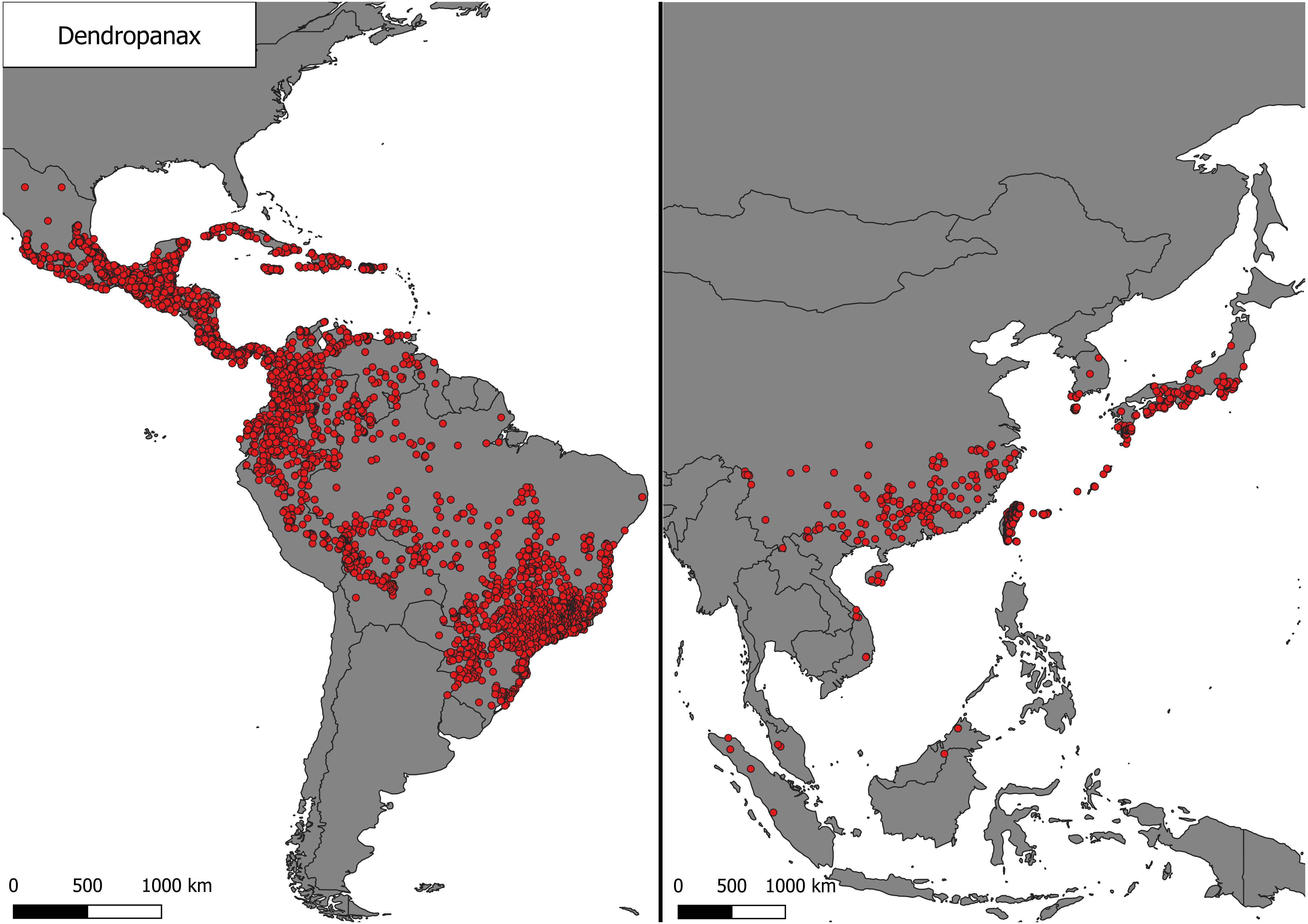

Dendropanax
0
500
1000 km
0
500
1000 km

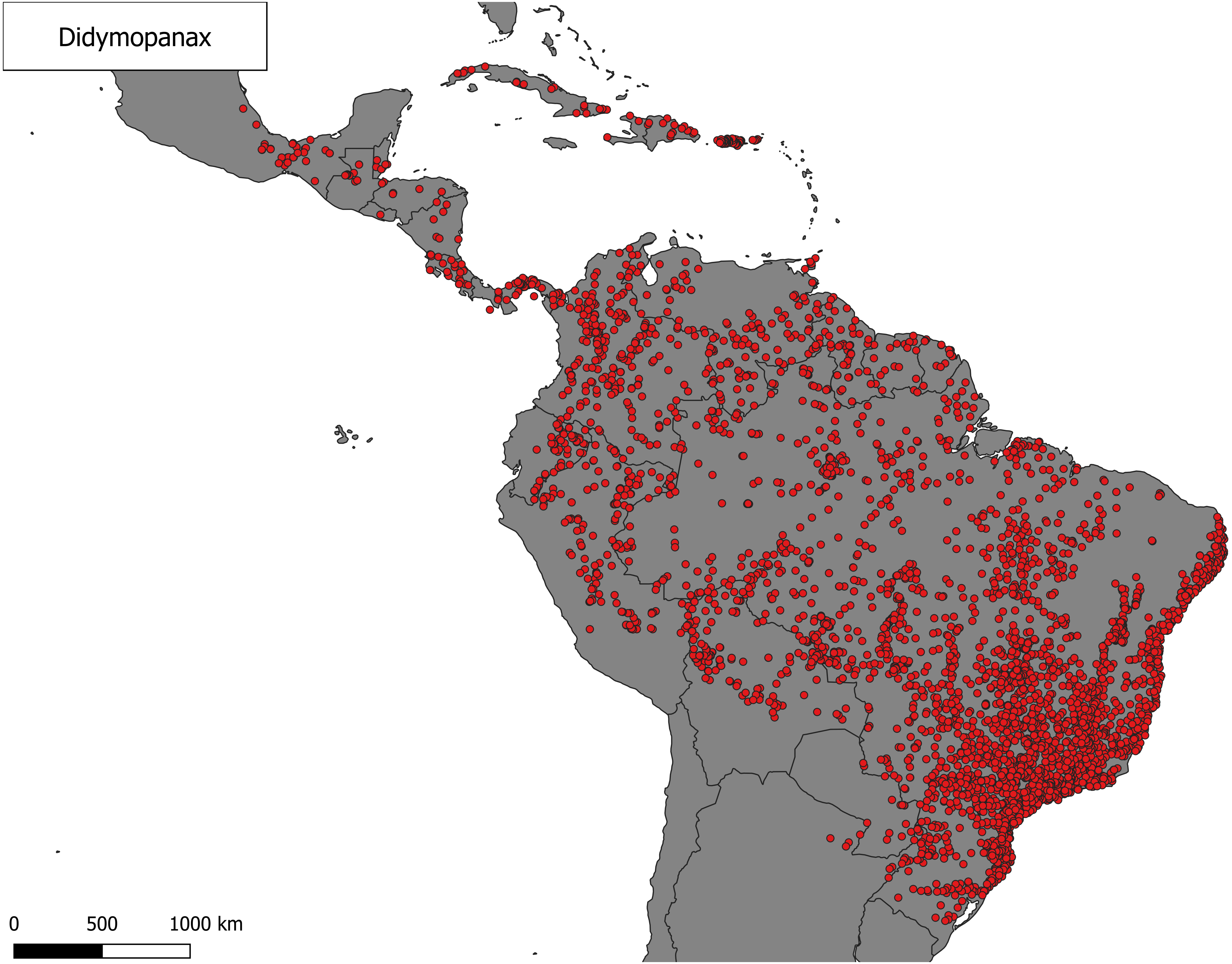
Didymopanax
0
500
1000 km

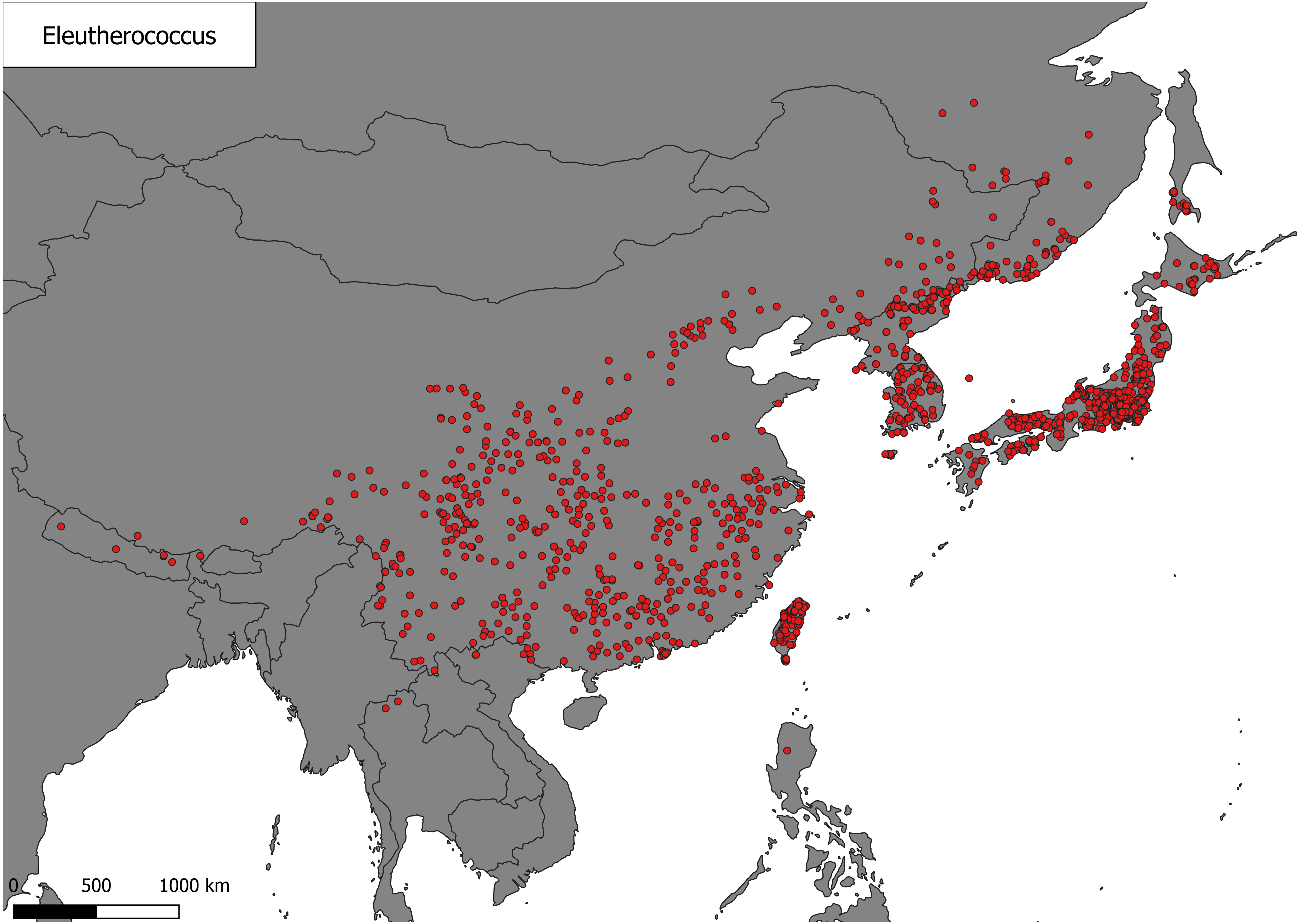

Eleutherococcus
0
500
1000 km

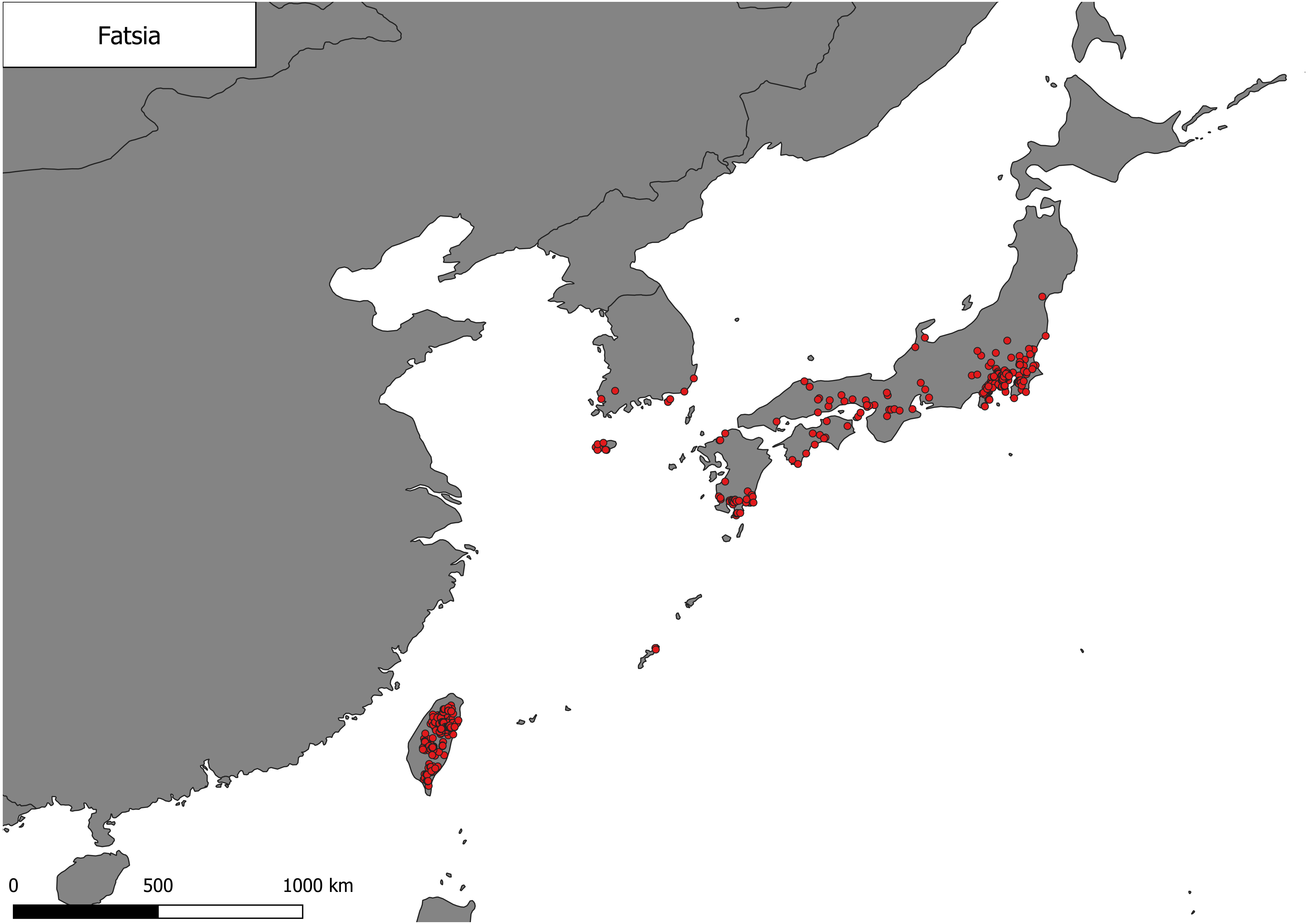
Fatsia
0
500
1000 km

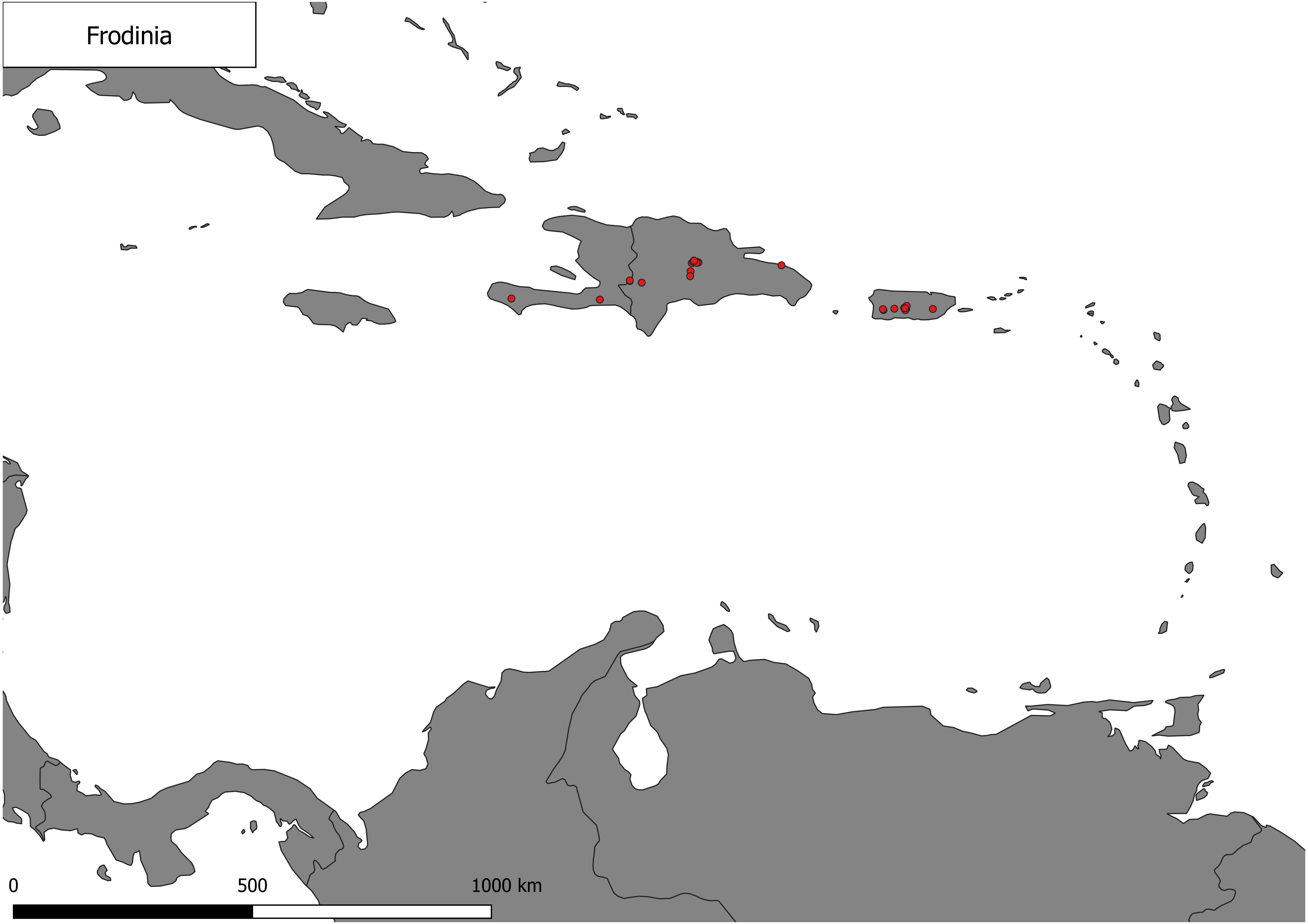
Frodinia
0
500
1000 km

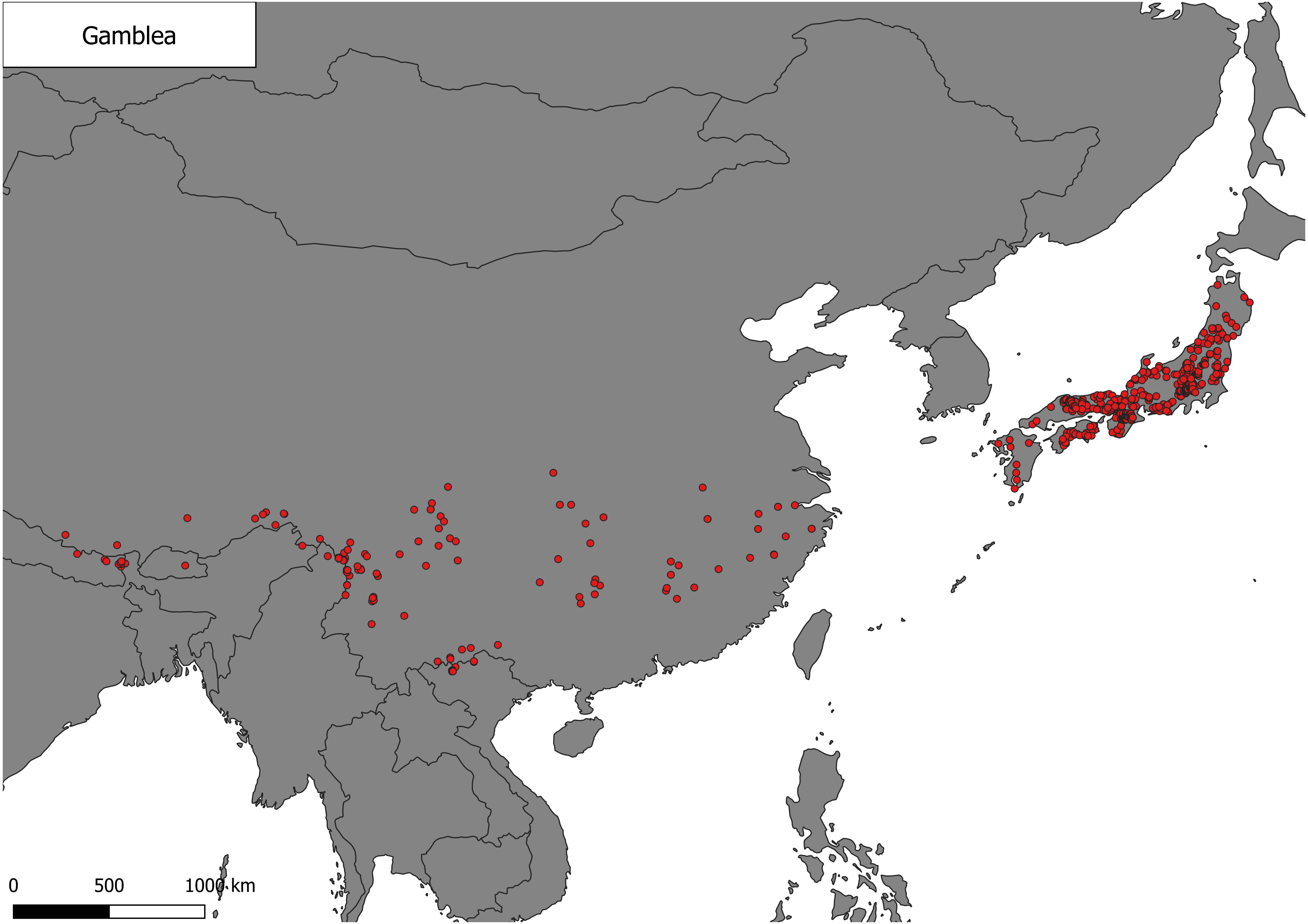
Gamblea
0
500
1000 km

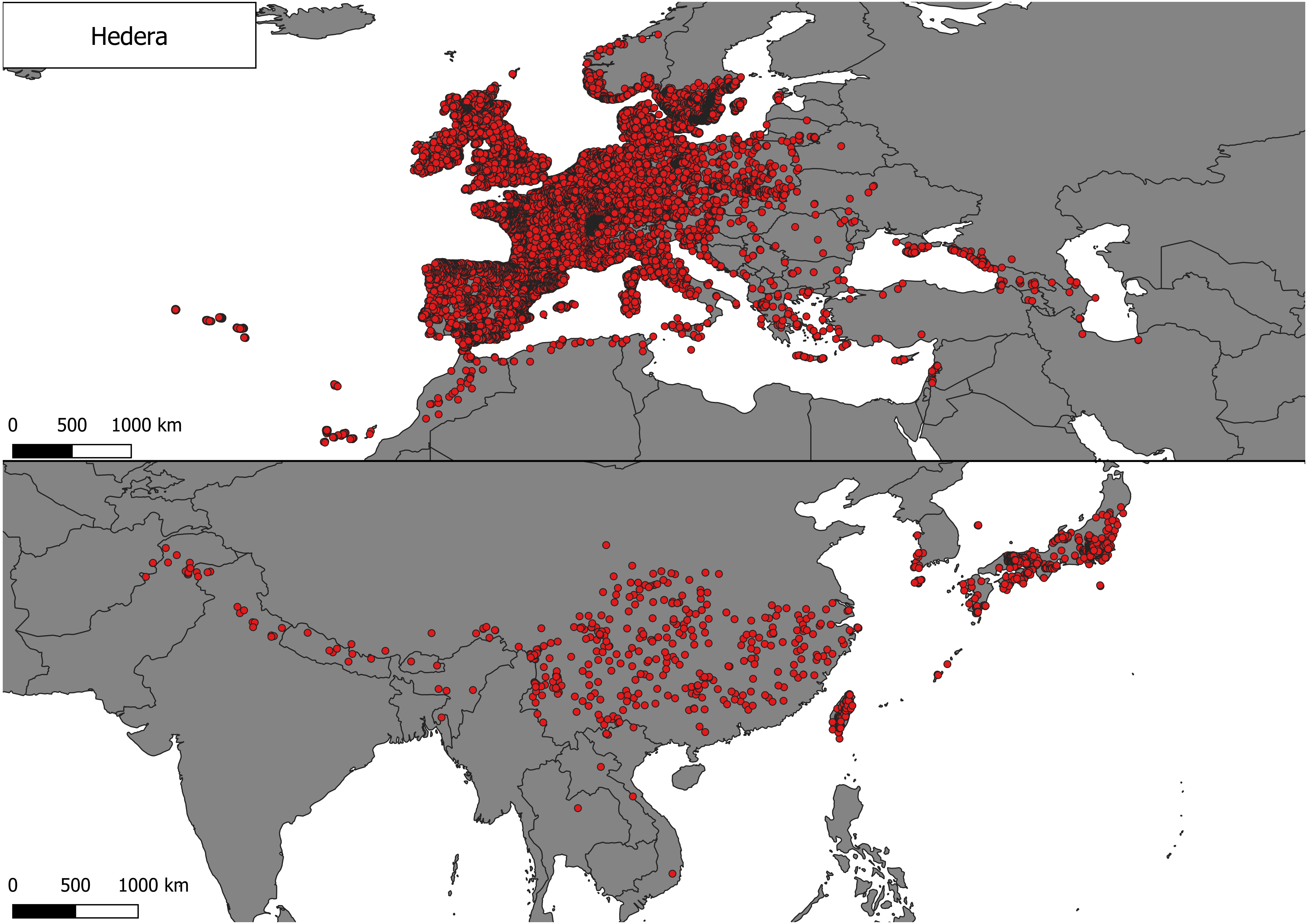

Hedera
0 500 1000 km
0 500 1000 km

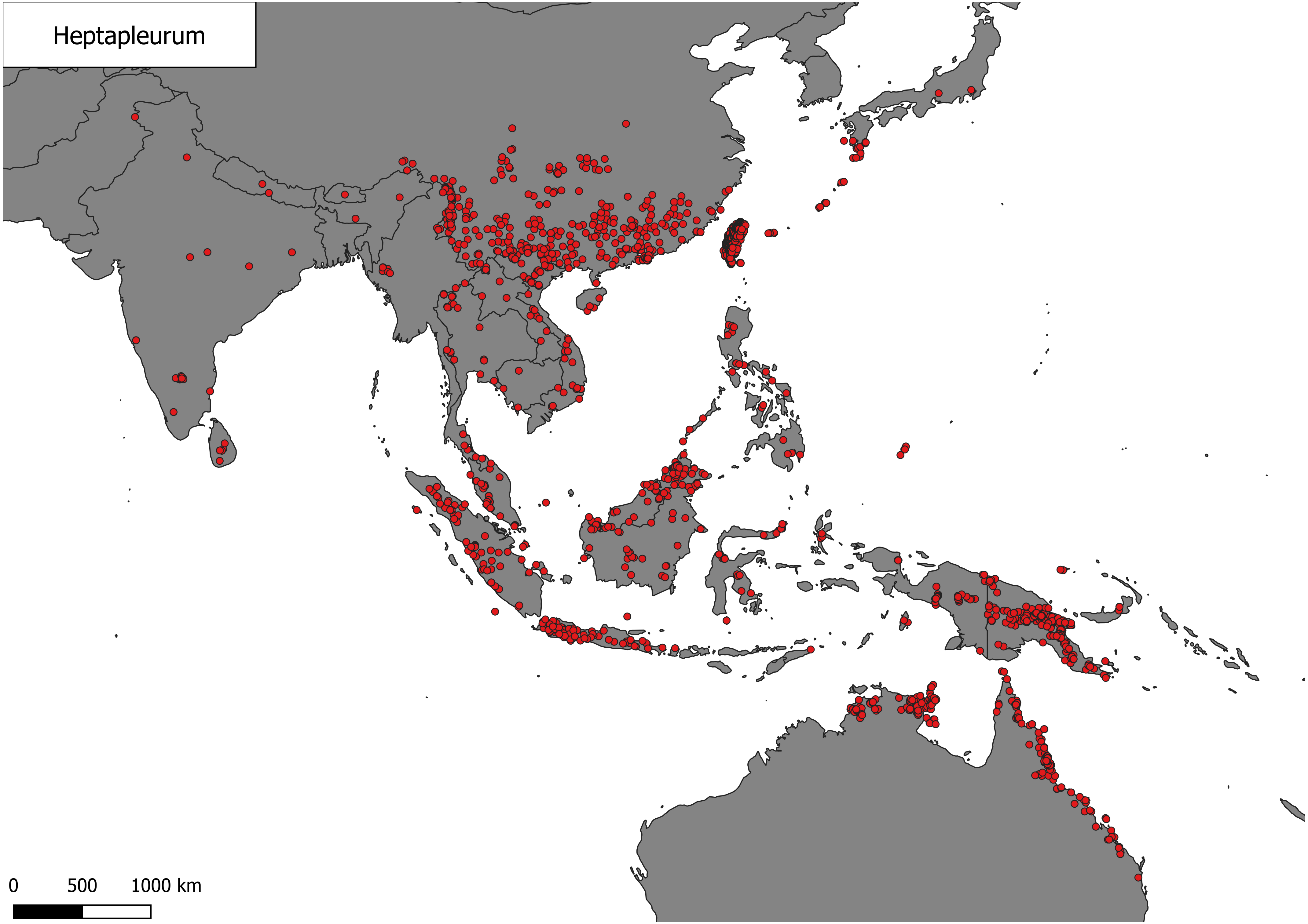
Heptapleurum
0
500
1000 km

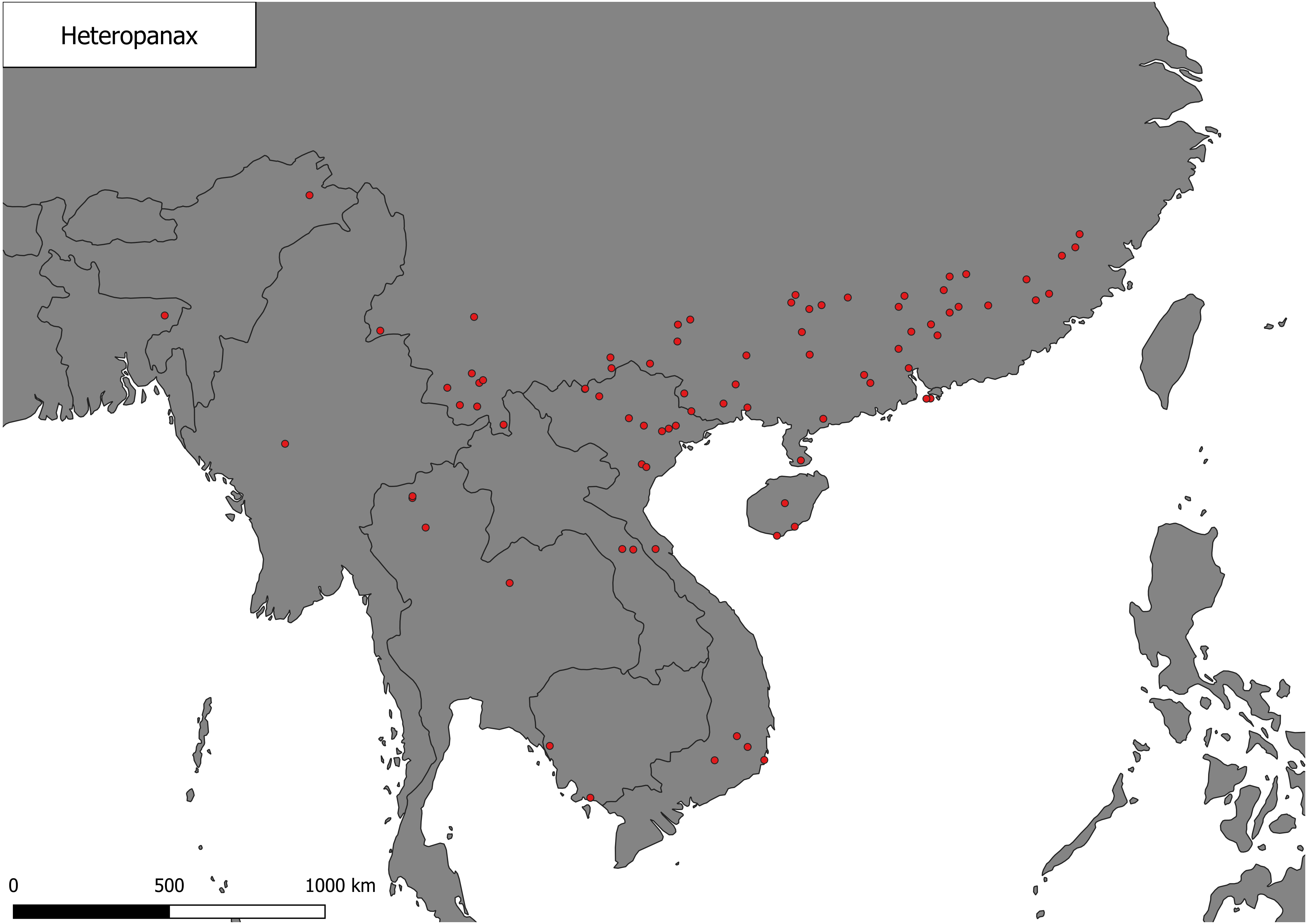

Heteropanax
0
500
1000 km

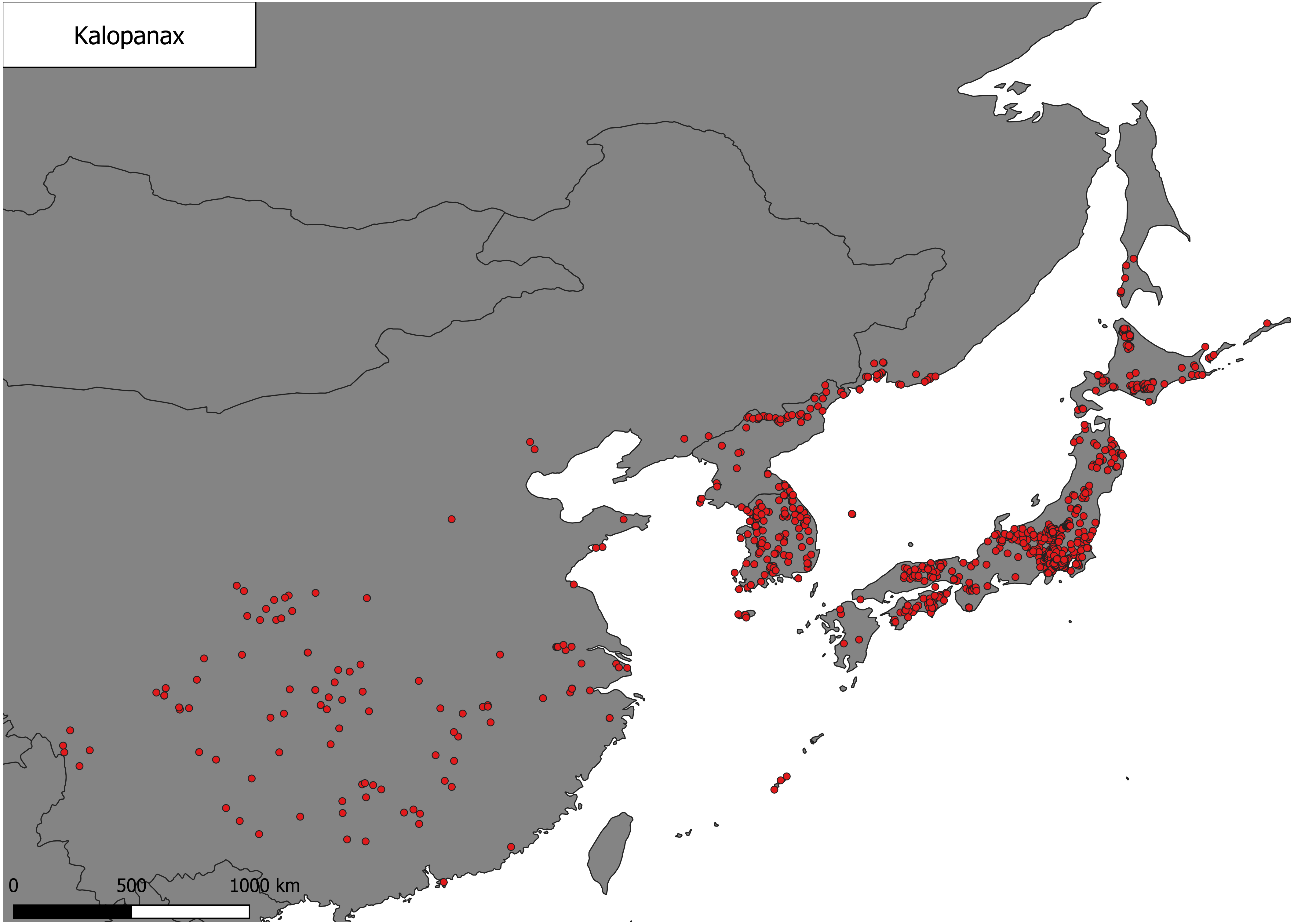
Kalopanax
0
500
1000 km

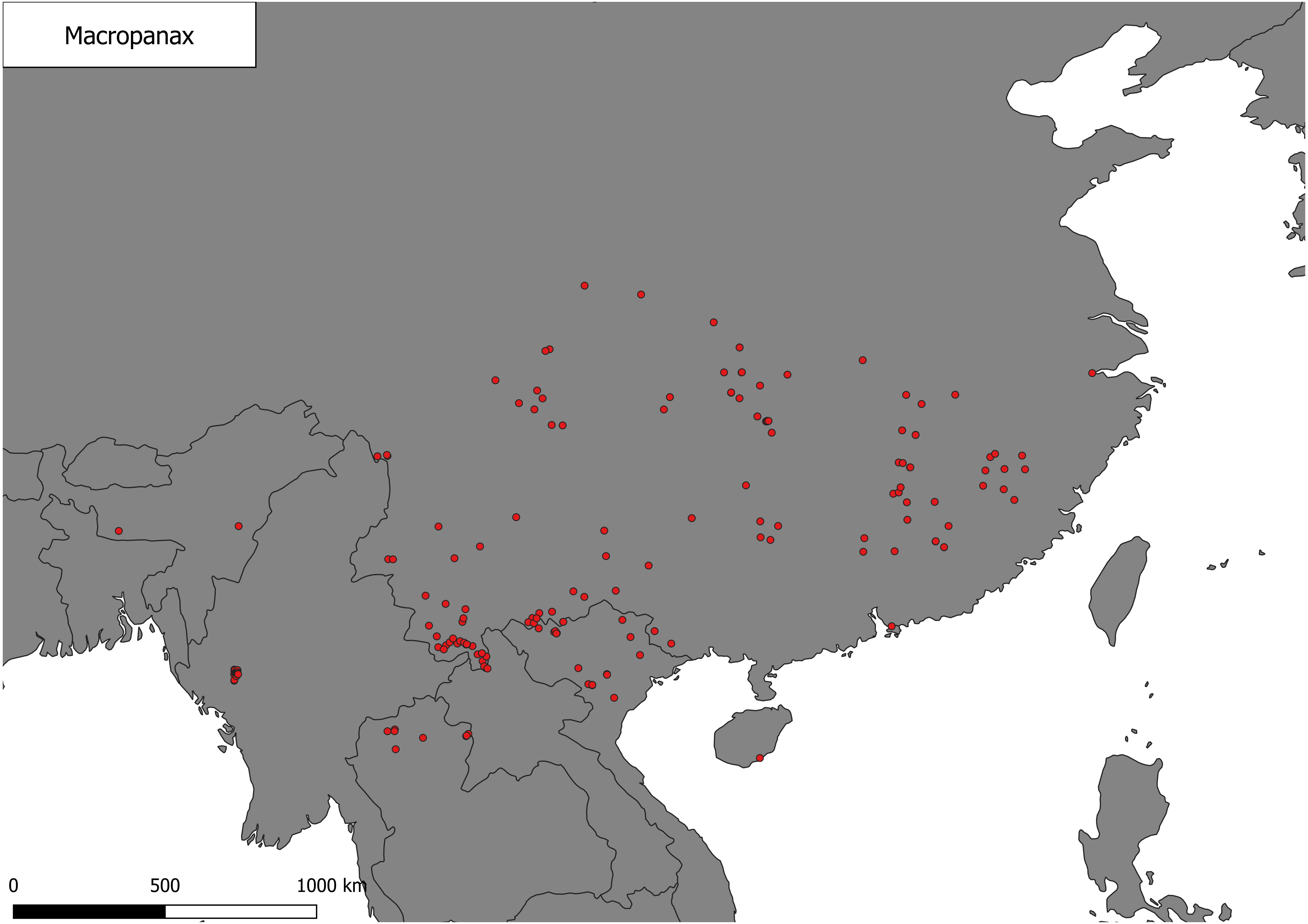

Macropanax
0
500
1000 km

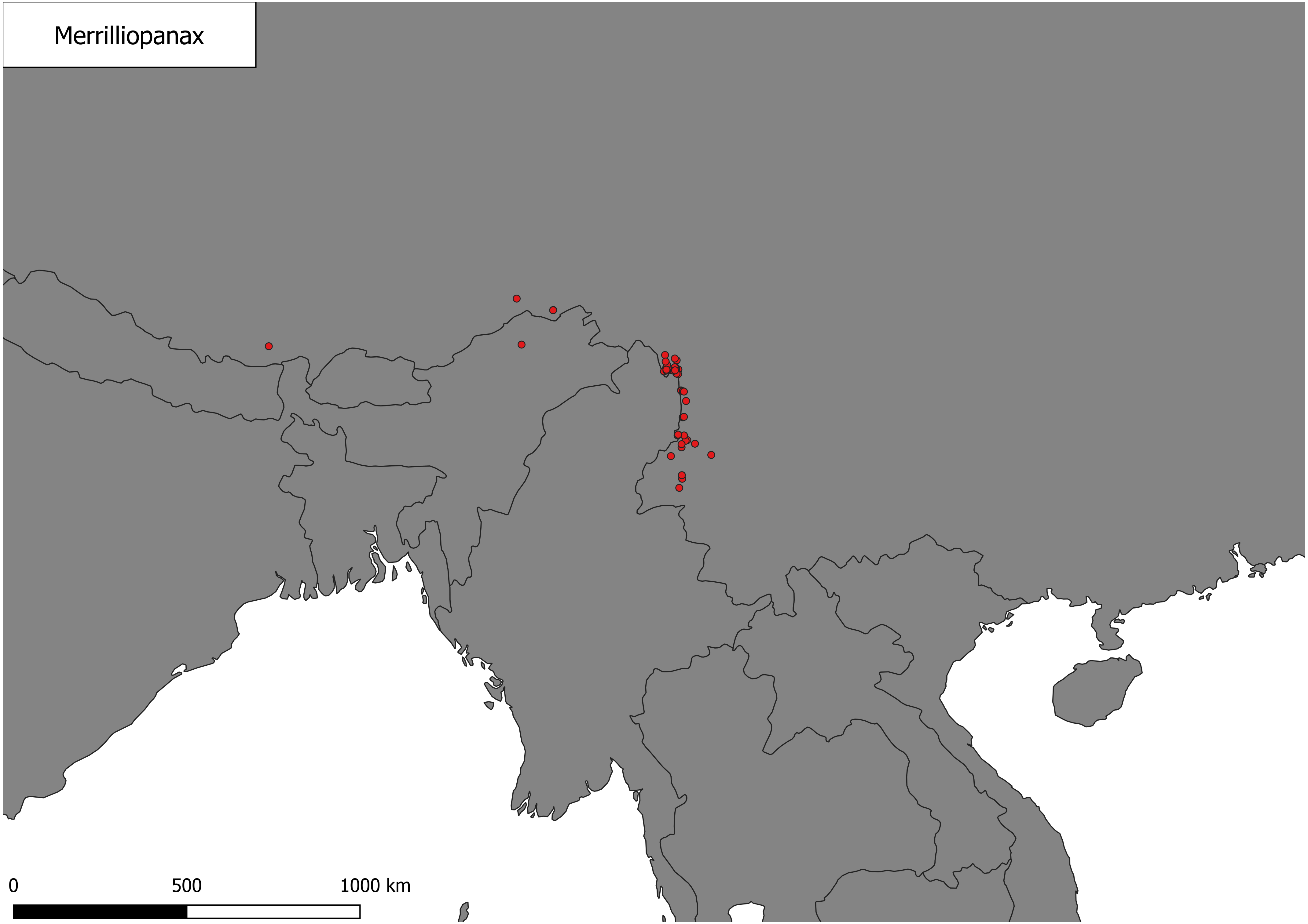
Merrilliopanax
0
500
1000 km

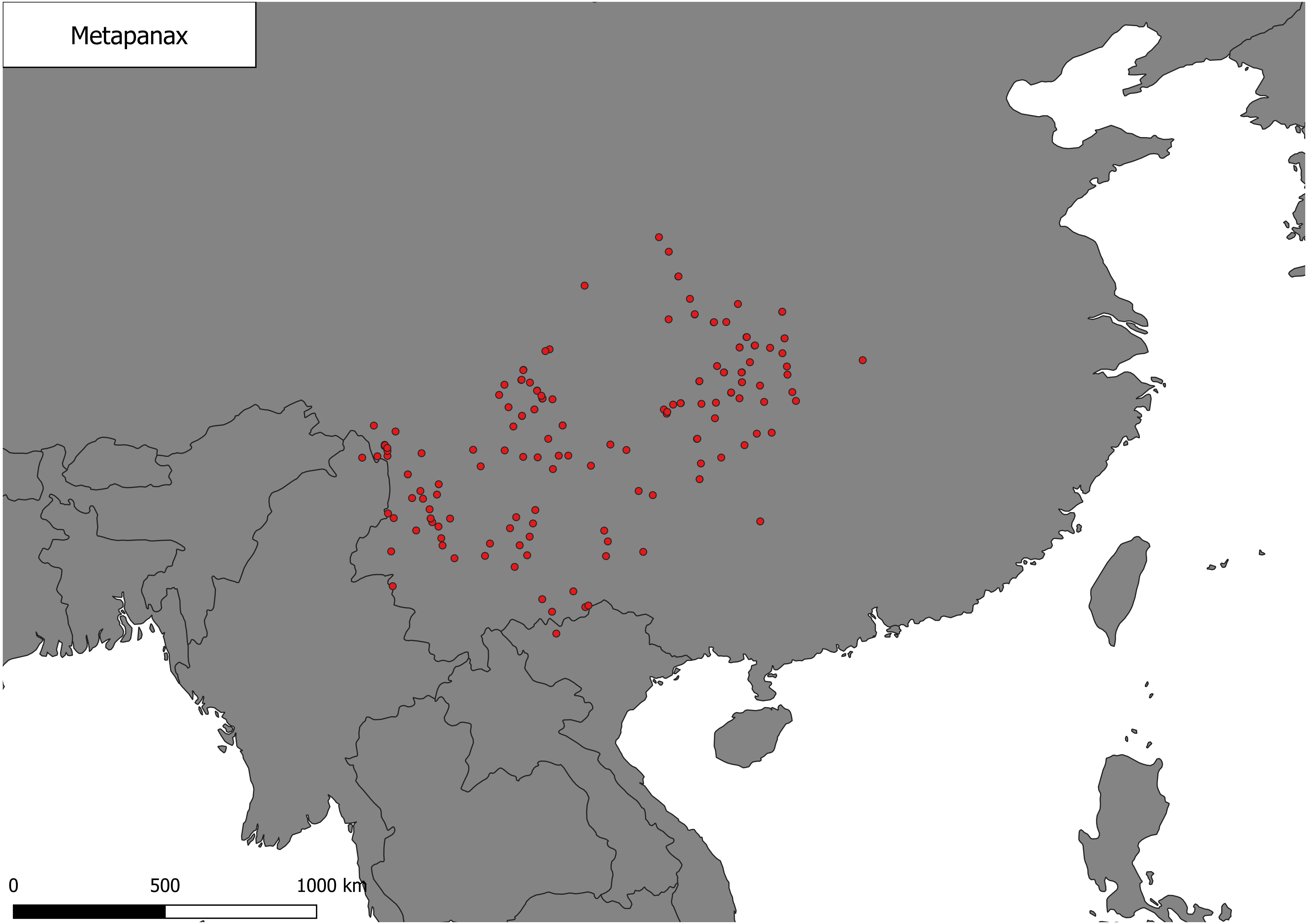
Metapanax
0
500
1000 km

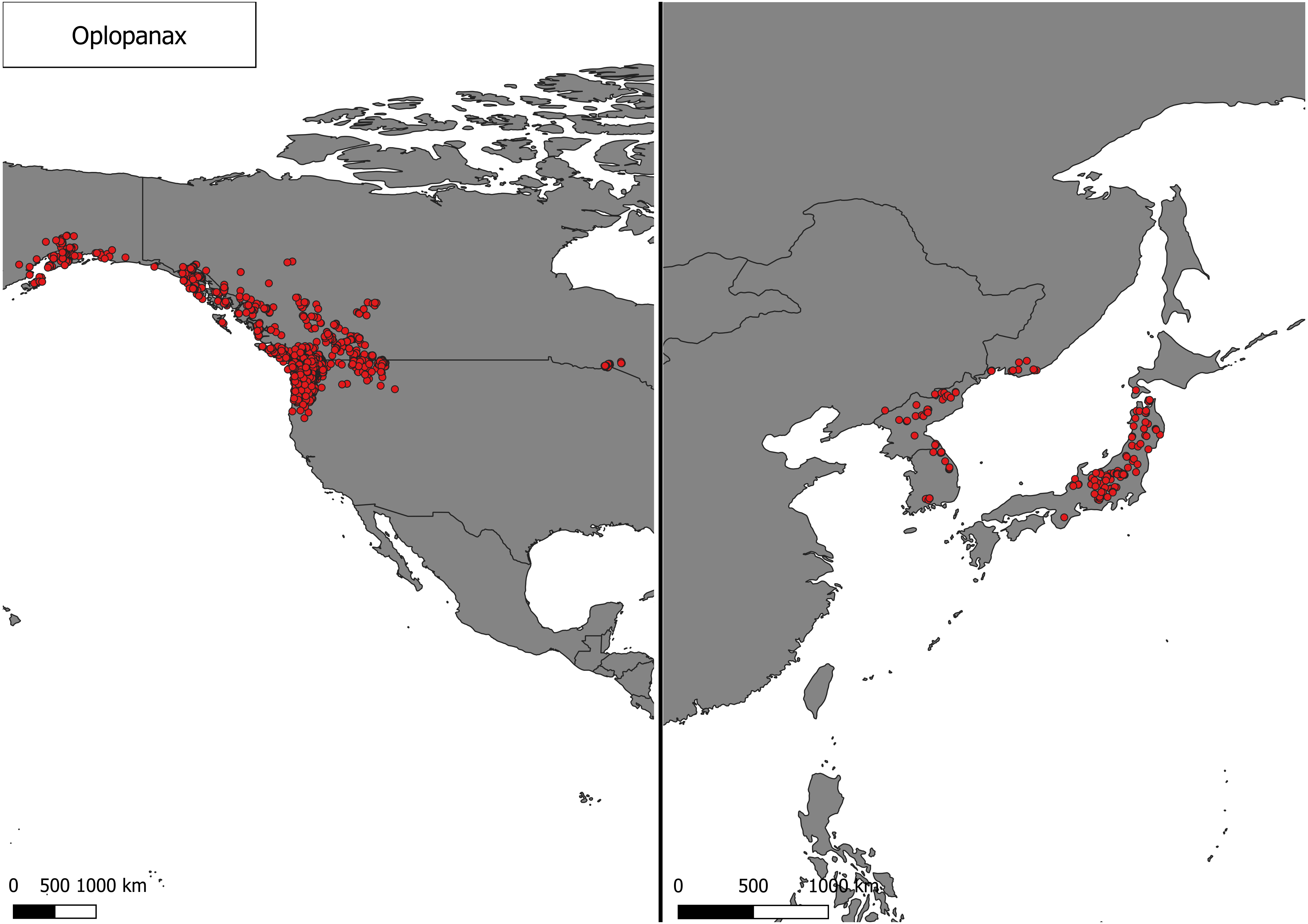
Oplopanax
0
500 1000 km
0
500
1000 km

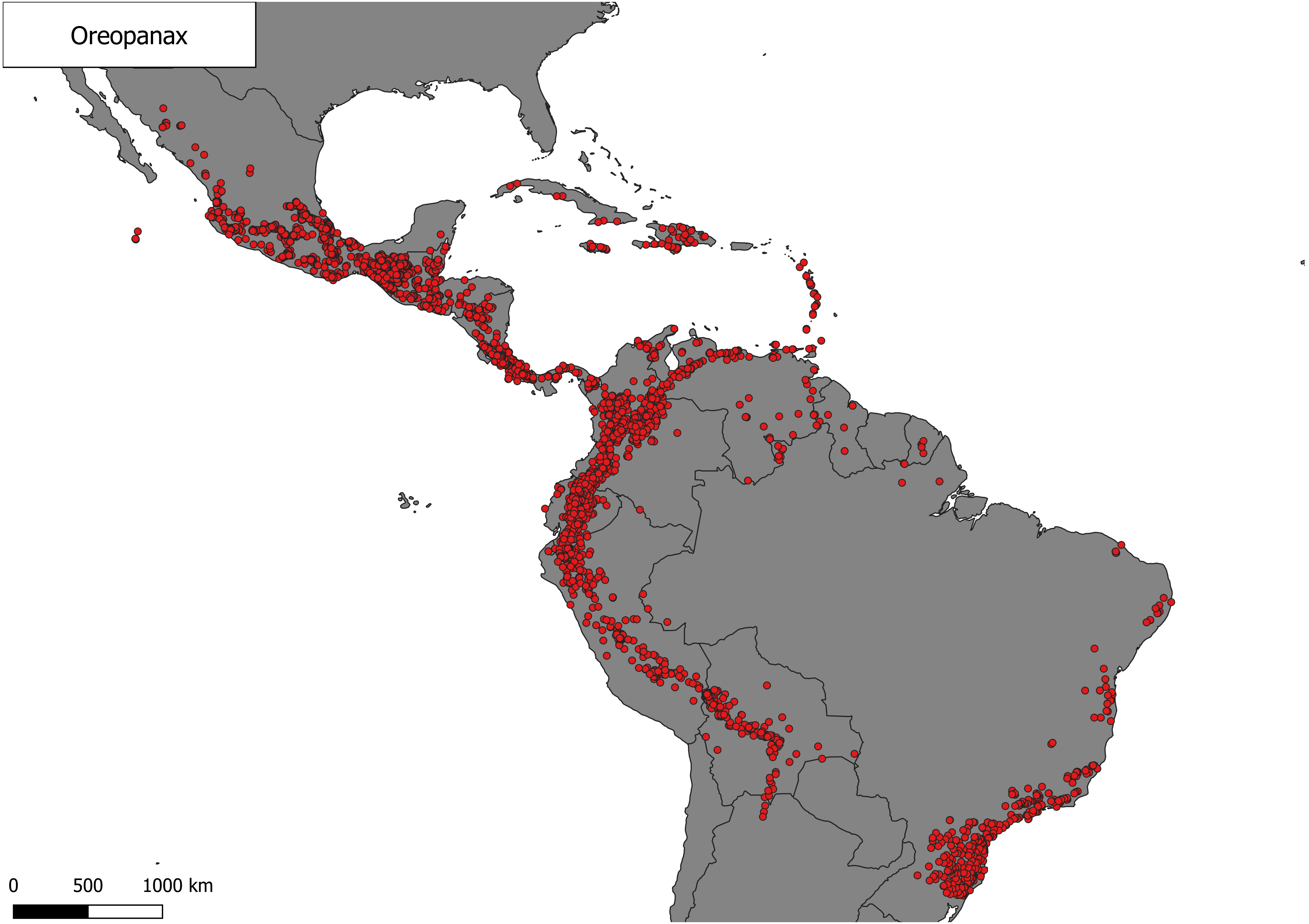

Oreopanax
0
500
1000 km

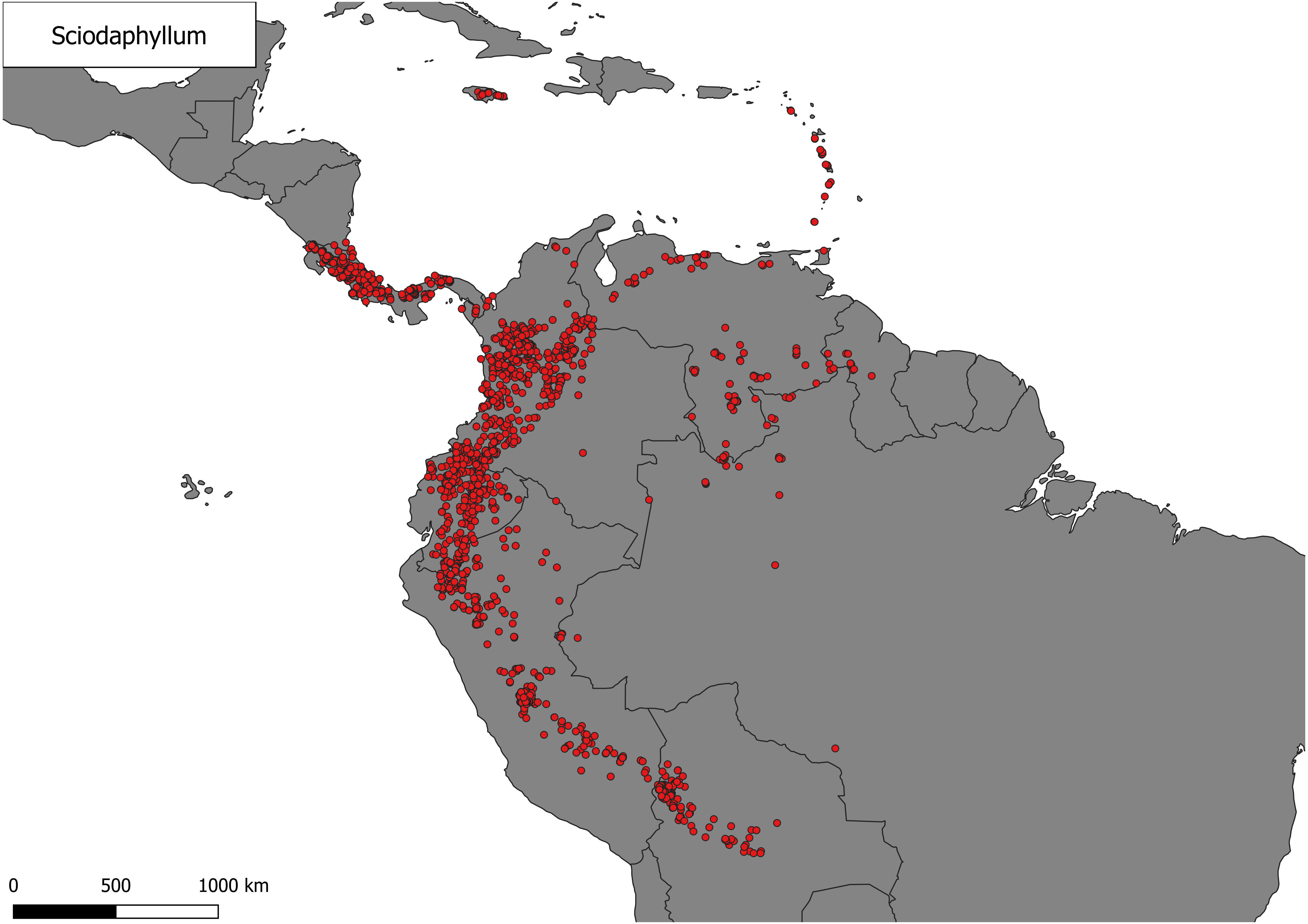
Sciodaphyllum
0
500
1000 km

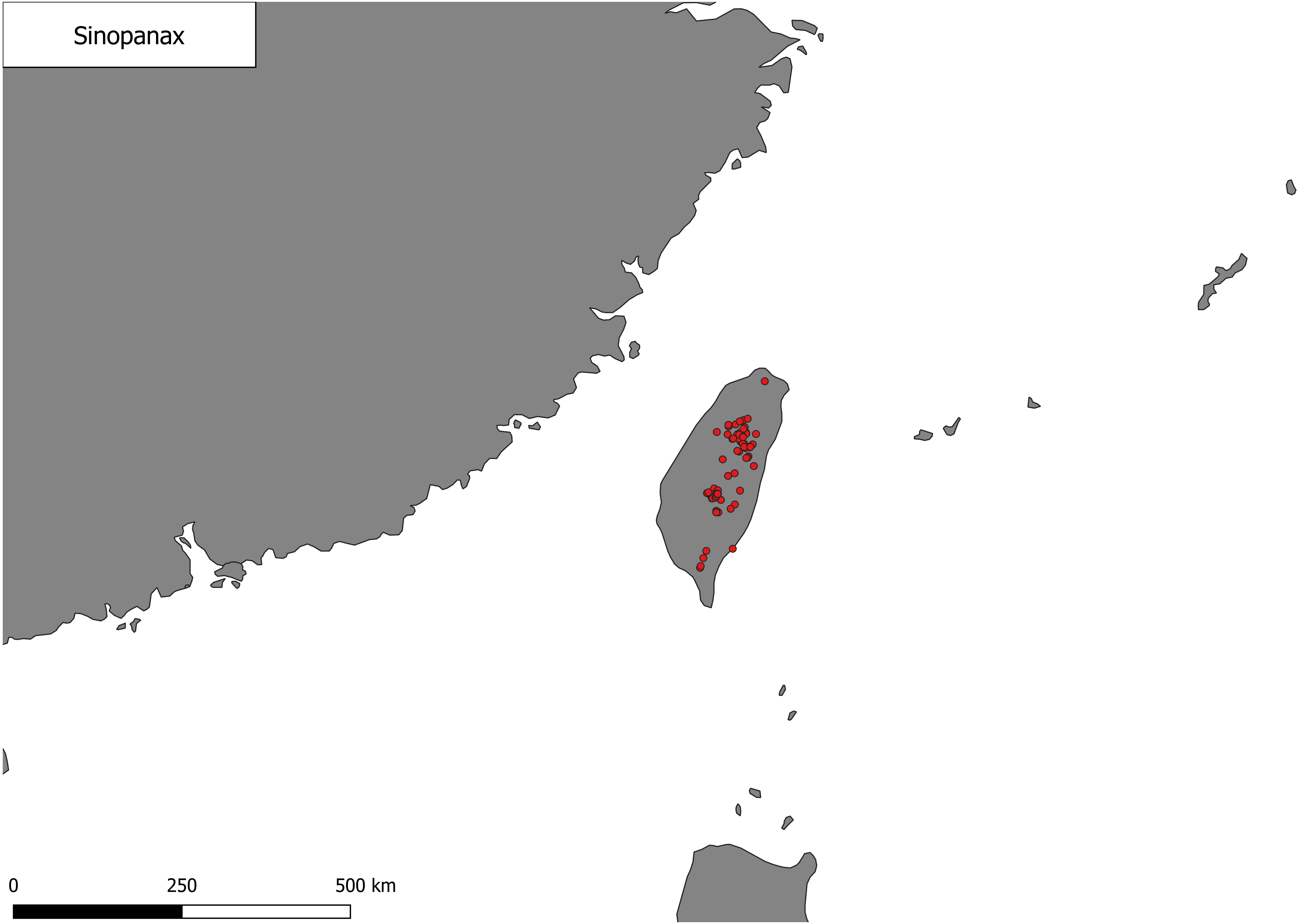
Sinopanax
0
250
500 km

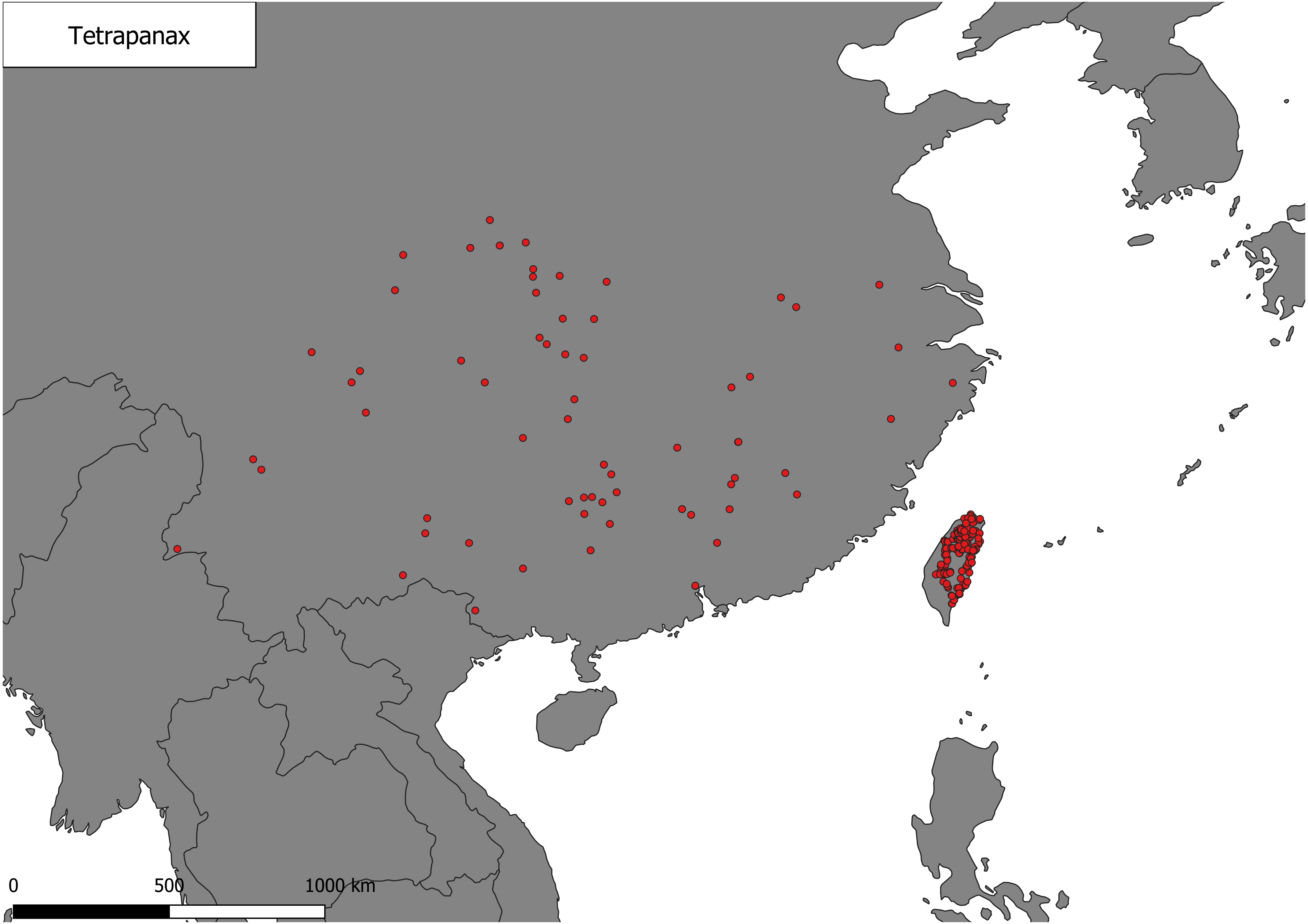
Tetrapanax
0
500
1000 km

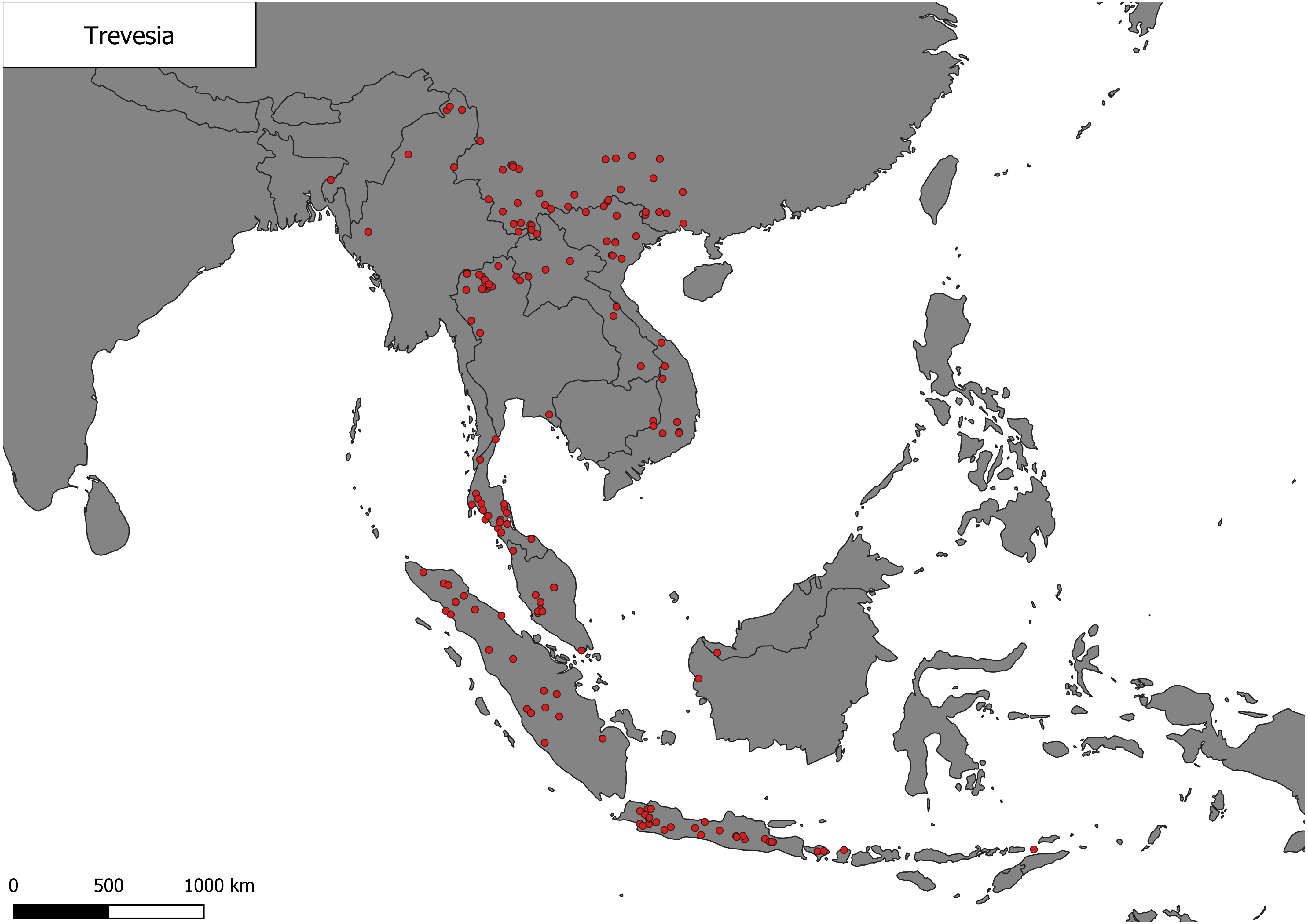
Trevesia
0
500
1000 km
