## Supplementary material for "Tropical-temperate dichotomy falls apart in the Asian Palmate Group of Araliaceae": World regionalizations

Appendix S3.1. Latitudinal classification

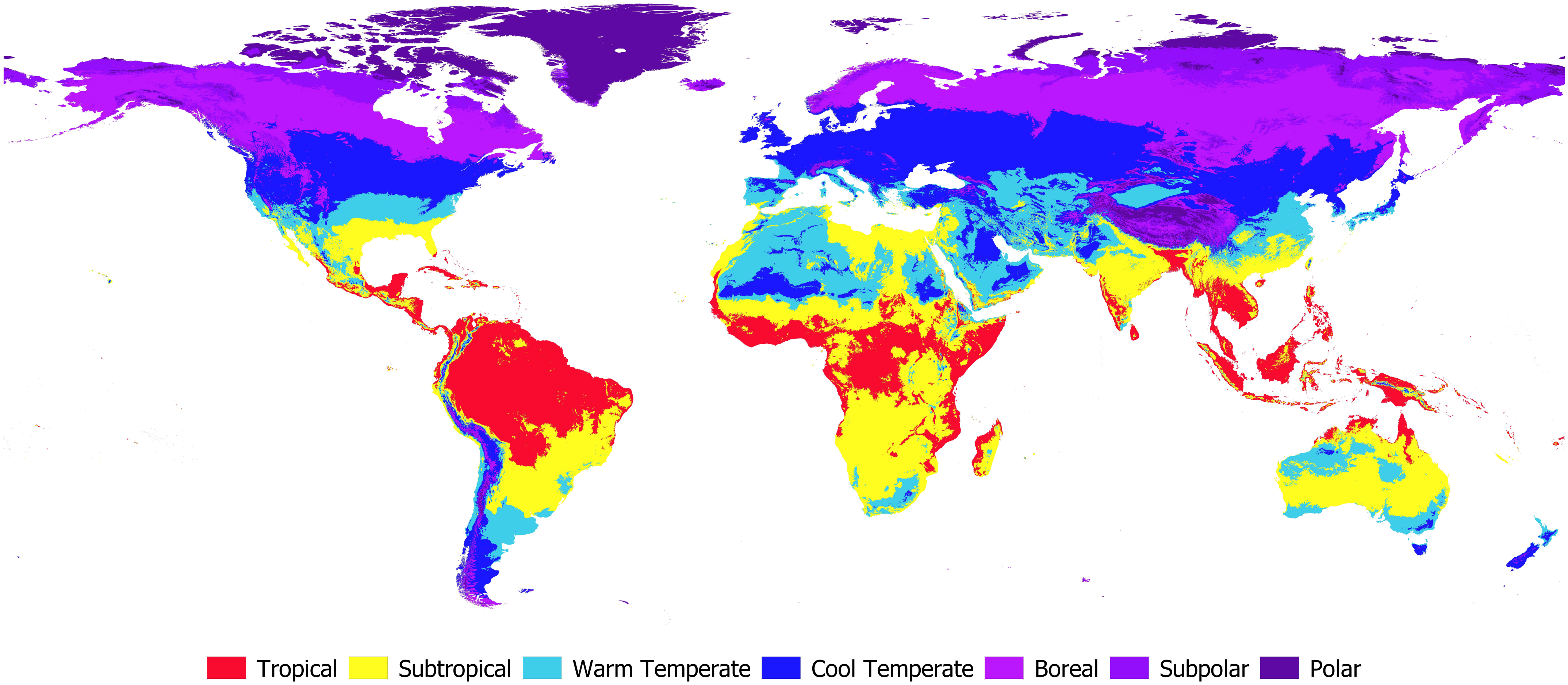

Appendix S3.2. Köppen's classification (Köppen & Geiger, 1936)

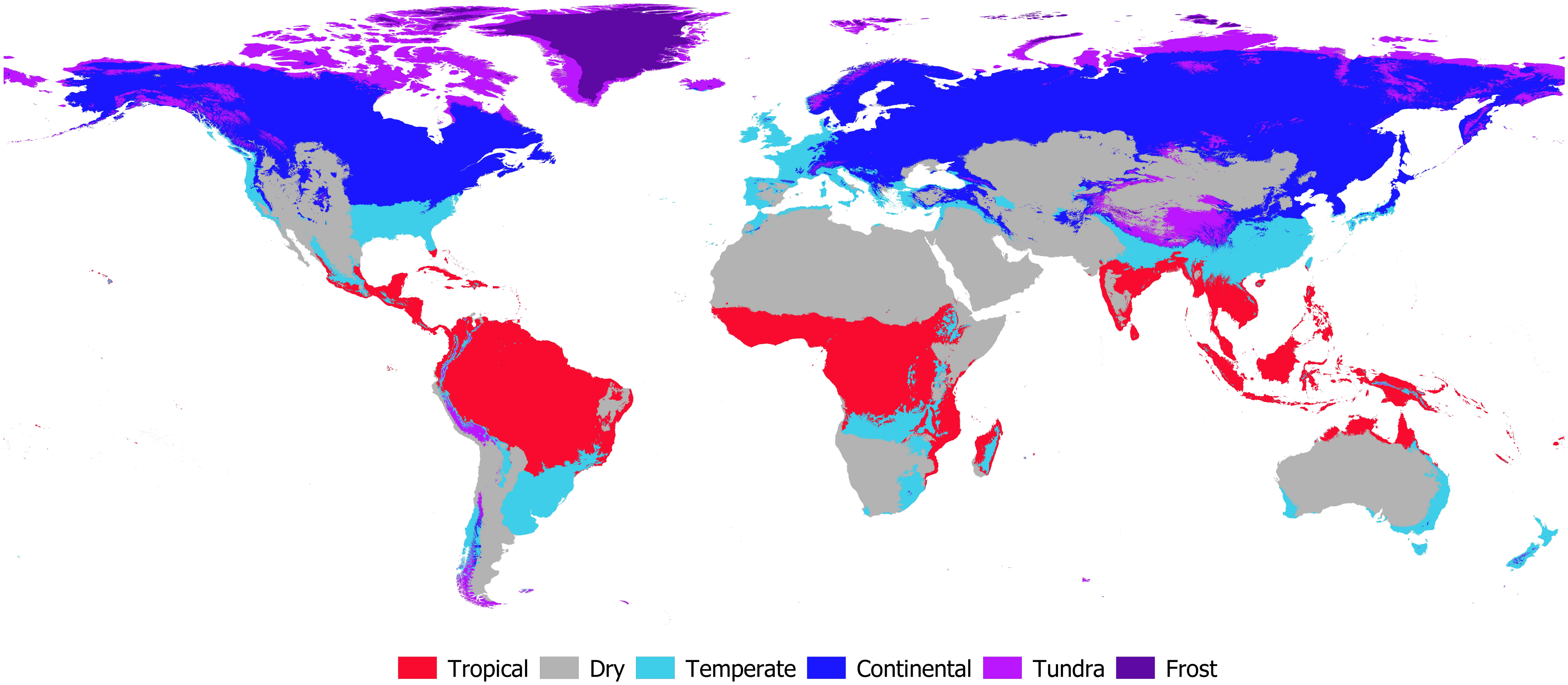

Appendix S3.3. Holdridge's classification (Holdridge, 1997)

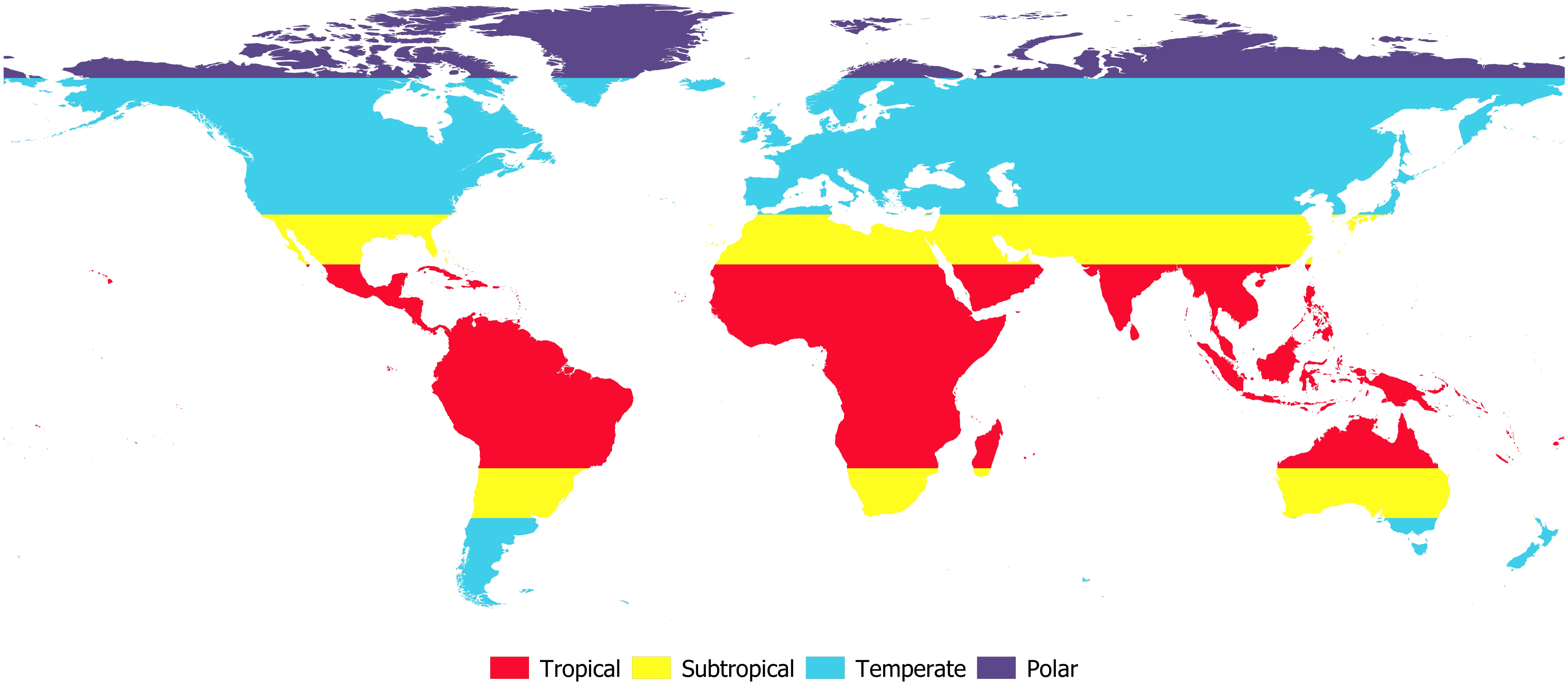

Appendix S3.4. Metzger's classification (Metzger et al., 2012)

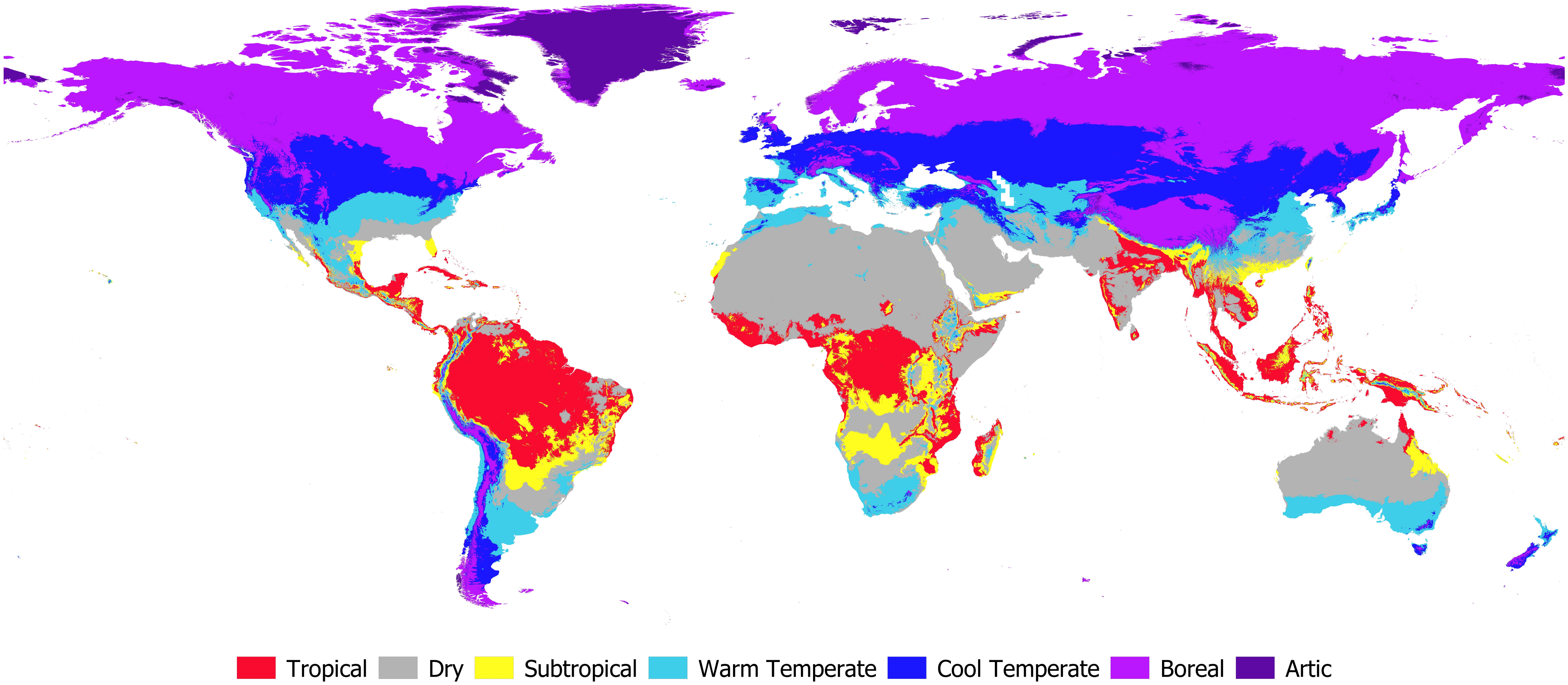

Appendix S3.5. Ecoregions simplified classification (Dinerstein et al., 2017)

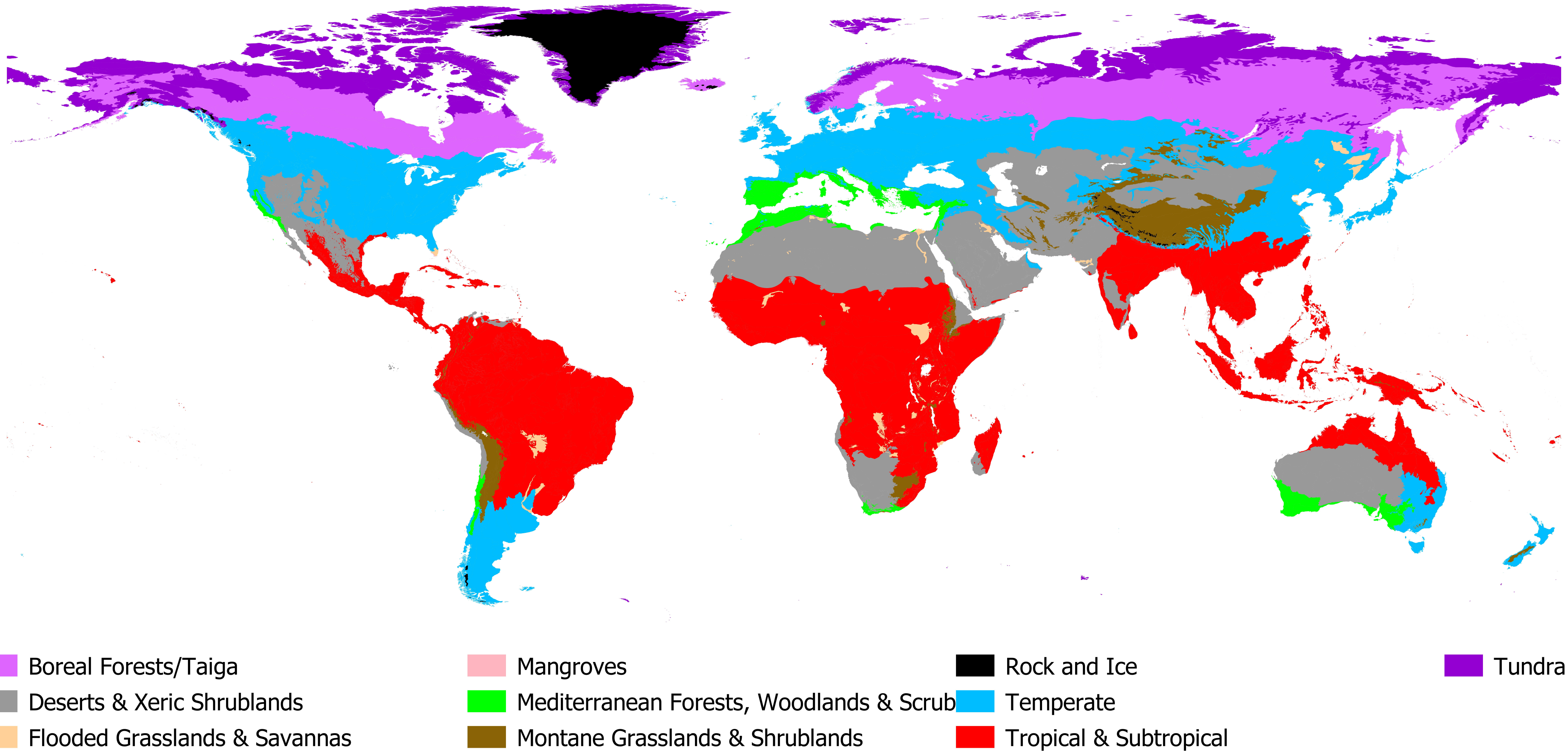
