## Supplementary material for "Tropical-temperate dichotomy falls apart in the Asian Palmate Group of Araliaceae": Heat maps of Asian Palmate Group genera

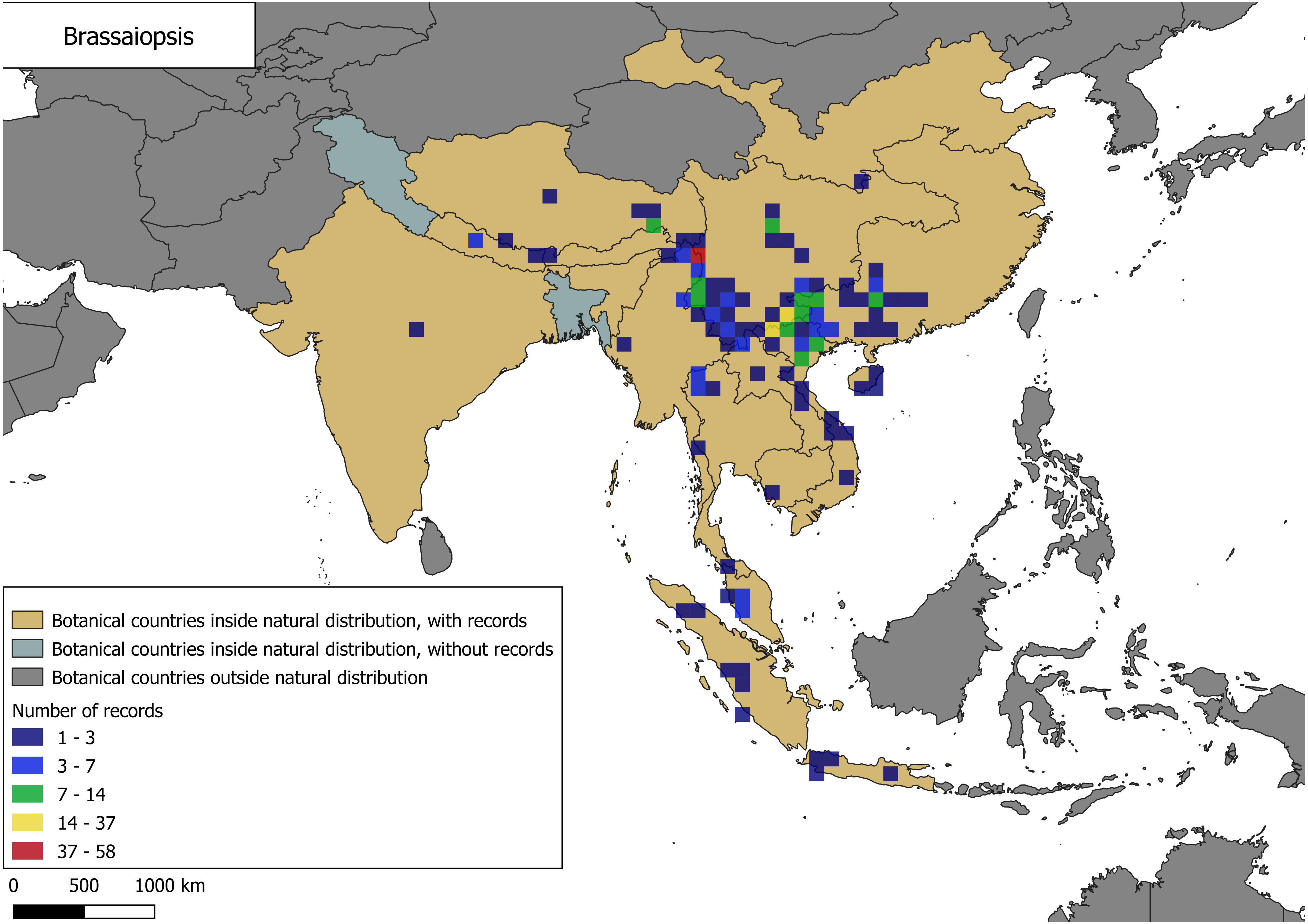
Brassaiopsis
Botanical countries inside natural distribution, with records
Botanical countries inside natural distribution, without records
Botanical countries outside natural distribution
Number of records
1 - 3
3 - 7
7 - 14
14 - 37
37 - 58
0
500
1000 km

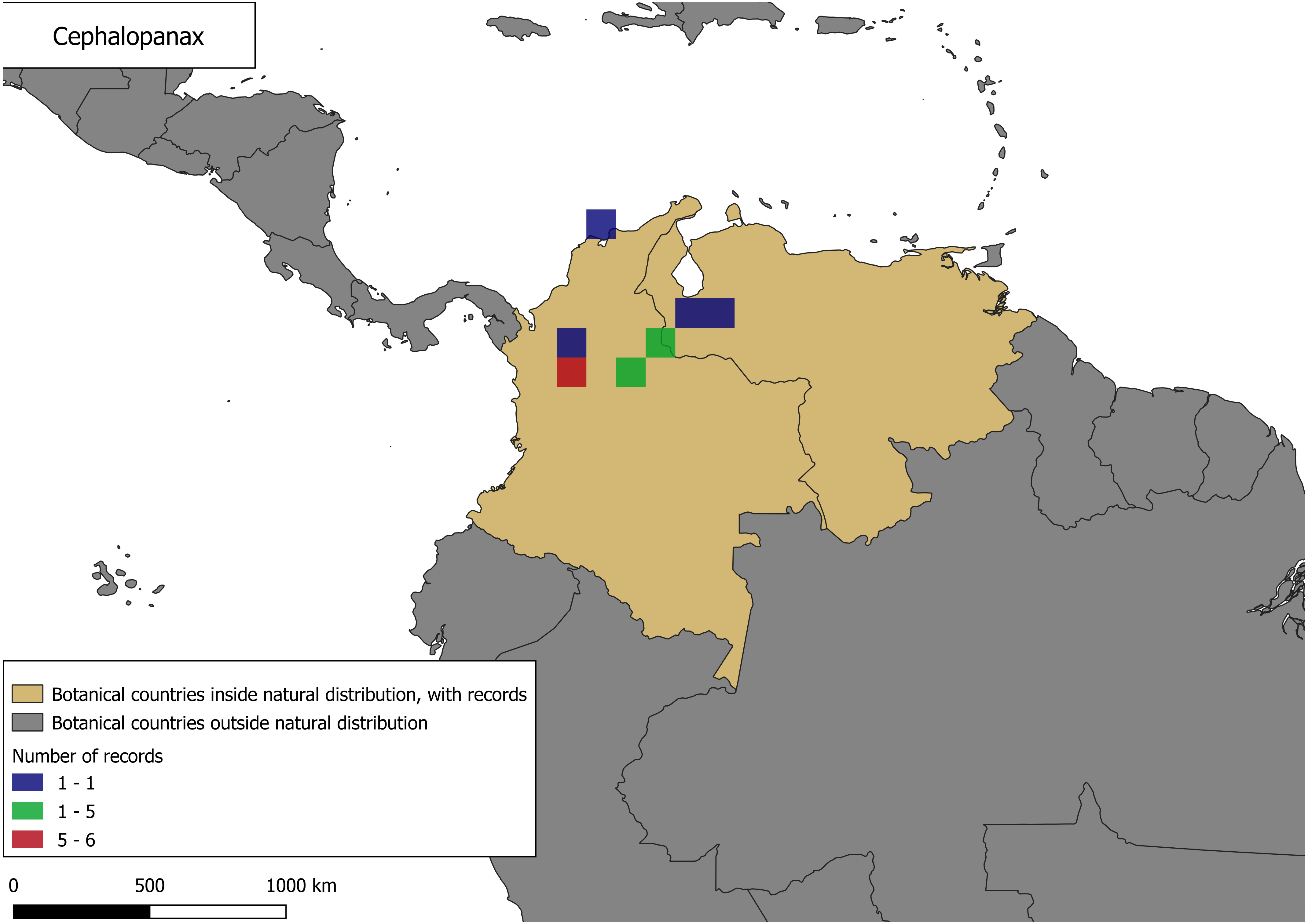
Cephalopanax
Botanical countries inside natural distribution, with records
Botanical countries outside natural distribution
Number of records
1 - 1
1 - 5
5 - 6
0
500
1000 km

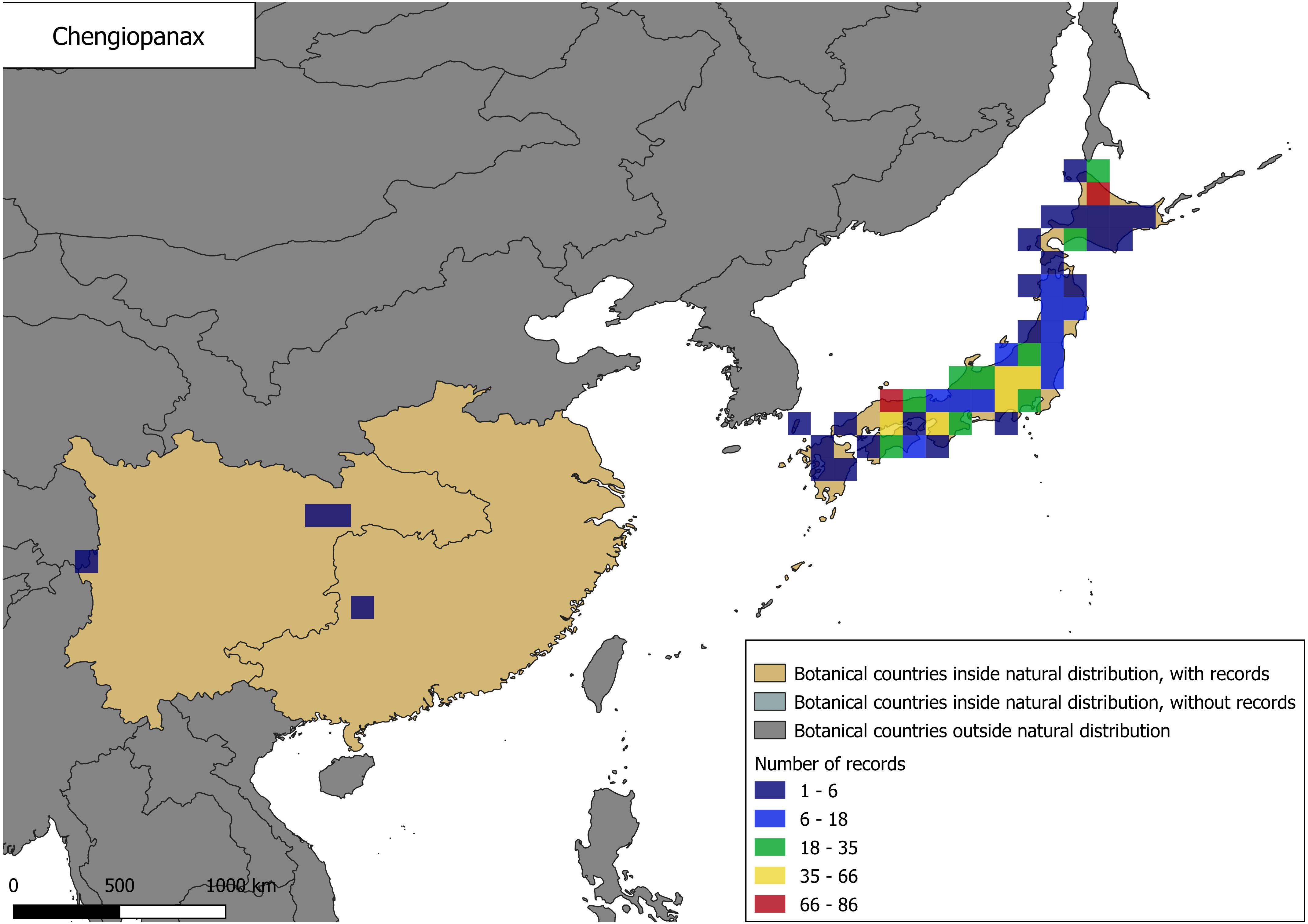
Chengiopanax
Botanical countries inside natural distribution, with records
Botanical countries inside natural distribution, without records
Botanical countries outside natural distribution
Number of records
1 - 6
6 - 18
18 - 35
35 - 66
66 - 86
0
500
1000 km

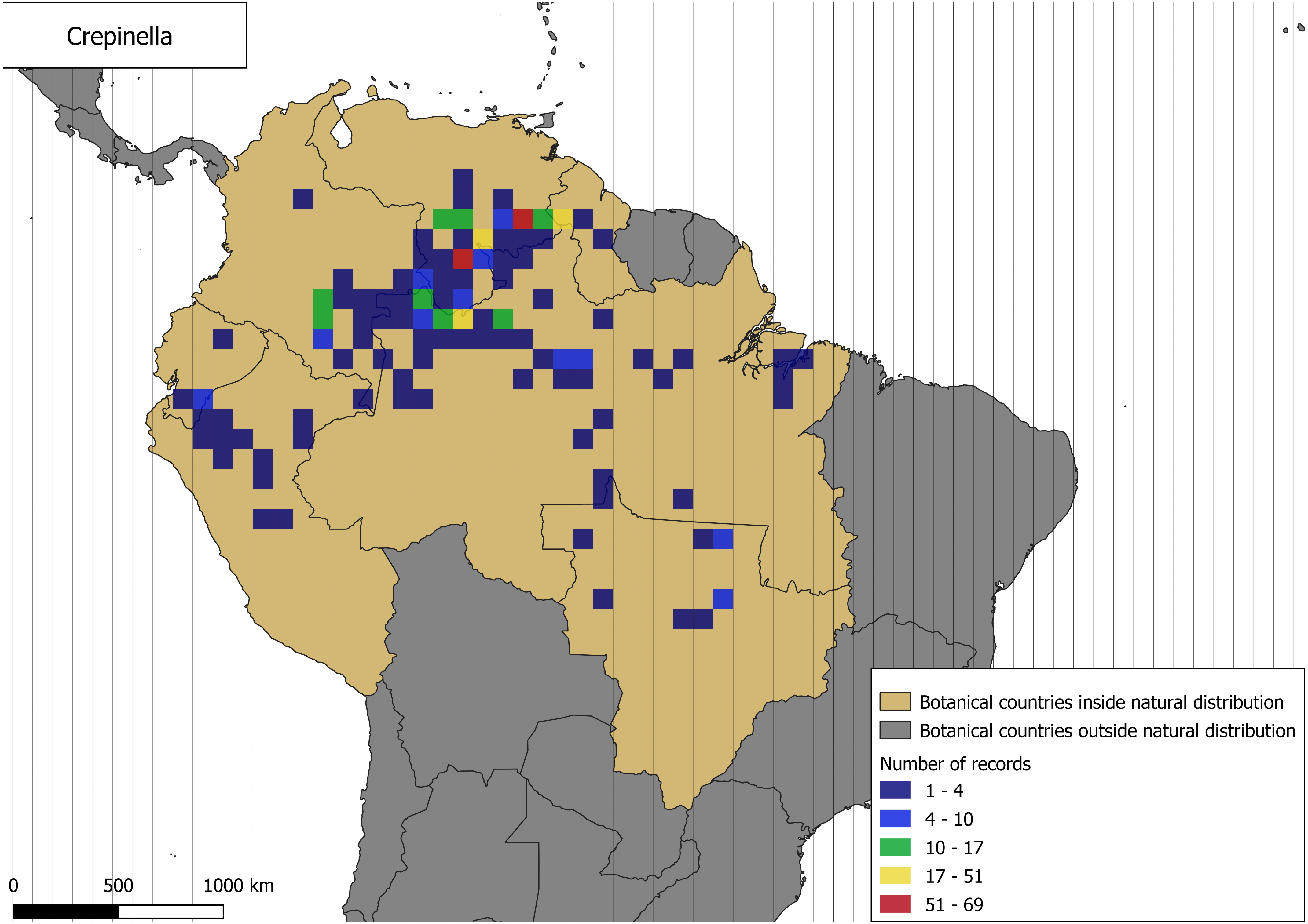
Crepinella
Botanical countries inside natural distribution
Botanical countries outside natural distribution
Number of records
1 - 4
4 - 10
10 - 17
17 - 51
51 - 69
0
500
1000 km

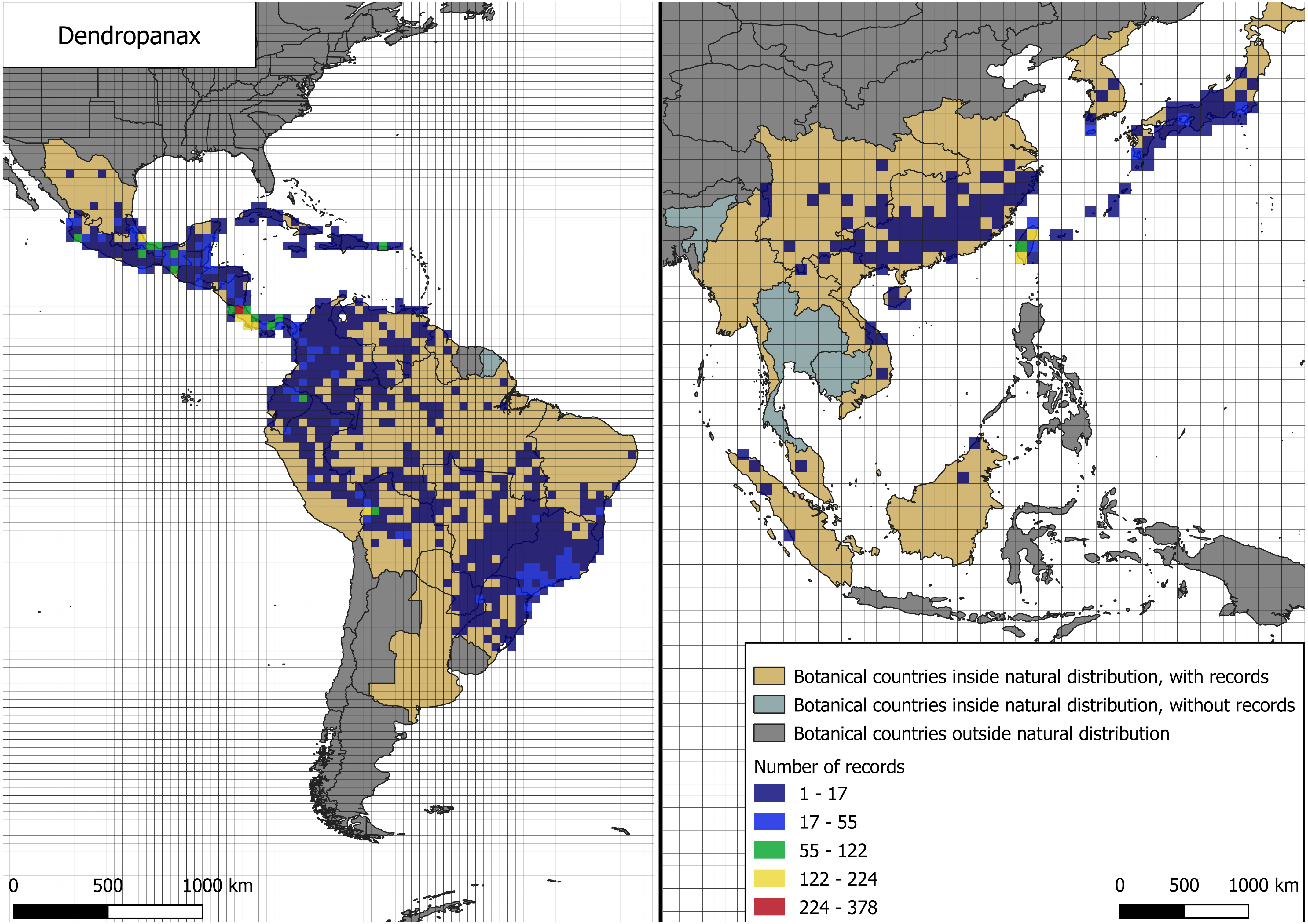

Dendropanax
Botanical countries inside natural distribution, with records
Botanical countries inside natural distribution, without records
Botanical countries outside natural distribution
Number of records
1 - 17
17 - 55
55 - 122
122 - 224
224 - 378
0 500 1000 km
0 500 1000 km

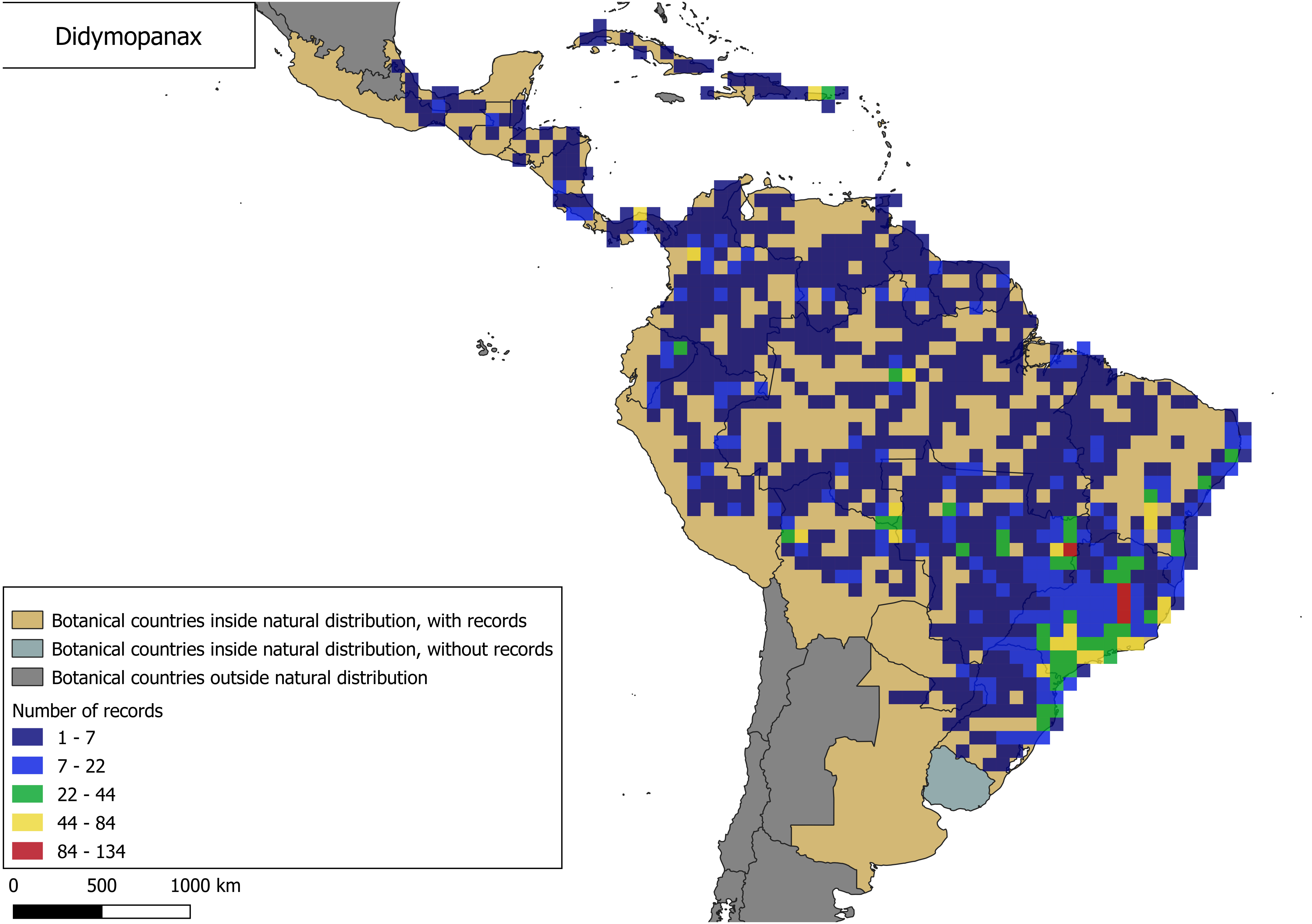

Didymopanax
Botanical countries inside natural distribution, with records
Botanical countries inside natural distribution, without records
Botanical countries outside natural distribution
Number of records
1 - 7
7 - 22
22 - 44
44 - 84
84 - 134
0
500
1000 km

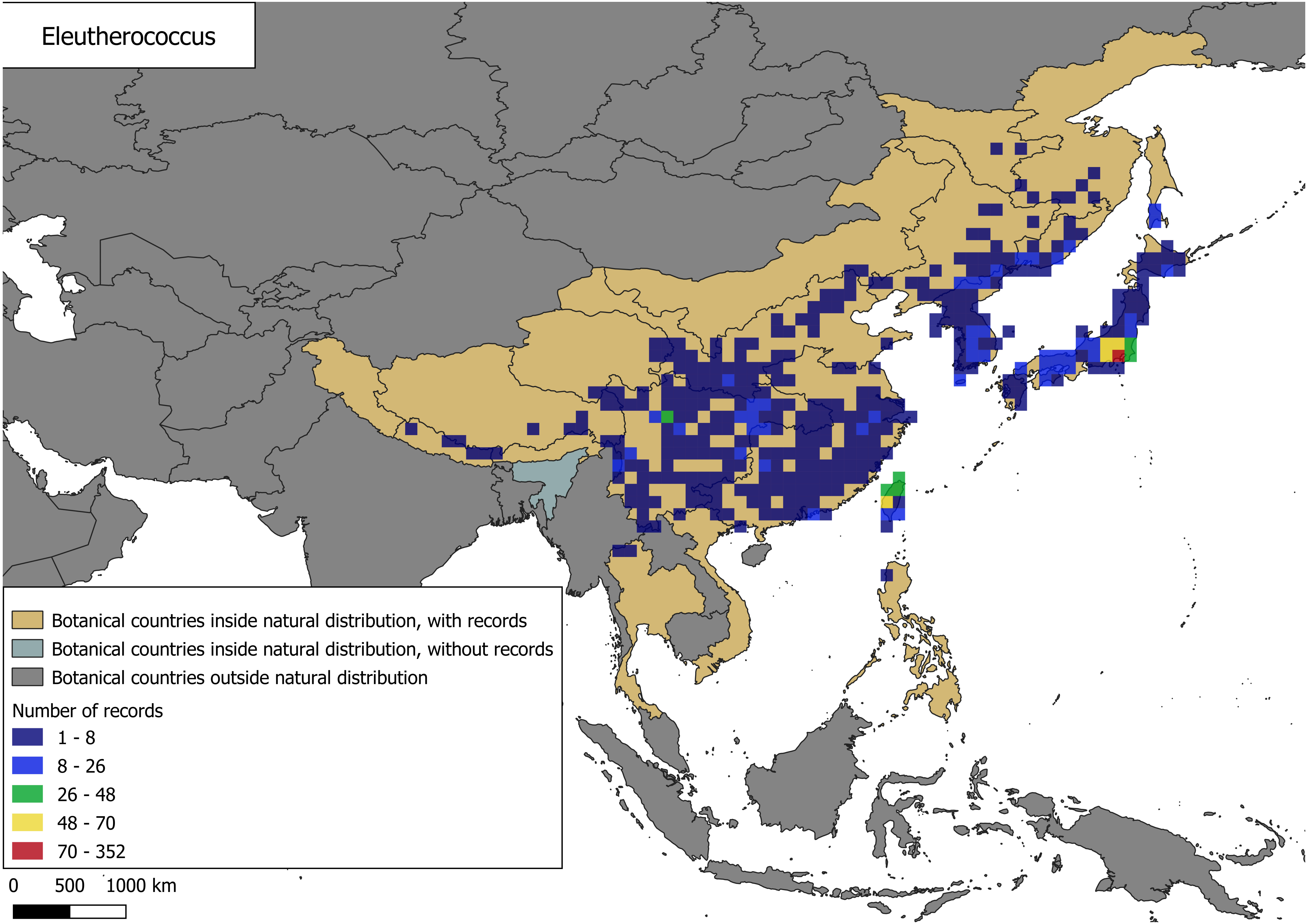
Eleutherococcus
Botanical countries inside natural distribution, with records
Botanical countries inside natural distribution, without records
Botanical countries outside natural distribution
Number of records
1 - 8
8 - 26
26 - 48
48 - 70
70 - 352
0
500
1000 km

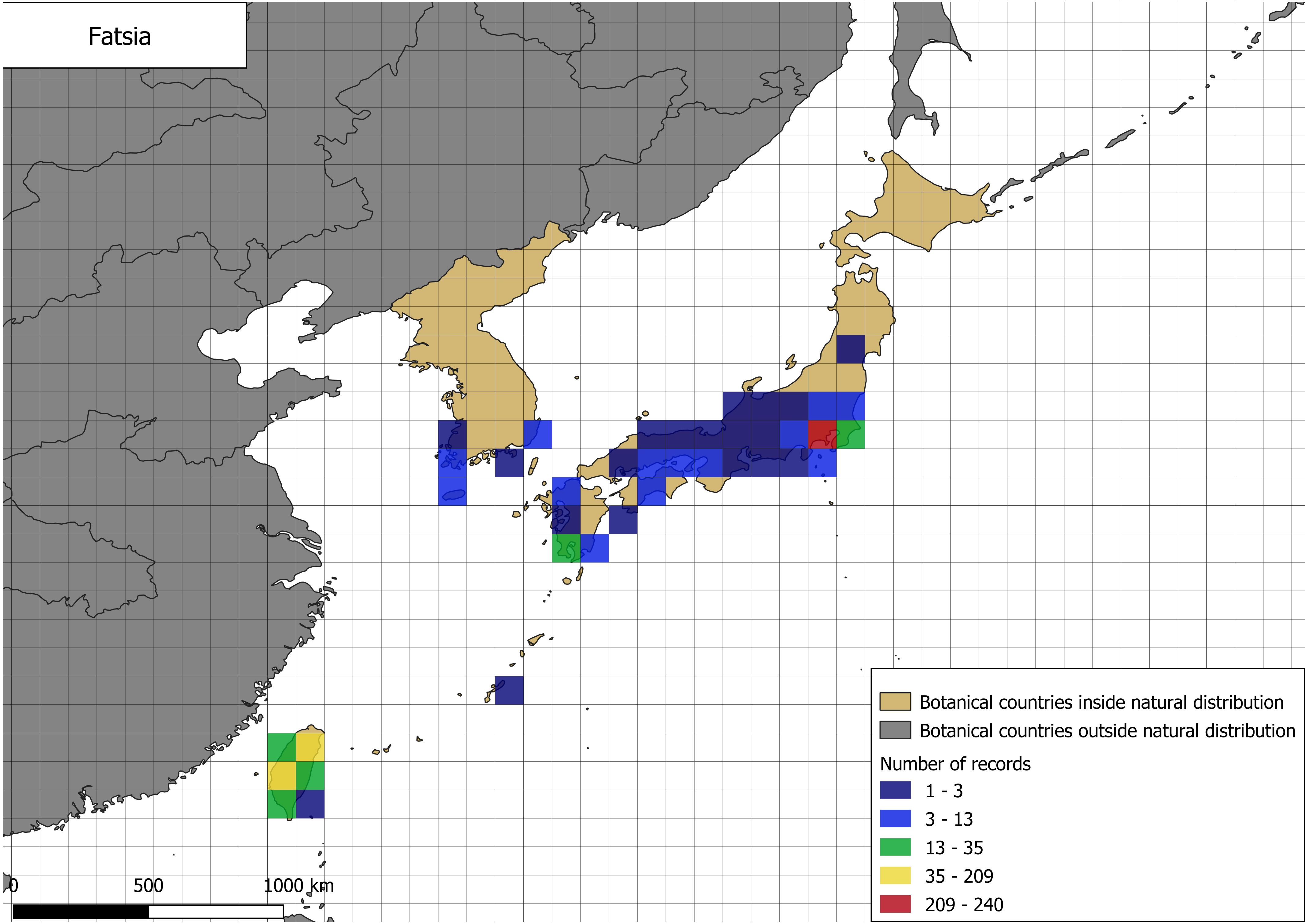
Fatsia
Botanical countries inside natural distribution
Botanical countries outside natural distribution
Number of records
1 - 3
3 - 13
13 - 35
35 - 209
209 - 240
0
500
1000 km

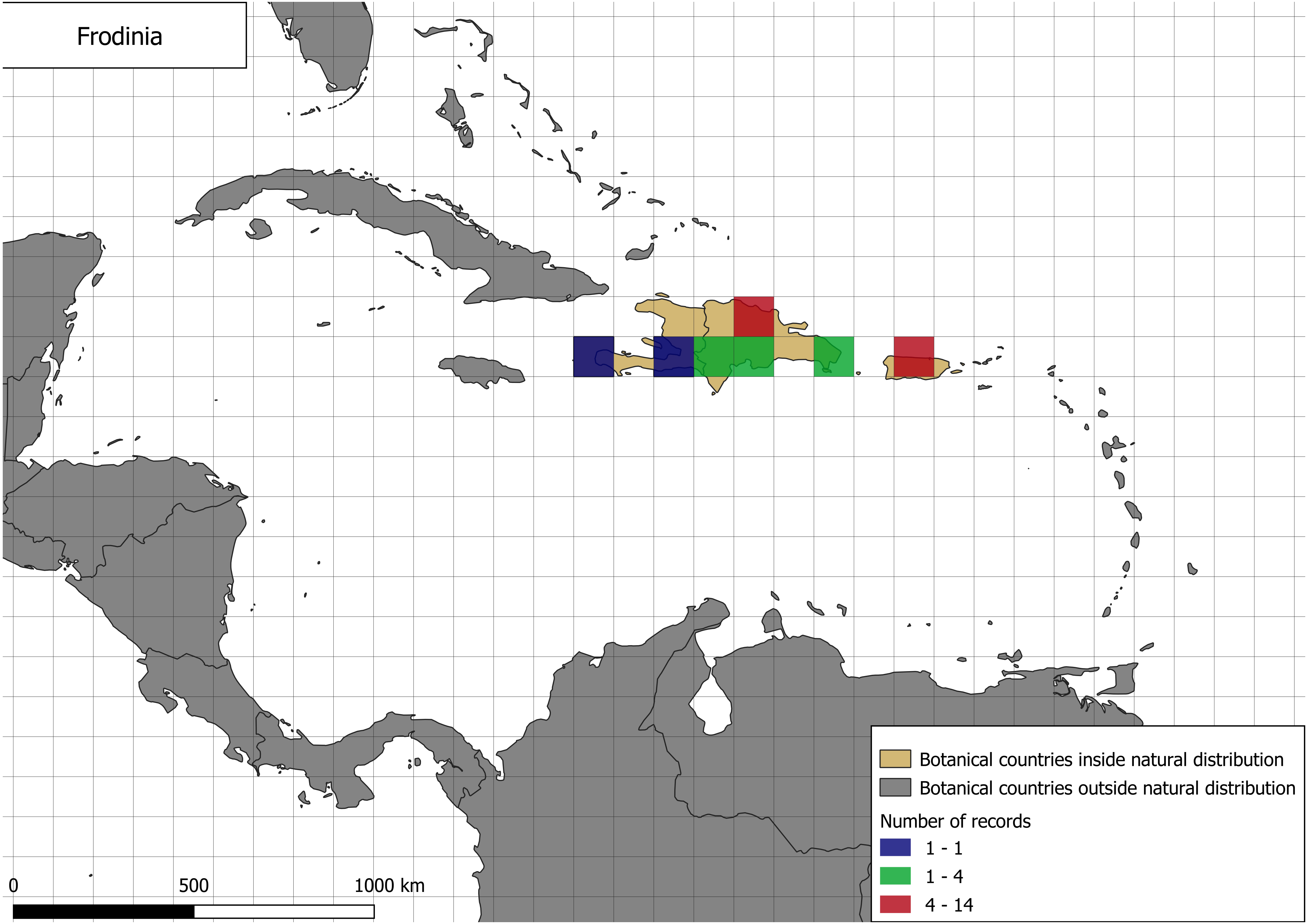
Frodinia
Botanical countries inside natural distribution
Botanical countries outside natural distribution
Number of records
1 - 1
1 - 4
4 - 14
0
500
1000 km

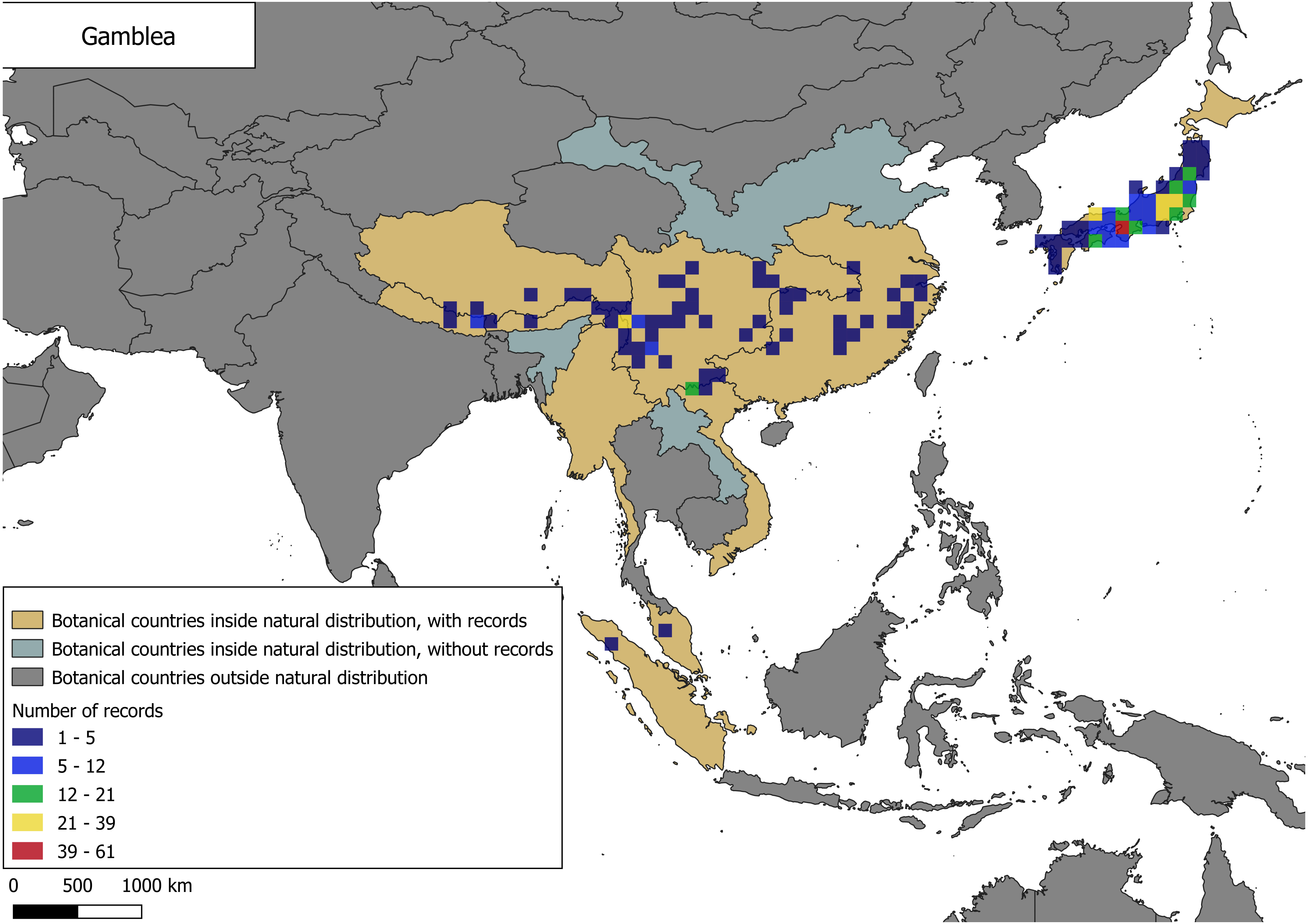
Gamblea
Botanical countries inside natural distribution, with records
Botanical countries inside natural distribution, without records
Botanical countries outside natural distribution
Number of records
1 - 5
5 - 12
12 - 21
21 - 39
39 - 61
0
500
1000 km

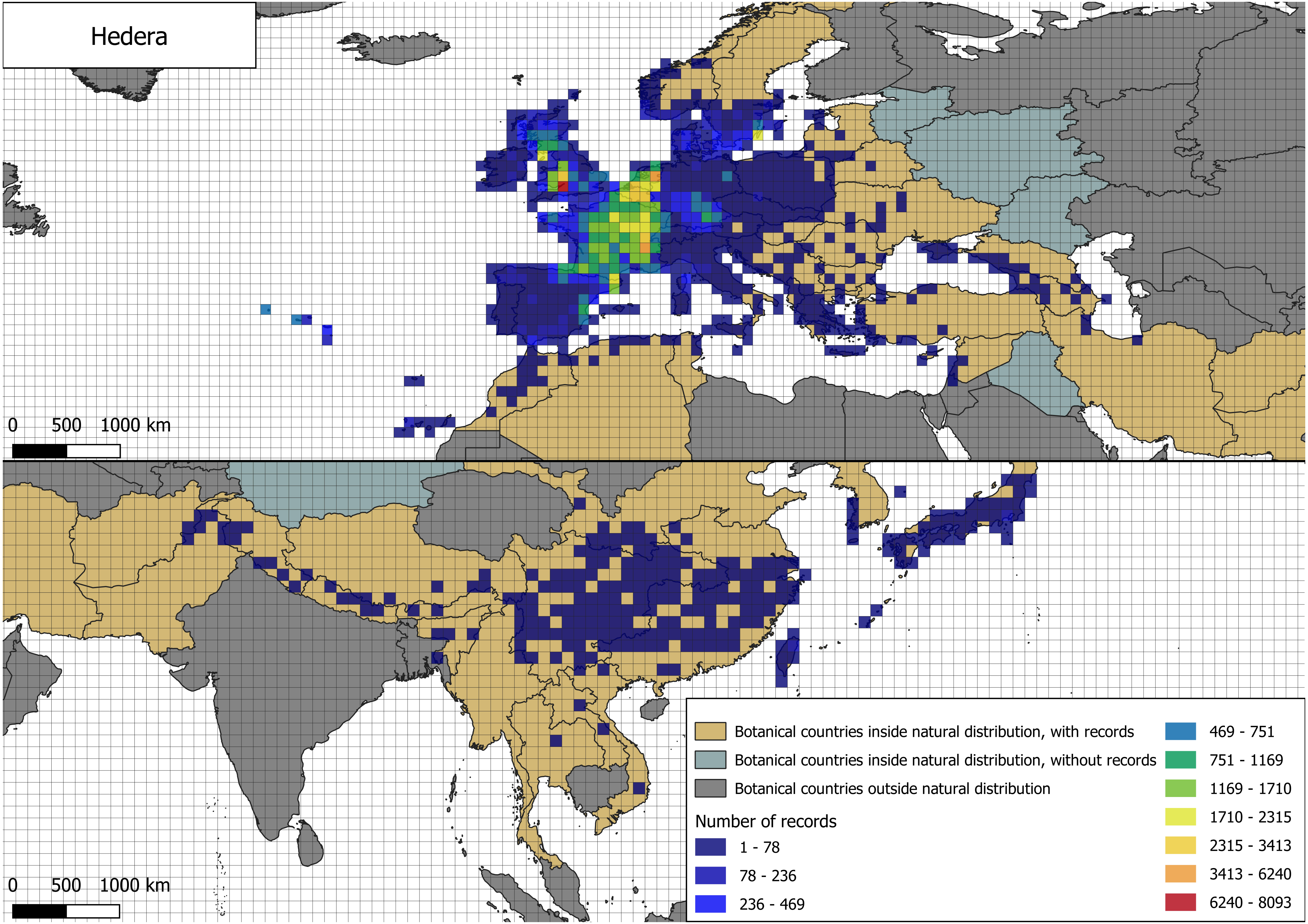
Hedera
0 500 1000 km
0 500 1000 km
Botanical countries inside natural distribution, with records
Botanical countries inside natural distribution, without records
Botanical countries outside natural distribution
Number of records
1 - 78
78 - 236
236 - 469
469 - 751
751 - 1169
1169 - 1710
1710 - 2315
2315 - 3413
3413 - 6240
6240 - 8093

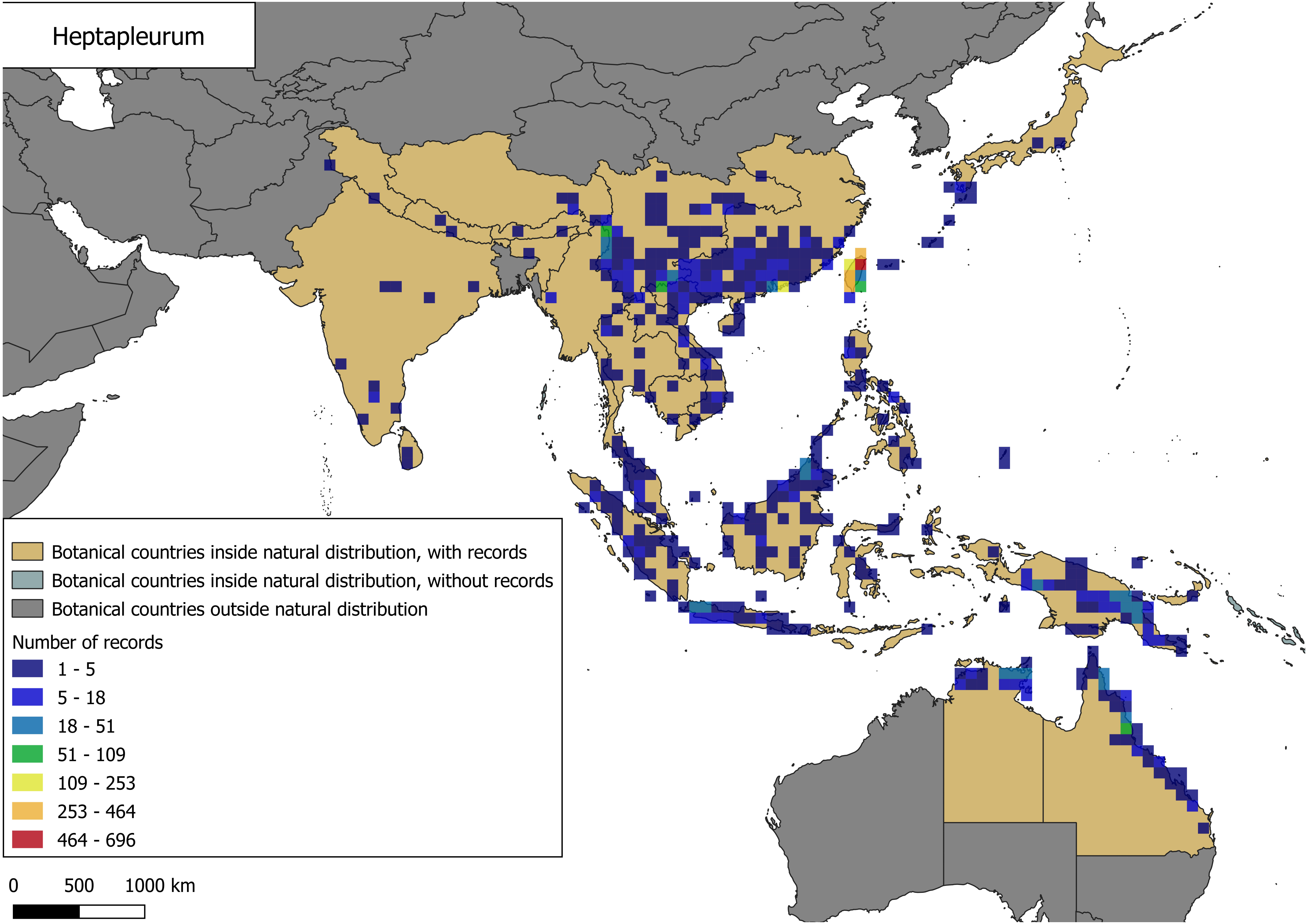
Heptapleurum
Botanical countries inside natural distribution, with records
Botanical countries inside natural distribution, without records
Botanical countries outside natural distribution
Number of records
1 - 5
5 - 18
18 - 51
51 - 109
109 - 253
253 - 464
464 - 696
0
500
1000 km

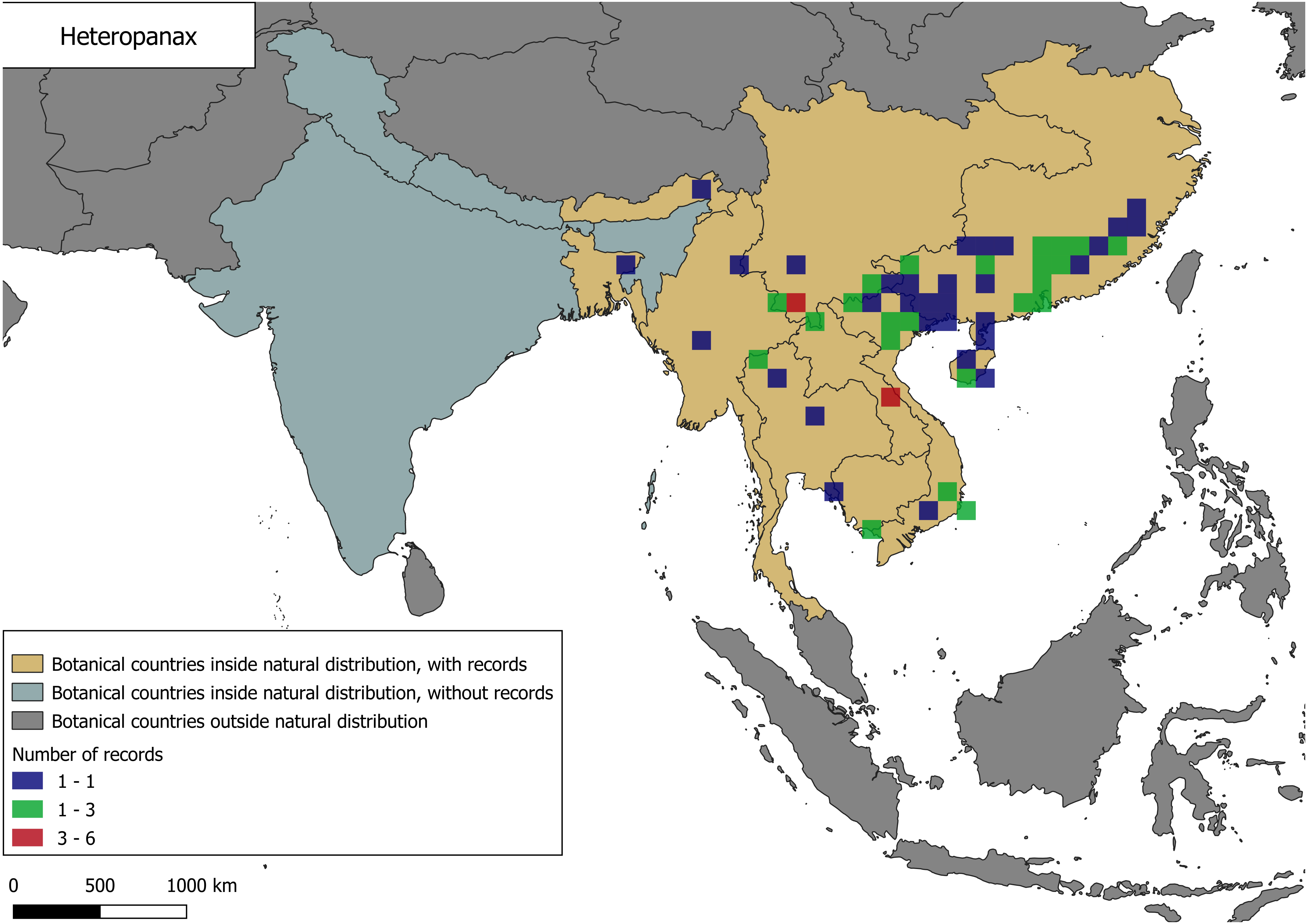
Heteropanax
Botanical countries inside natural distribution, with records
Botanical countries inside natural distribution, without records
Botanical countries outside natural distribution
Number of records
1 - 1
1 - 3
3 - 6
0
500
1000 km

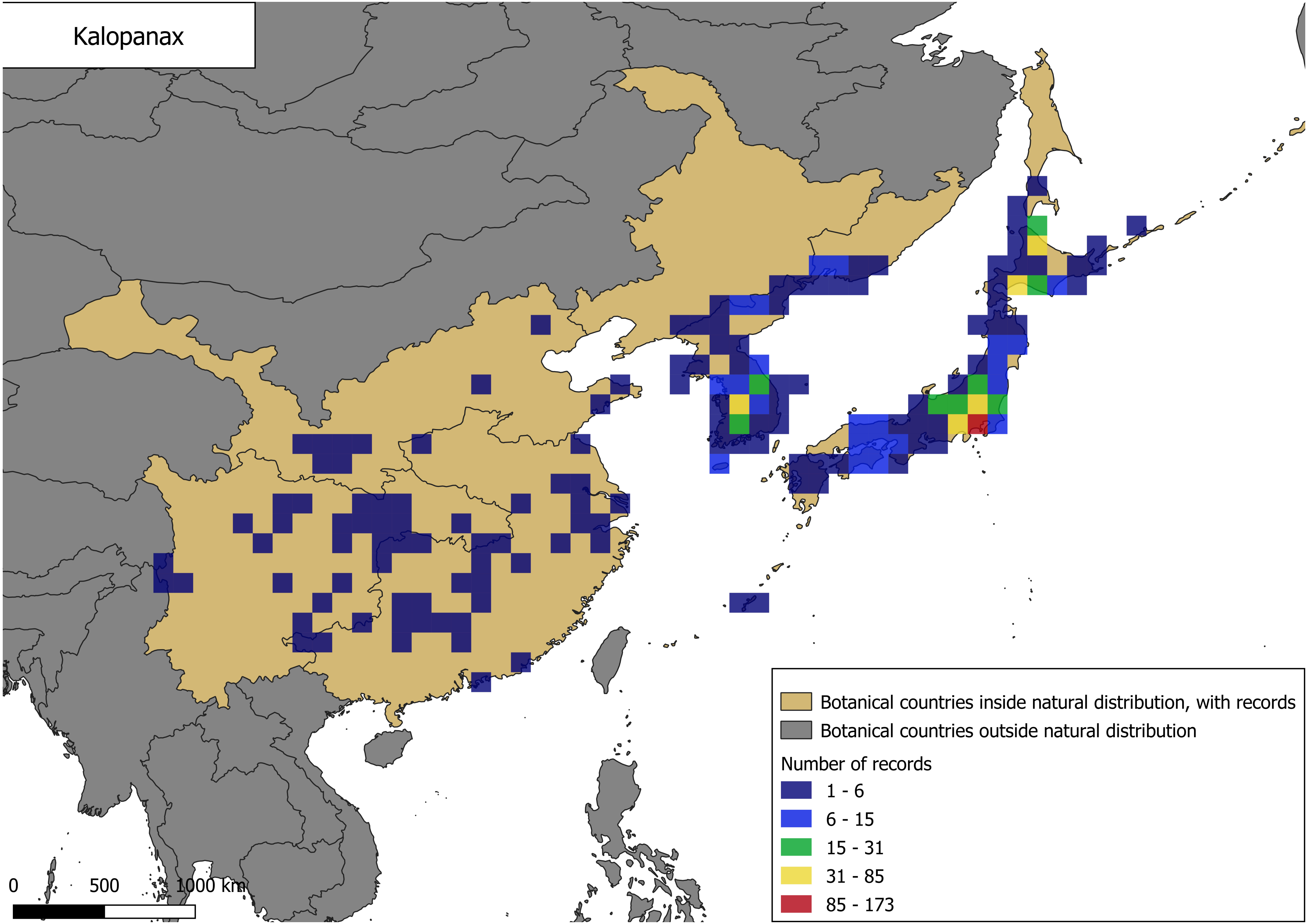
Kalopanax
Botanical countries inside natural distribution, with records
Botanical countries outside natural distribution
Number of records
1 - 6
6 - 15
15 - 31
31 - 85
85 - 173
0
500
1000 km

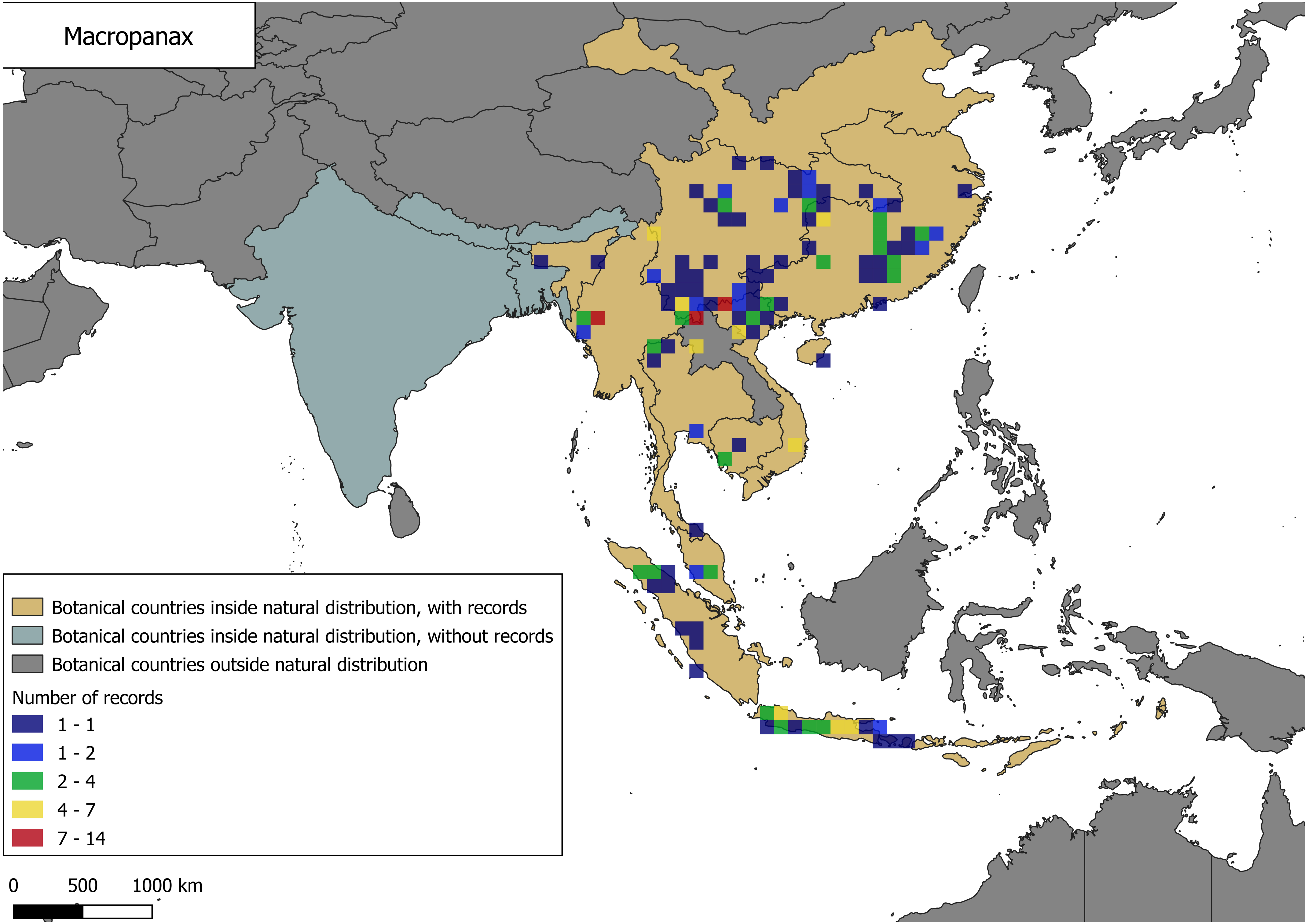
Macropanax
Botanical countries inside natural distribution, with records
Botanical countries inside natural distribution, without records
Botanical countries outside natural distribution
Number of records
1 - 1
1 - 2
2 - 4
4 - 7
7 - 14
0
500
1000 km

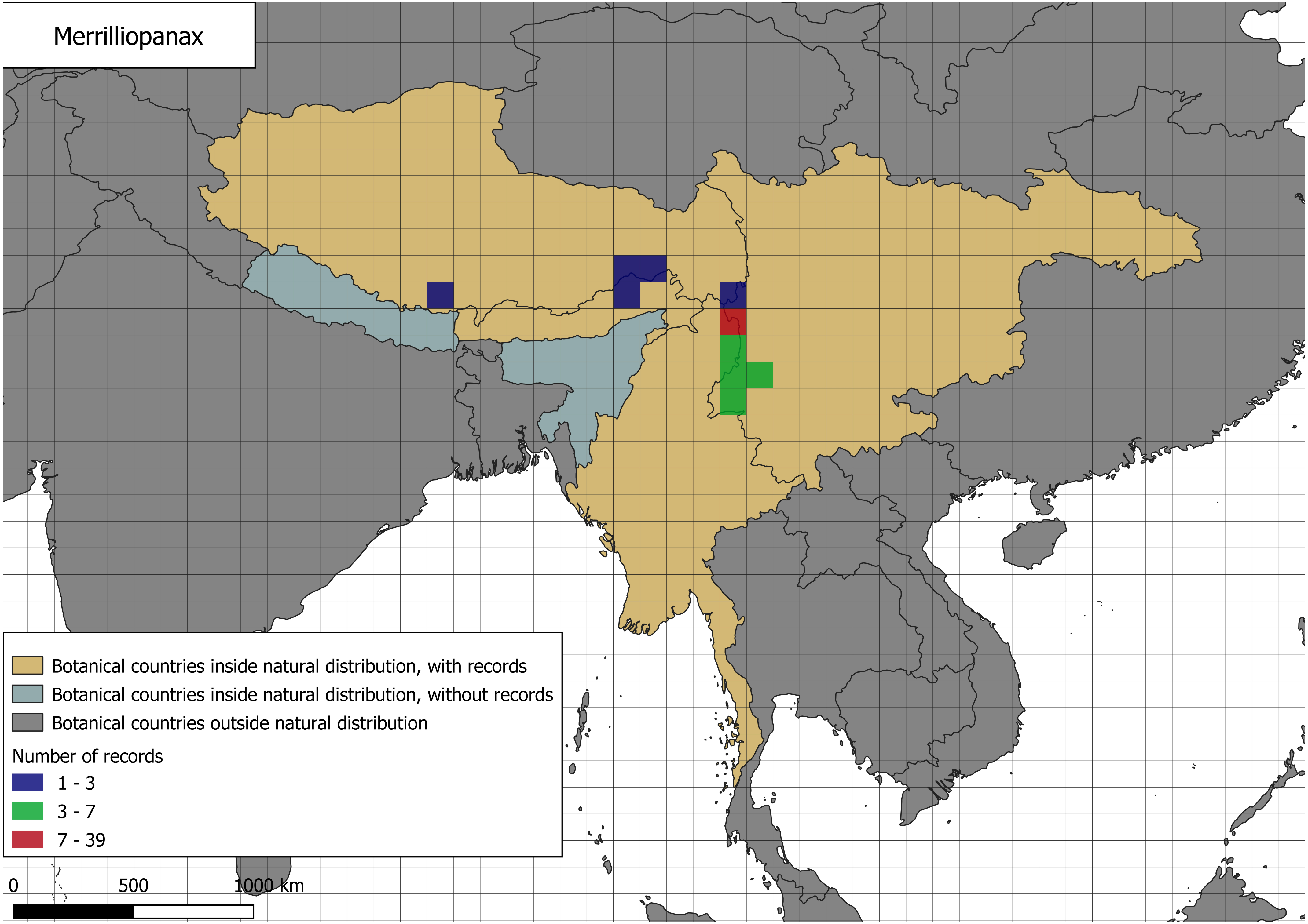
Merrilliopanax
Botanical countries inside natural distribution, with records
Botanical countries inside natural distribution, without records
Botanical countries outside natural distribution
Number of records
1 - 3
3 - 7
7 - 39
0
500
1000 km

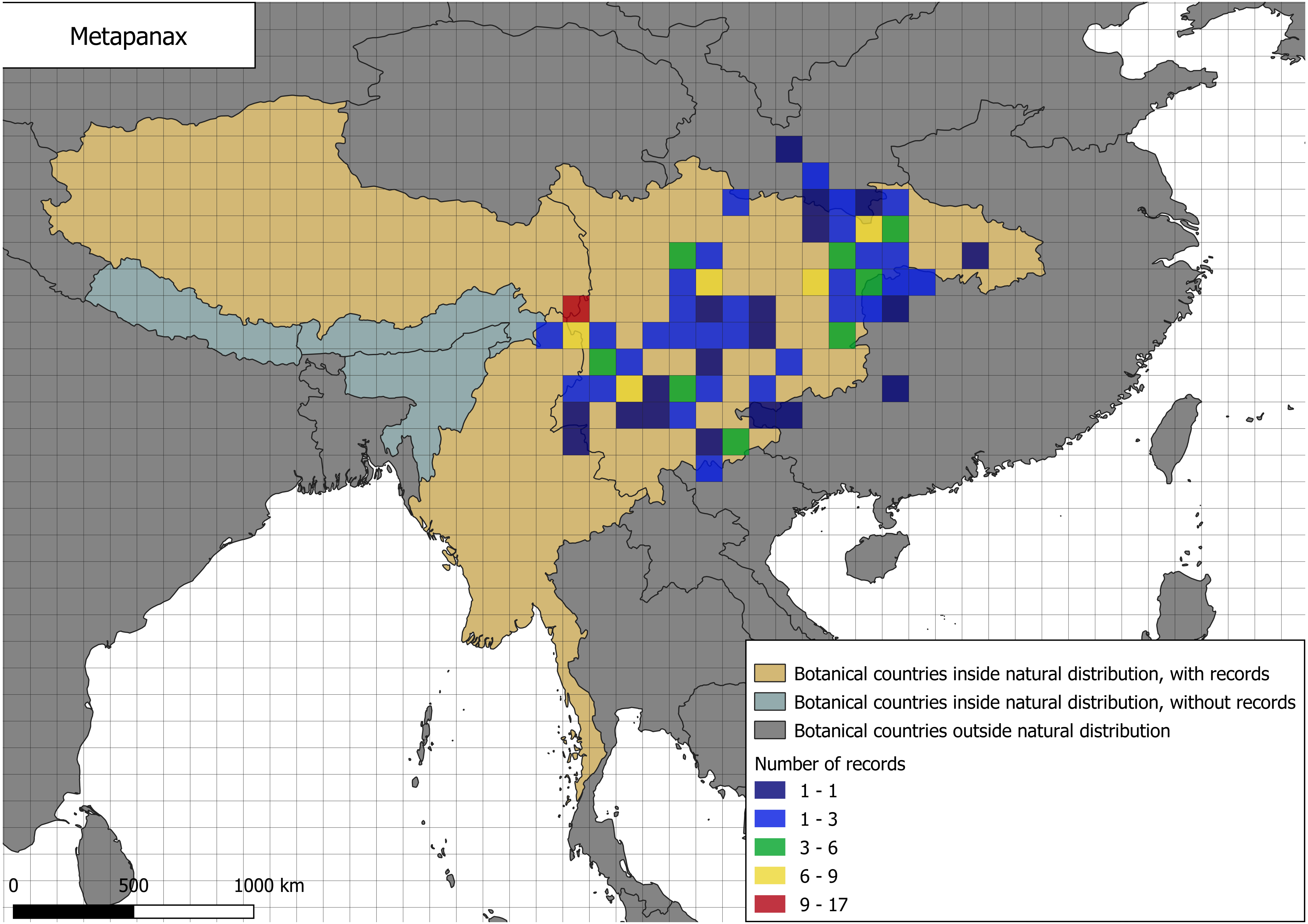

Metapanax
Botanical countries inside natural distribution, with records
Botanical countries inside natural distribution, without records
Botanical countries outside natural distribution
Number of records
1 - 1
1 - 3
3 - 6
6 - 9
9 - 17
0
500
1000 km

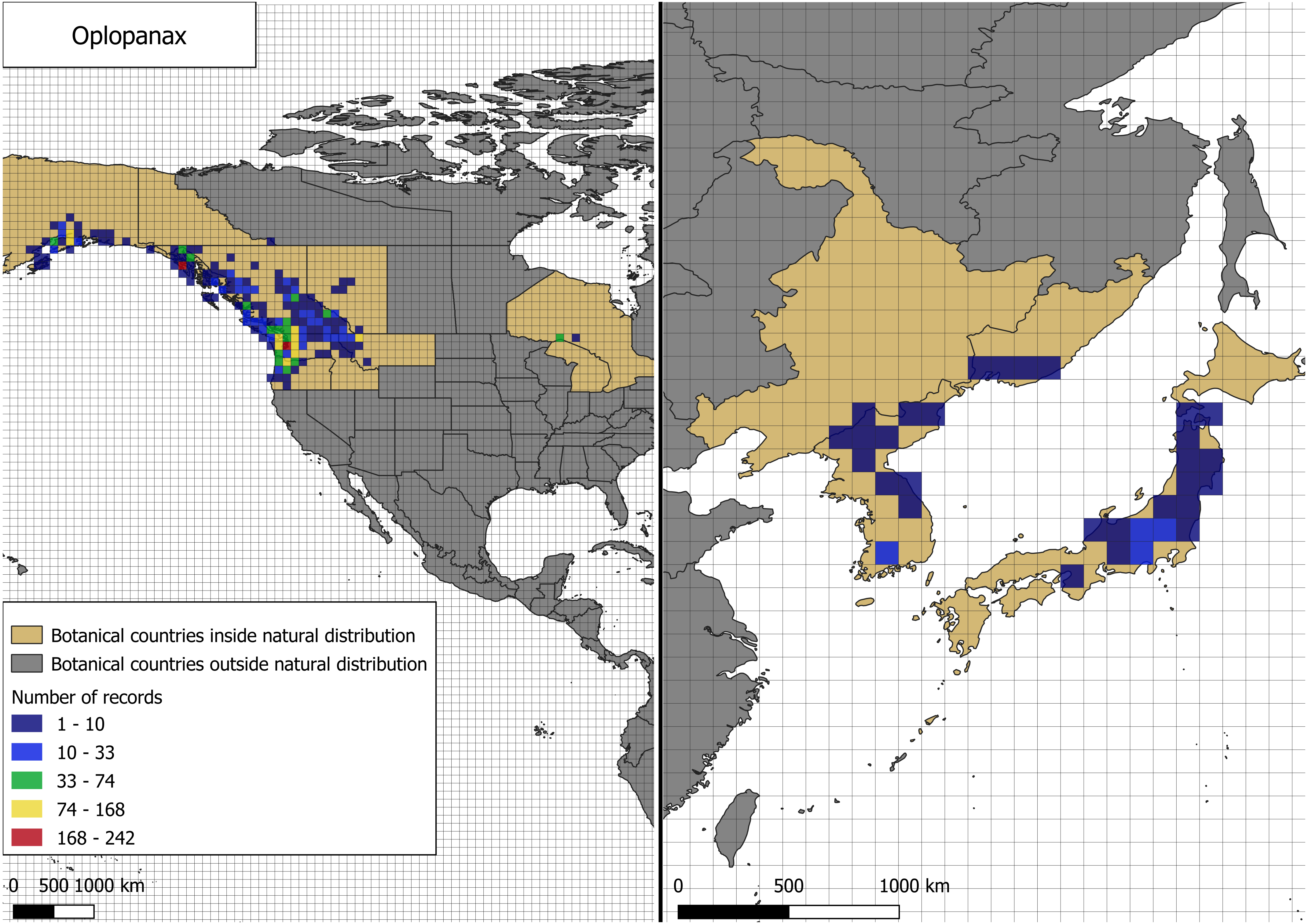
Oplopanax
Botanical countries inside natural distribution
Botanical countries outside natural distribution
Number of records
1 - 10
10 - 33
33 - 74
74 - 168
168 - 242
0 500 1000 km
0 500 1000 km

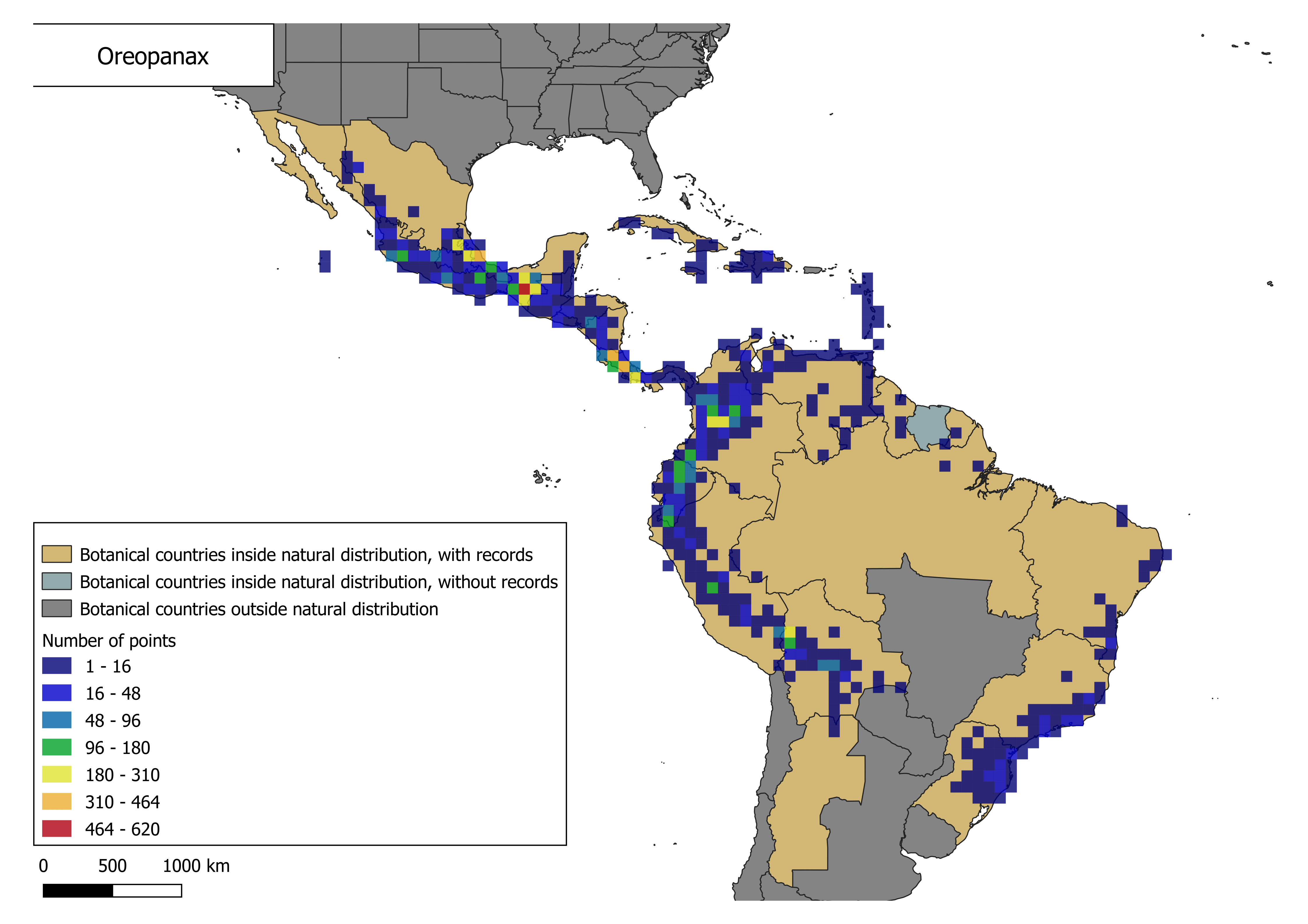
Oreopanax
Botanical countries inside natural distribution, with records
Botanical countries inside natural distribution, without records
Botanical countries outside natural distribution
Number of points
1 - 16
16 - 48
48 - 96
96 - 180
180 - 310
310 - 464
464 - 620
0
500
1000 km

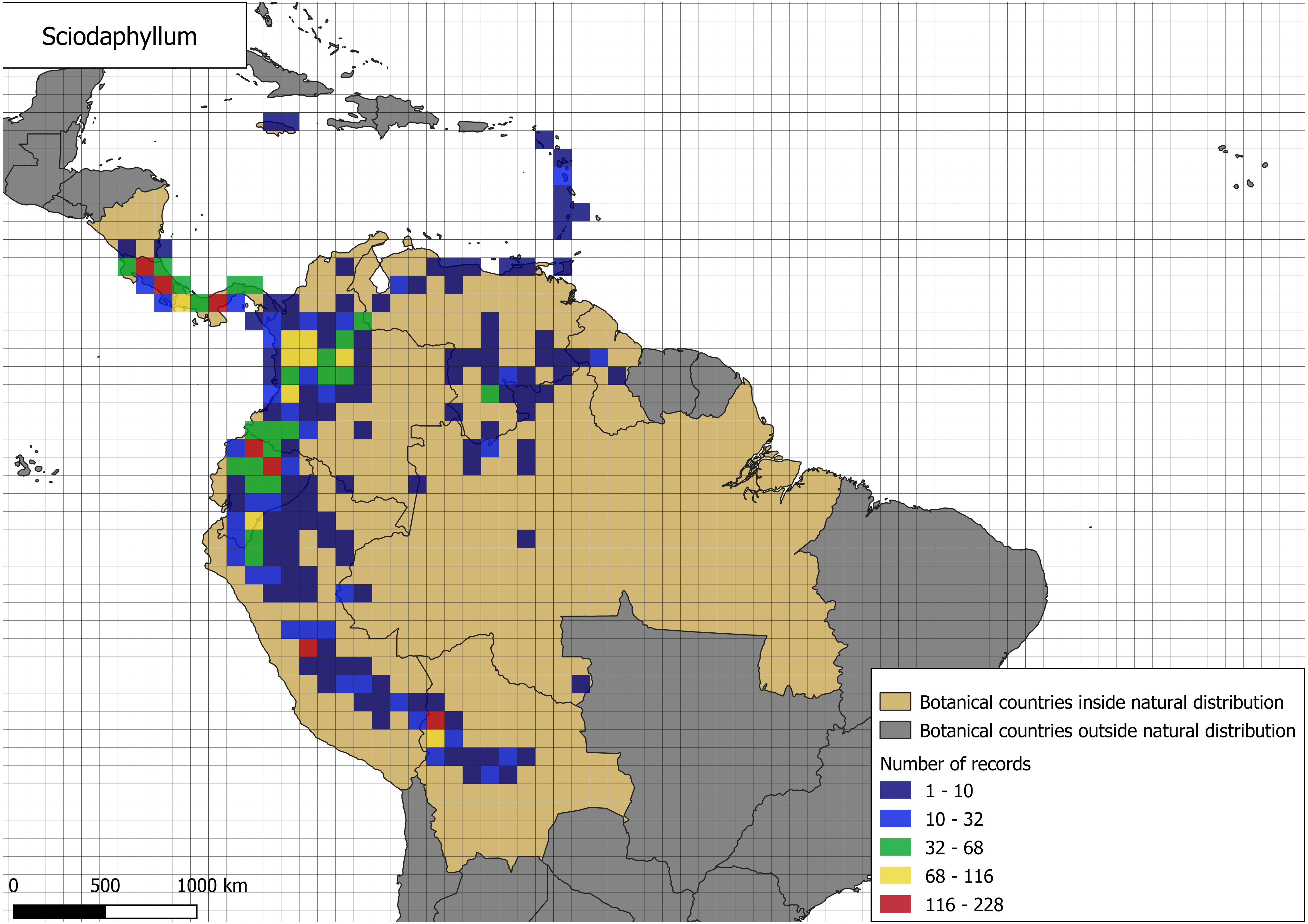
Sciodaphyllum
Botanical countries inside natural distribution
Botanical countries outside natural distribution
Number of records
1 - 10
10 - 32
32 - 68
68 - 116
116 - 228
0
500
1000 km

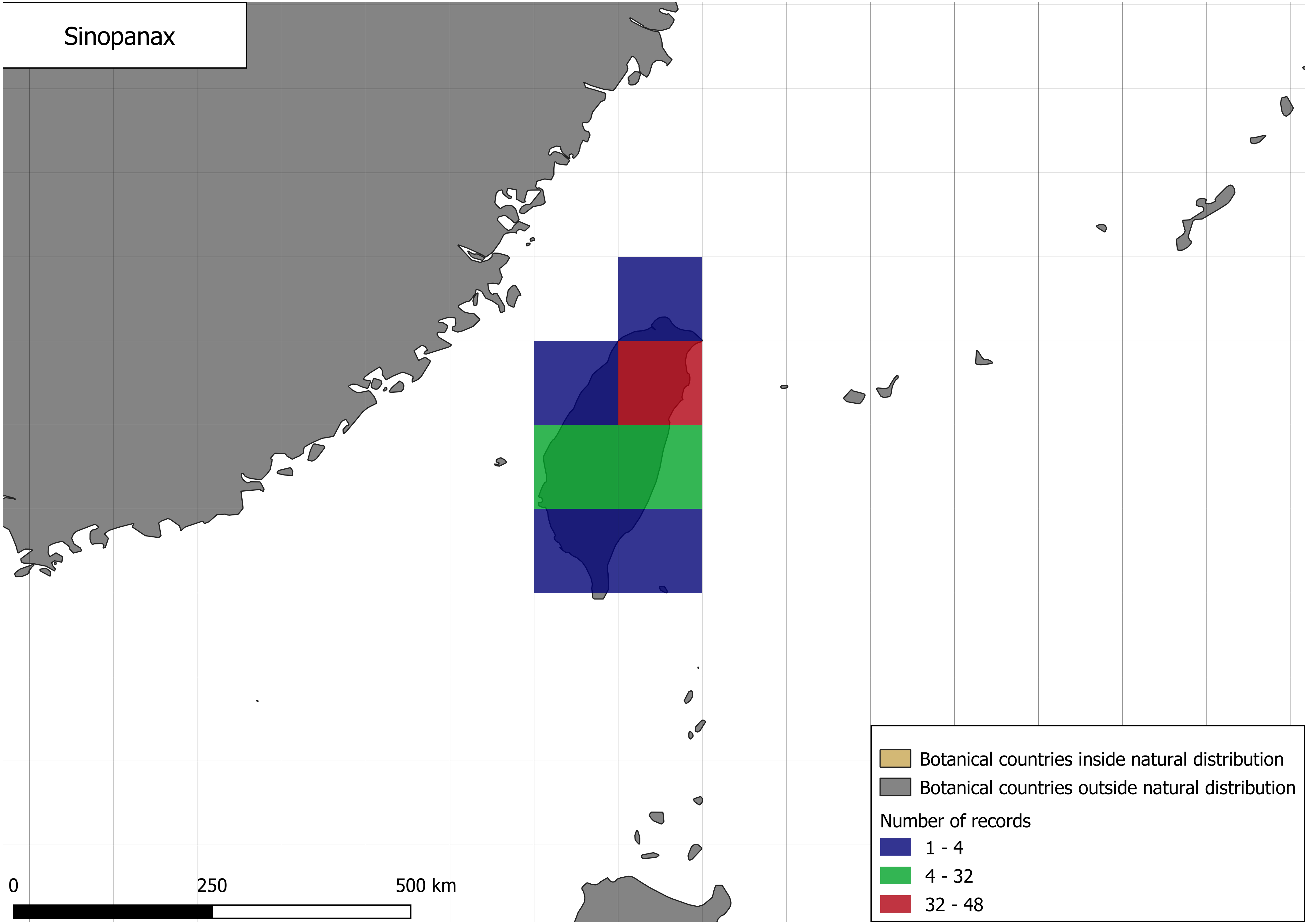
Sinopanax
Botanical countries inside natural distribution
Botanical countries outside natural distribution
Number of records
1 - 4
4 - 32
32 - 48
0
250
500 km

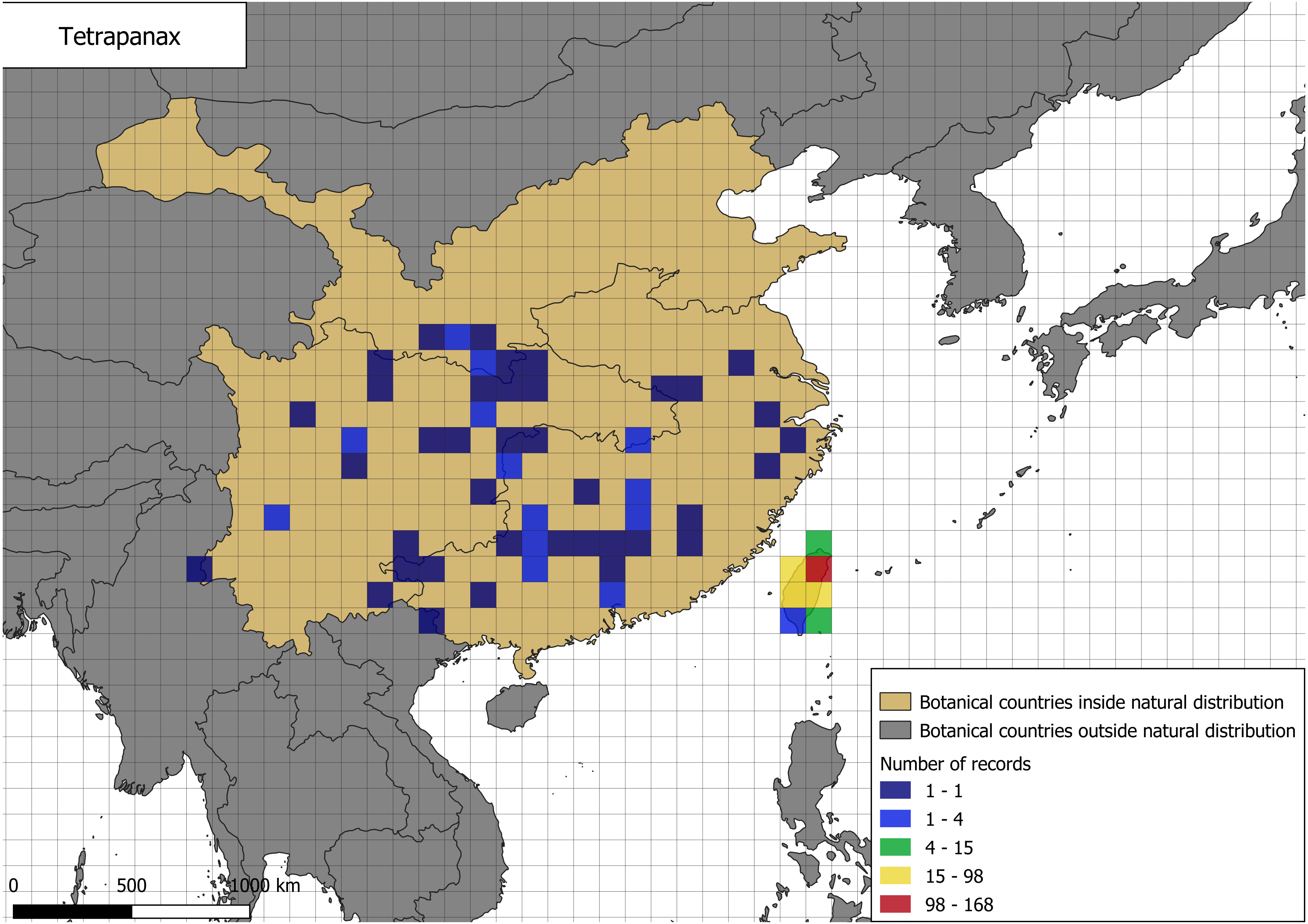
Tetrapanax
Botanical countries inside natural distribution
Botanical countries outside natural distribution
Number of records
1 - 1
1 - 4
4 - 15
15 - 98
98 - 168
0
500
1000 km

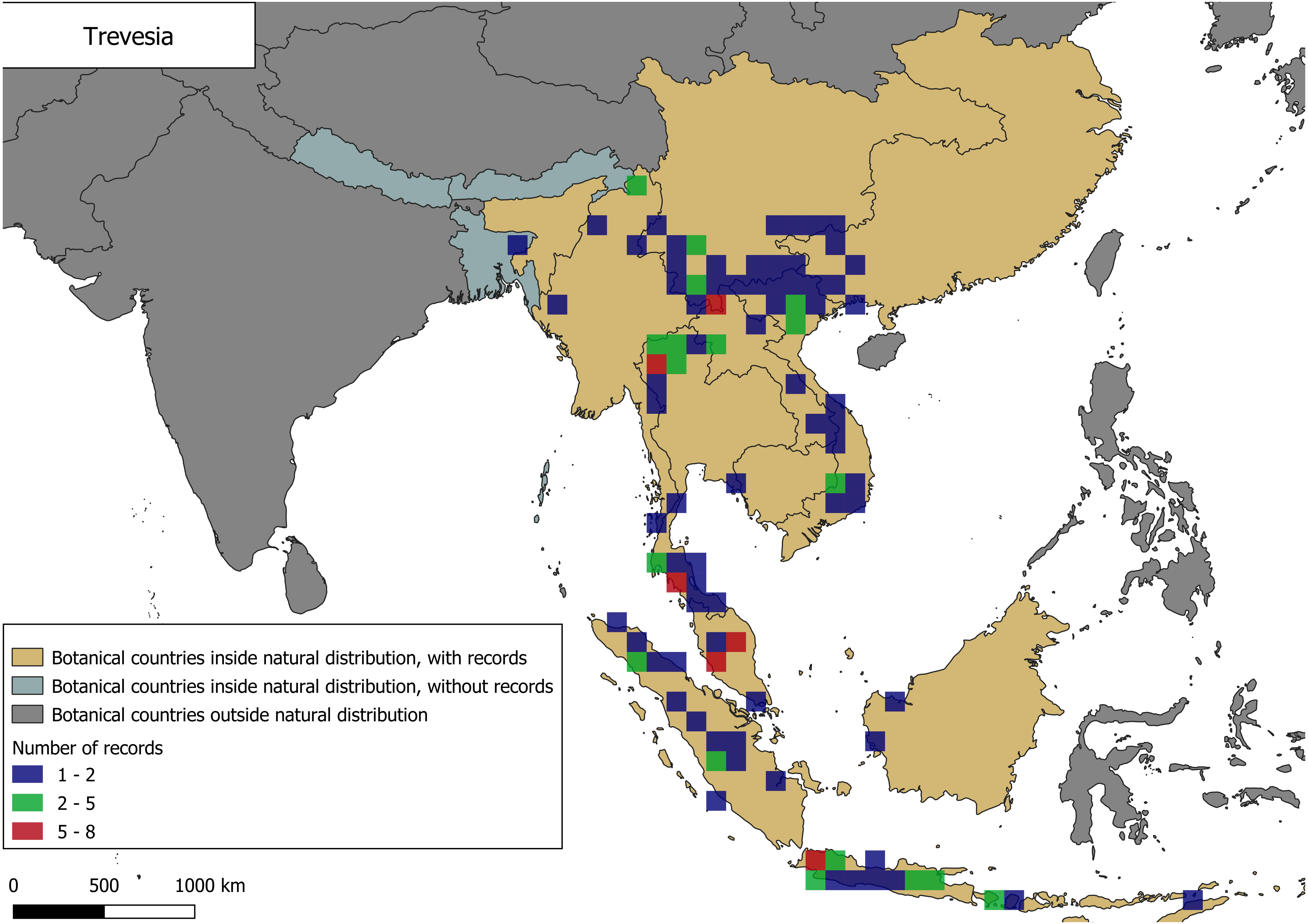
Trevesia
Botanical countries inside natural distribution, with records
Botanical countries inside natural distribution, without records
Botanical countries outside natural distribution
Number of records
1 - 2
2 - 5
5 - 8
0
500
1000 km
